# A Framework for NGN1-Induced Sensory Neuron Differentiation for Disease Modelling and Drug Screening

**DOI:** 10.64898/2026.09.08.750023

**Authors:** Anika Neureiter, Marlene Menke, Veronica Donde, Cedric Günter, Lennart Müller, Florian Kraft, Maike F. Dohrn, Noortje W. M. van den Braak, Andelain Erickson, Lars Buschmann, Anil Kumar Kalia, Susanne Schmitz, Roman Rolke, Ingo Kurth, Martin Zenke, Angelika Lampert

**Author notes:** Corresponding Author: Correspondence and requests for materials should be addressed to Angelika Lampert, Institute of Neurophysiology, Uniklinik RWTH Aachen, Pauwelsstr. 30, 52074 Aachen, Germany.

## Abstract

**Background:** Neuropathic pain is a burdensome, difficult-to-treat, and highly heterogeneous condition with limited therapeutic options, underscoring the need for robust and reproducible human disease models. Human induced pluripotent stem cell (iPSC)–derived sensory neurons provide a promising platform for patient-specific disease modelling and drug screening; however, their translational use is hampered by variability in differentiation efficiency, cellular composition, and functional maturation across protocols and cell lines.

**Methods:** Here, we present a standardized and potentially scalable framework for NGN1-driven differentiation of human iPSCs into sensory neurons. Building on a previously published two-step protocol (1), we systematically deconstructed and optimized each stage of differentiation across a large panel of genetically diverse iPSC lines.

**Results:** We identified robust parameters for neural crest-like cell (NCLC) generation, established a flow-cytometry–based quality-control strategy for NCLCs, and defined optimal combinations of seeding density and lentiviral multiplicity of infection to maximize sensory-neuron progenitor yield.

To improve culture homogeneity, we compared antimitotic selection strategies and demonstrated that tightly timed Ara-C treatment combined with low progenitor seeding density yields consistently pure sensory neuron cultures. We further evaluated maturation under physiologically relevant glucose conditions and performed a systematic review of media compositions to derive two defined maturation media. Morphological, immunocytochemical, transcriptomic, and electrophysiological analyses revealed that time in culture is a major determinant of maturation, while specific supplements such as prostaglandin E₂ (PGE₂) selectively enhance transcriptional signatures associated with nociceptor identity without substantially altering global network activity. Bulk RNA sequencing demonstrated broad expression of sensory neuron and pain-related markers and gene programs across conditions, with long-term maturation and PGE₂ treatment showing the highest similarity to human dorsal root ganglion reference data. Functional assessment using multi-electrode arrays enabled the detection of donor-specific electrophysiological phenotypes, including reproducible hyperexcitability in small-fiber neuropathy patient-derived lines.

**Conclusions:** This study establishes a modular, reproducible NGN1-based differentiation workflow with integrated quality checkpoints that accommodates iPSC line-to-line variability. The framework provides a practical foundation for translational sensory-neuron research, patient-specific disease modelling, and scalable drug screening applications.

## 1. Background

Neuropathic pain affects about 8 % of the general population worldwide and patients often endure years of ineffective treatments, as current therapeutic options provide only limited relief (2–5). This therapeutic stagnation reflects the substantial heterogeneity of patient populations and the diverse underlying pathomechanisms of neuropathic pain (2). Human induced pluripotent stem cell (iPSC)-based disease models offer a promising strategy to partially overcome these limitations. By reprogramming somatic cells from patients and differentiating them into relevant cell types such as sensory neurons, the patients’ genetic makeup is preserved and patient-specific disease mechanisms can be recapitulated *in vitro* (6–9). Such models hold great potential for high-throughput drug screening and the development of personalized therapeutic approaches tailored to the genetic and pathophysiological characteristics of individual patients.

However, a major bottleneck for the clinical and translational use of iPSC-derived sensory neurons is the limited reproducibility of differentiation protocols (9, 10). In the field of sensory neuron differentiation, laboratories employ a wide variety of protocols that differ in materials, timelines, media compositions, and quality control strategies. Even small variations — such as the timing of progenitor passaging — can shift lineage outcomes toward mechanoreceptors or nociceptors (11). Additional factors, including iPSC passage number, epigenetic memory, and clone-specific differentiation biases, further contribute to variability (12–15). As a result, even when the same protocol is applied to multiple cell lines in one laboratory, the resulting sensory neuron cultures can differ markedly in purity, gene expression profiles, and functional properties (16). This variability hampers inter-laboratory comparability and limits the transferability and reproducibility of findings.

To enable the use of iPSC-derived sensory neurons in translational neuropathic pain research in the future, robust and efficient differentiation protocols are needed that function reliably across diverse genetic backgrounds and generate well-characterized neuronal populations with defined functional properties. For disease modelling, iPSC-derived sensory neurons should resemble human dorsal root ganglion (DRG) neurons as closely as possible, including the acquisition of nociceptor identity, expression of key neuronal and pain-related markers, and appropriate morphological and functional characteristics. Moreover, clinically suitable protocols must rely on straightforward steps that can be automated and adapted to xeno-free conditions.

Transcription factor-mediated differentiation has emerged as a powerful strategy for generating neuronal subtypes with high efficiency. Ectopic expression of the basic helix-loop-helix transcription factor NEUROGENIN-2 (NGN2) rapidly drives iPSCs toward a glutamatergic neuronal fate in one- or two-step protocols (17–20). For nociceptor-enriched cultures, NGN1 is particularly relevant: during development, NGN2 drives the first wave of sensory neurogenesis (mechanoreceptors and proprioceptors), whereas NGN1 drives the second wave that gives rise primarily to nociceptors in mice (21). Schrenk-Siemens et al. established an NGN1-based differentiation protocol in which iPSCs are first directed towards a neural crest-like cells (NCLCs) fate, cryopreserved, and subsequently differentiated into nociceptor-like sensory neurons via lentiviral mediated doxycycline-inducible NGN1 expression (1). This approach offers several advantages: (I) iPSCs do not need genetic modification, enabling rapid processing of multiple patient lines soon after reprogramming; (II) NCLC formation accommodates cell-line-specific developmental timing and efficiency; (III) cryopreservation provides a convenient checkpoint for quality control; and (IV) the workflow requires only standard iPSC culture equipment.

The aim of this study was to standardize and optimize the NGN1-mediated sensory neuron differentiation protocol across multiple patient-derived iPSC lines to enable medium to high throughput neuropathic pain disease modelling and drug screenings in the future. To achieve this, we deconstructed the protocol into defined sub-steps and systematically evaluated critical parameters influencing differentiation success. Multiple iPSC lines from 11 donors with distinct genetic backgrounds were included to intentionally introduce a high degree of variability and to ensure that the protocol is broadly applicable across diverse cell lines. To identify the most suitable sensory neuron maturation medium, we compared the formulation of sensory neuron maturation media from 35 publications and created two media that contain widely used components (**SI Table 1**). These media were tested for their suitability to model neuropathic pain on multi electrode arrays (MEA) by differentiating cells from a patient with small fibre neuropathy (SFN), one patient with inherited erythromelalgia (IEM), and one healthy control. To further improve the quality of the resulting sensory neurons, we applied different maturation strategies and benchmarked morphological and transcriptional features against human DRG reference data. Finally, as a proof of concept for scalability and functional relevance, we performed MEA recordings in a 48-well format across five independent cell lines and multiple maturation conditions. We defined functional relevance based on the ability to resolve genotype-dependent differences in spontaneous activity. Together, this work establishes a standardized and scalable framework that can be further adapted for translational disease modelling and drug screening applications.

## 2. Methods

### 2.1 Cell Lines Used in the Study

iPSC lines used in this study are listed in SI Table 2. Following reprogramming, all iPSC lines underwent standard quality control procedures, including assessment of pluripotency marker expression, evaluation of colony morphology, and routine testing for mycoplasma contamination.

### 2.2 iPSC Culture

Pluripotent stem cells were maintained either on truncated human recombinant vitronectin (VTN-N, Life Technologies) or on stem-cell-qualified Geltrex (growth factor reduced, Gibco). Cultures were supplied with StemMACS™ iPS-Brew XF (Miltenyi Biotec). Coating and medium preparation were performed according to the manufacturers’ instructions, and the medium was refreshed daily. Cells were passaged when cultures reached approximately 70 % confluence or when compact colony centres became apparent. For routine passaging, cells were detached using 0.5 mM EDTA (Gibco) in PBS (Gibco) and replated at a split ratio of 1:6 to 1:10. Cultures were maintained in a humidified incubator at 37 °C with 5 % CO₂. For cryopreservation, cells were detached with 0.5 mM EDTA and frozen in a medium containing 90 % iPSC culture medium, 10 % DMSO, and 10 µM Y-27632. Freezing was performed overnight at −80 °C using a Mr. Frosty™ freezing container, followed by long-term storage in the vapor phase of liquid nitrogen.

### 2.3 Production of Lentiviral Particles

Lentiviral particles enabling co-expression of NGN1, eGFP, and puromycin resistance, or expression of rtTA, were generated in HEK293T/17 cells (ATCC CRL-11268) using calcium phosphate–based co-transfection as published previously (1). Cells were transfected with the helper plasmids pRSV-REV, pMD2.G, and pMDLg/pRRE together with either FUW-TetO-Ngn1-P2A-EGFP-T2A-Puro or FUW-rtTA (SI Figure 1) at a ratio of 1:1:2:3. All plasmids were provided by Dr. Katrin Schrenk-Siemens (Institute of Pharmacology, University of Heidelberg). Viral supernatants were harvested 48 and 72 hours after transfection. The NGN1- and rtTA-encoding lentiviral preparations were combined and concentrated by precipitation with 80 µg/ml polybrene and 80 µg/ml chondroitin sulfate (Sigma Aldrich). Viral titers were assessed by infecting 1 × 10⁵ HEK293T cells with defined volumes of the concentrated virus. 24 hours after inducing eGFP expression with 2 µg/ml doxycycline (Sigma Aldrich) in standard culture medium (DMEM (Gibco) supplemented with 10 % FBS (Sigma Aldrich) and 1% sodium pyruvate (Gibco), the proportion of transduced cells was quantified by flow cytometry (BD FACSCanto).

### 2.4 Differentiation of Neural Crest-like Cells and Sensory Neurons

Sensory neuron differentiation driven by ectopic NGN1 expression was carried out using a two-step procedure adapted from Schrenk-Siemens et al. (1), with several modifications. When iPSC cultures reached ∼70 % confluence, cell colonies were lifted as clusters using 0.5 mM EDTA (Gibco) in PBS (Gibco) and transferred to uncoated, tissue-culture-treated 10 cm dishes (Greiner #664160). Aggregates were maintained in Sphere Medium (50 % DMEM/F12 + GlutaMAX, 50 % Neurobasal, supplemented with 0.5 x B27, 0.5 x N2, 0.5× GlutaMAX, 2.5 µg/ml insulin, 10 U/ml penicillin, 1 µg/ml streptomycin) containing 10 ng/ml human EGF, 10 ng/ml human basic FGF (Peprotech), and 10 µM Y-27632 (Stem Cell Technologies). Medium was replaced every other day, omitting Y-27632 after the initial plating. Once spheres adhered and NCLCs migrated outward, spheres were removed manually by aspiration. NCLCs were dissociated using Accutase (Sigma-Aldrich), counted using a CellDrop (DeNovix) with Trypan Blue exclusion, and cryopreserved in Sphere Medium containing 10 % DMSO until further use. For sensory neuron differentiation (step 2), thawed NCLCs were plated onto culture dishes coated with poly-ornithine (Sigma Aldrich), bovine fibronectin (Gibco), and laminin (10 µg/ml each, Sigma Aldrich), LN521 (10 µg/ml, BioLamina), or Geltrex (1:100, hESC-qualified, Gibco) at a density of 0.5-1.5 × 10⁵ cells/cm² in Sphere Medium. After 24–48 hours, cells were co-transduced with lentiviral vectors encoding *NGN1-P2A-eGFP-T2A-PuroR* and *rtTA* at a multiplicity of infection (MOI) of 0.2–4, calculated based on the number of cells seeded after thawing. Transduction medium consisted of Sphere Medium containing 10 mM HEPES and 8 µg/ml protamine sulfate (Sigma Aldrich). The following day, cultures were switched to neuronal induction medium consisting of Sphere Medium supplemented with 20 ng/ml human BDNF, 20 ng/ml human GDNF, 20 ng/ml β-NGF (Peprotech), and 2 µg/ml doxycycline (Sigma Aldrich) to activate transgene expression. Doxycycline was supplied from day (d) 0 to d10 of differentiation, and 75 % of the medium was refreshed daily during this period. On day 5, cells were dissociated with Accutase and replated onto coverslips at 50,000 cells/cm², or onto 48-Cytoview MEA Plates with 15,000-60,000 cells/well. All culture ware, incl. MEA plates, were precoated with poly-ornithine (50 µg/ml) followed by 10 µg/ml laminin/fibronectin, LN521 (10 µg/ml, BioLamina) or Geltrex (1:100) over night at 37°C. After replating, cultures were maintained either in Sphere Medium supplemented with 20 ng/ml BDNF, GDNF, and NGF or transitioned to core medium or physiological medium SI Table 1. 50 % medium changes were performed until d11. Between d7 and d13, cultures were treated with 2 µM cytarabine (Ara-C). Ara-C was diluted 2× in sensory neuron medium, and 50 % of the culture medium was replaced. After 16–20 hours, a complete medium change was performed. Subsequently, half-medium changes without Ara-C were carried out every 2 - 3 days or 3-4 days until the end of the differentiation period.

### 2.5 Culture of the Mouse Schwann Cell Line MSC-80 and MSC-80 - Conditioned Medium

MSC-80 cells were cultured on tissue-culture-treated dishes in DMEM (Gibco) supplemented with 10 % FBS (Sigma-Aldrich), 1 % penicillin/streptomycin, and 1 % sodium pyruvate (Gibco). Cells were passaged at 70–80 % confluence using 0.05 % trypsin. Conditioned medium (cond.med.) was generated when MSC-80 cultures reached approximately 50 % confluence. At this point, the culture medium was replaced with core medium and supernatants were collected every second day for one week. Collected medium was filtered through a 0.2 µm filter and stored at −80 °C until use. For maturation experiments, MSC-80 conditioned medium was diluted 1:2 in core medium. Co-culture experiments were initiated on d 21 of sensory neuron differentiation. MSC-80 cells were seeded directly onto premature sensory neurons at a density of 5,000 cells per MEA well or 15,000 cells per coverslip. Co-cultures were maintained according to the sensory neuron maturation protocol.

### 2.6 Immunofluorescence Stainings

For immunofluorescence staining, cells were cultured and differentiated on glass coverslips pre-treated with 1 M HCl, sterilized for 30 min with 70% EtOH and 1 h UV-light and subsequently coated overnight with 50 µg/ml poly-L-ornithine followed by either 10 µg/ml laminin and fibronectin, 10 µg/ml LN521 (BioLamina) or Geltrex (Gibco, 1:100). At the experimental endpoint, cells were fixed in ice-cold PFA (Cell Signaling Technologies or Sigma Aldrich) for 15 minutes, permeabilized for 10 minutes with 0.1 % Triton X-100 (Sigma Aldrich) in PBS, and blocked for 1 hour in 5 % normal goat serum (Pan Biotech) or bovine serum albumin (Sigma Aldrich)e in PBS (blocking buffer). Primary antibodies were diluted in blocking buffer according to SI Table 3 and incubated overnight at 4 °C. The following day, samples were washed three times for 5–10 minutes with PBS and incubated with fluorophore-conjugated secondary antibody diluted in blocking buffer for 1–2 hours at room temperature, protected from light. After three additional PBS washes, DAPI (NucBlue™ Fixed Cell ReadyProbes™ Reagenz (DAPI)) or Hoechst 33342 (NucBlue™ Live ReadyProbes™ Reagenz (Hoechst 33342)) was added during the second wash step. Finally, cells were mounted on glass slides using ProLong™ Glass Antifade Mountant (Life Technologies) and imaged with an EVOS M5000 microscope (Life Technologies) or Zeiss LSM 700 attached to an Axio Observer Z1.

### 2.7 Calcium Imaging

Calcium imaging was performed using a Hamamatsu ORCA-Flash 4.0 LT Plus camera with 340/380 nm illumination provided by a CoolLED pE-340fura system, reflected through a T400lp dichroic mirror and collected through an ET510/80m emission filter (Chroma Technology). Recordings were acquired using a Fluar 20× objective (Zeiss, Cat# 420150-9900-000) mounted on a Zeiss Axiovert 10 microscope for all time points, except day for 32 experiments, which was acquired on a Zeiss Axio Vert.A1. Image acquisition was controlled with μManager v2.0 (Edelstein et al., 2014) at 1 Hz with 50 ms exposure time. Frames were acquired at 10-second intervals during baseline and compound application, and at 20-second intervals during washout periods. For dye loading, coverslips containing NGN1-differentiated neurons were incubated with freshly prepared Fura-2 AM (Life Technologies, Cat# F1225) diluted to 2 µM in bath solution for 15–20 minutes at room temperature in the dark. The bath solution contained (in mM): 140 NaCl, 3 KCl, 1 MgCl₂, 1 CaCl₂, 10 HEPES, and 20 glucose (pH 7.4). After loading, coverslips were transferred to an RC-25 Open Diamond Bath Imaging Chamber (Warner Instruments) mounted in a PH-3 chamber and perfused continuously with bath solution for 20 –30 minutes before imaging. Illumination intensity at 340 and 380 nm and camera thresholds were adjusted to optimize signal-to-noise ratio while minimizing photobleaching. Spontaneous activity was recorded for 15 minutes under constant perfusion, followed by sequential application of menthol (500 µM, 2 min), capsaicin (1 µM, 2 min), and veratridine (50 µM), each separated by 5 - minute washout periods. All recordings were performed at room temperature (22-24 °C). Veratridine (Alomone Labs, Cat# V-110), menthol (Sigma-Aldrich, Cat# M27720), and capsaicin (Sigma-Aldrich, Cat# M2028) were dissolved at 1000× concentration in 70 % ethanol, stored at −20 °C, and diluted in bath solution immediately before use. Fura-2 AM aliquots were stored at −20 °C, and a fresh working aliquot was thawed for each experimental day. Intracellular calcium signals were extracted and analysed as follows: Mean grey value was exported for manually drawn regions of interest (ROIs) in ImageJ (22) covering the soma of cells to be analyzed, and five regions devoid of cells and processes for an average background subtraction per image. Mean grey values for background-subtracted ROIs recorded in the 340 nm channel were divided by those recorded in the 380 nm channel to obtain the ratiometric value (R = F_340_/F_380_) in Microsoft Excel. F_340_/F_380_ during the first 4 minutes of recording were averaged (R_0_), and all values were normalized to this baseline (R/R_0_). Baseline-normalized values were exported to GraphPad prism 9.0 (San Diego, CA) to visualize calcium transients and compute means. A transient increase of ≥20% from baseline (R/R_0_ ≥ 1.2) during compound application was categorized as a responding cell, or as a spontaneously active cell if occurring during the 15 minutes of baseline recording.

### 2.8 Flow Cytometry (FACS)

NCLCs and iPSCs were cultured and differentiated as described before. Spheres that remained in suspension were transferred onto coated coverslips and harvested immediately after attachment. All cells were dissociated with Accutase for 5 min to obtain a single-cell suspension. Following centrifugation, cells were resuspended in FACS buffer and counted using a CellDrop (DeNovix), including viability assessment with Trypan Blue (Gibco). For staining, 1 × 10⁵ cells were incubated in 100 µl FACS buffer containing the respective antibodies (SI Table 3) for 30–60 minutes at 4 °C. After staining, cells were washed three times with PBS and resuspended in fresh FACS buffer for acquisition. Flow-cytometric measurements were performed on a BD FACSCanto, and data were analyzed using FCS Express 7 (De Novo Software). Cell populations were first gated based on forward and side scatter characteristics. Negative and positive populations were defined using isotype controls, applying a background fluorescence threshold of 0.5 %. All plots shown in the manuscript were exported directly from the analysis software.

### 2.9 Multi-Electrode Array (MEA) Recordings

MEA Recordings were performed with an Axion Maestro 1 (Axion Biosystems). 48-well CytoView MEA plates (Axion Biosystems) were pre-treated with 1 % Tergazyme (Merck Millipore) freshly diluted in water for 2 hours, followed by three washes with sterile water. For sterilization, wells were incubated with 70 % ethanol for 30 minutes and air-dried under sterile conditions. Plates were then filled with DMEM (Gibco) and equilibrated at 37 °C, 5 % CO₂, and 100 % humidity for several days before coating. For coating, plates were washed three times with sterile water and incubated overnight at 37 °C with 50 µl of 50 µg/ml poly-L-ornithine (Sigma-Aldrich). The next day, plates were washed again and coated with either 10 µg/ml laminin (Sigma Aldrich)+ 10 µg/ml bovine fibronectin (Gibco) diluted in Neurobasal medium, or 10 µg/ml LN521 (BioLamina) diluted in PBS containing MgCl₂ and CaCl₂ (Gibco). Coating was performed overnight at 37 °C. Sensory neuron progenitors were routinely passaged on day 5 of differentiation. Cell suspensions were adjusted to the desired concentration in 50 µl and seeded directly onto the MEA electrode area. After attachment, wells were filled with 250 µl of complete sensory neuron medium. Subsequent culture followed the NGN1-mediated sensory neuron differentiation protocol. Recordings were performed at defined time points. MEA plates were placed in the Maestro system pre-heated to 37 °C, 5 % CO_2_ and equilibrated for 10 minutes before acquisition. Spontaneous activity was recorded for 15 minutes using AxIS software (Axion Biosystems). For temperature-dependent experiments, recordings were performed sequentially at 37 °C, 25 °C and 42 °C. At each temperature, 15 min-recordings were initiated 10 minutes after the temperature control unit reached the target value. Recordings were performed using the following settings: Configuration = Neural Real-Time, spontaneous and a Butterworth filter with a high pass filter cut-off frequency of 200Hz and low pass filter cut-off frequency of 3000Hz. Spike detector was set to detect only events crossing the 6x standard deviation of noise per electrode. Bursts were detected based on the Inter-Spike-Interval (ISI) Threshold set to a maximum Inter-Spike Interval of 100 ms and a Minimum Number of Spikes of 5. Network Bursts were defined by a maximum ISI of 100 ms, minimum number of spikes of 50 and a minimum of 35 % participating electrodes. 20 ms window size is applied to compute the area under the cross-correlation and area under the normalized cross-correlation synchrony metrics. Active electrodes were defined by a minimum spike rate of 0.083 spikes/min. All raw parameters (spike-based, burst-based, and variability-related features; SI Figure 4) were extracted using the Axis Software (Axion Biosystems). All extracted parameters were annotated and stored in a structured database. Analyses were performed in Python using pandas, numpy, seaborn, matplotlib, sklearn (StandardScaler, PCA, UMAP), scipy, and pathlib. Due to variability across electrodes, maturation experiments were analyzed at the electrode level. Only electrodes that reached >0.01 Hz at any time point were included. For each condition (patient × treatment × day), electrode-level values were averaged to obtain one representative value per patient. Median ± IQR across patients was visualized. Dimensionality reduction (PCA and UMAP) was performed on all activity and burst parameters (excluding network metrics). Treatment medians across patients were used as input. Temporal dynamics were quantified by extracting the time-to-maximum and maximum value for each parameter; medians across patients were plotted for selected conditions. To summarize functional phenotypes, mean MEA parameters per patient were grouped into activity features (Number of Spikes, Mean Firing Rate (Hz) and ISI Coefficient of Variation) and burst features (Number of Bursts, Burst Duration - Avg (s), Number of Spikes per Burst – Avg, Mean ISI within Burst - Avg, Median ISI within Burst – Avg, Inter-Burst Interval - Avg (s), IBI Coefficient of Variation, Normalized Duration IQR, Burst Frequency (Hz) and Burst Percentage). Selected features were log-transformed (log₁₀(x + 1)) and standardized using z-score normalization. Composite descriptors were calculated as: Activity vector = mean of all standardized activity features and Burst vector = mean of all standardized burst features. For conditions without detectable bursting, burst vectors were set to zero. These vectors were used to visualize functional states in a two-dimensional activity–burst space. For disease-modeling comparisons, data from defined differentiation stages (day 70 and day 140) were analyzed. Mean values per parameter and patient were plotted. To compare heterogeneous electrophysiological features, values were normalized per feature using min–max scaling. Missing values were retained and visualized as blanks. Hierarchical clustering was performed on normalized data, with missing values imputed using the feature mean.

For statistical analysis of the iPSC derived sensory neuron repsonse to excitatory agents used the Spike (.spk) files produced by the Axis-Software (Axion Biosystems). Data were loaded using the pyaxion package (1.0.2). All analysis was carried out on two-minute windows taken from a baseline period and a post-injection period. In case the baseline and injection period were recorded in a single file, the recording period was split into baseline and injection period based on external metadata about the injection time of the first compound. The two-minute baseline window was taken from the middle of the baseline period, while the post-injection period was aligned to 1 s from the back (to avoid boundary effects) of the file and offset for the average injection time per well, resulting in a staggered pattern that on average has the same offset with respect to the injection time of each well. The spike vector of each electrode was time binned by sliding a one second window overlapping 50 % over the timestamps, counting the spikes per window (no boundary correction). The timeseries of each window was then transformed into a firing frequency histogram using equally spaced bins of 1 Hz between 0 and 50 Hz. Data of all differentiations and wells were pooled for further analysis. Based on the histogram, the Jensen-Shannon Distance (JSD) was calculated as given in equation 1 using a base 2 logarithm in the Kullback-Leibler Divergence calculation, resulting in values between 0 (perfect resemblance of histograms) to 1 (no resemblance at all).

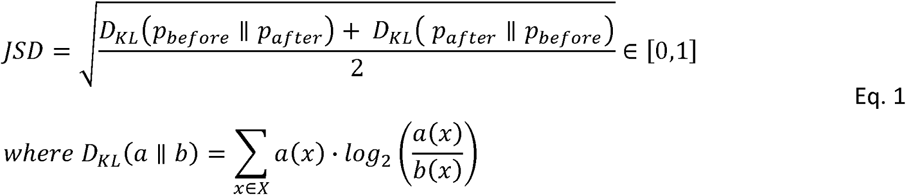

The JSD was used to evaluate whether electrodes react to the compound addition in any way (JSD > 0). To ensure mathematical stability of further analysis, electrodes were filtered on the condition that they had to show at least 5 spikes in the baseline window, equating to a firing rate threshold of 0.042 Hz. Additionally, z-score outlier removal was performed using a threshold of 3. To further elucidate and statistically assess the change in firing behavior, the relative change in mean firing frequency *Δf̄_rel_* inside the windows was calculated using equation 2. Comparative tests were performed between the compounds and their respective solvents (DMSO ↔ Capsaicin, EtOH ↔ Menthol, AITC, αβATP). Statistical significance was tested using a two-sided Mann-Whitney-U test with Benjamini-Hochberg false discovery rate correction. Significance levels are given by * = 0.05, ** = 0.01, *** = 0.001. All steps were performed using numpy (2.2.5),scipy (1.15.2) and python3 (3.12.11)

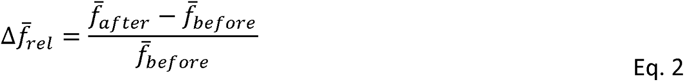

### 2.10 mRNA Expression Levels Assessed by qRT-PCR

Cell pellets were collected, snap-frozen in liquid nitrogen, and stored at −80 °C until processing. Total RNA was isolated using the NucleoSpin RNA Mini Kit (Macherey-Nagel) according to the manufacturer’s instructions. RNA concentration and purity were assessed using a NanoDrop spectrophotometer (Thermo Fisher Scientific). For cDNA synthesis, 1 µg of total RNA was reverse-transcribed using the SensiFAST™ cDNA Synthesis Kit (Meridian Biosciences) with an extended reverse-transcription step of 25 minutes in a PeqStar thermal cycler (PeqLab). Resulting cDNA was diluted in nuclease-free water (Qiagen) to a final concentration of 2 ng/µl. qPCR reactions were performed using the SensiFAST™ SYBR® No-ROX Kit (Meridian Biosciences) in a total volume of 13 µl, containing 10 ng cDNA and 250 nM of each primer SI Table 4. Cycling conditions were as follows: 95 °C for 5 s (denaturation) and 60 °C for 30 s (annealing/extension) for 40 cycles, followed by melt-curve analysis from 60 °C to 95 °C in 1 °C increments. Data were analyzed using CFX Manager (Bio-Rad). Relative gene expression was calculated using the ΔΔCT method with GAPDH and H1C0RF43 as reference genes. For each cell line, values represent the mean ± SD of three technical replicates. Data visualization was performed in Python 3.12.7.

### 2.11 Soma Size Measurements

Soma size of iPSC-derived neurons was determined in phase contrast images taken with the EVOS M5000 Imaging System (Life Technologies) with 10x and 20x magnification. Soma size quantification was performed in ImageJ/Fiji (version 1.54p, National Institutes of Health, USA) running with Java 1.8.0_453 (64-bit). Before analysis, the image scale was calibrated with the set scale function. Soma area was measured by manual definition of regions of interest (ROIs) using the polygon selection to create soma masks. The set measurements function, with Area and Fit Ellipse enabled, was used to obtain measurements. Each ROI with corresponding overlays was saved individually for traceability. For better comparison, the soma diameter was calculated based on the soma area measured. The soma diameter correlated with the soma area based on 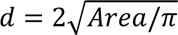 (SI Figure 9C).

### 2.12 Transcriptomics Using RNAseq

After the final MEA recording, cell pellets were snap-frozen in liquid nitrogen and stored at −80 °C. Total RNA was isolated using the NucleoSpin RNA XS Micro Kit (Macherey-Nagel) according to the manufacturer’s instructions. RNA concentration and integrity was verified on a TapeStation (Agilent). Libraries for RNA-Seq were prepared using the NEBNext® Ultra™ II Directional RNA Library Prep Kit together with the NEBNext® rRNA Depletion Kit (human/mouse/rat) according to the manufacturer’s protocol. Sequencing was performed on an Illumina NovaSeq6000 at 2×159bp, generating approximately 15-20 million reads per sample. Raw data were demultiplexed and FASTQ files were generated using bcl_convert. Data were aligned to the GRCh38p14 genome with STAR aligner. Bulk RNA sequencing data were further processed using custom Python scripts. Raw feature-level count matrices were imported using the pandas package. Samples with sequencing depth below 2 million reads were excluded from downstream analyses. Counts were normalized using counts per million (CPM) by dividing raw counts by the total library size per sample and multiplying by 1 × 10⁶. Normalized values were log-transformed as log₂(CPM + 1) for downstream analyses and visualization. Expression of selected marker genes representing pan-neuronal, sensory neuron, mechanosensory, proprioceptor and nociceptor identities was visualized. Marker genes were curated based on established literature. To assess global transcriptional differences between treatments and cell lines, a PCA was performed using scikit-learn. Genes with zero variance were removed, and remaining genes were z-score normalized. PCA was computed on the top 1000 most variable genes across all samples. Count matrices were exported and analyzed using DeSeq2 (v. 1.50.2) and edgeR (v. 4.8.2) packages based on experimental design (d 70 of differentiation vs d168, and dbcAMP, retinoic acid or PGE_2_ addition). Gene set enrichment analysis (GSEA) was performed using pre-ranked gene lists derived from differential expression analyses. Gene sets were obtained from Gene Ontology (Biological Process; (GO:BP 2021)). For visualization, normalized enrichment scores (NES) were used to generate heatmaps of pathways related to sensory neuron development and nociceptor function, pain pathways, membrane potential pathways, synapse pathways, neural crest cell pathways, axon and dendrite pathways. To compare iPSC-derived sensory neurons with human dorsal root ganglia (DRG), publicly available single-cell RNA-seq data (23) were used as a reference. A pseudo-bulk approach was applied by summing counts across all cells within each DRG sample. Pseudo-bulk profiles were normalized using CPM and log₂-transformed identically to the iPSC-derived samples. Transcriptomic similarity between iPSC-derived neurons and the human DRG reference was quantified using Pearson correlation coefficients. Analyses were performed using either the top 3000 most variable genes or neuronal gene sets (SI Table 5). Correlation coefficients were transformed using Fisher’s z-transformation prior to statistical testing. Differences between treatment groups were assessed using one-way ANOVA followed by Tukey’s honestly significant difference (HSD) post hoc test. Statistical analyses were performed in Python using scipy and statsmodels.

## 3. Results

### 3.1 Implementation of the NGN1-Driven Differentiation of Sensory Neurons

To identify a protocol suitable for subsequent optimization and standardization, we first evaluated whether the NGN1-based sensory-neuron differentiation protocol (“NOCL3”) described by Schrenk-Siemens et al. (2022) is robustly implemented in our laboratory. Three control iPSC lines from independent donors (B1, Fib7_1, and Fib13_30) were selected for this initial assessment. Expanded iPSCs were induced to form spheres, and within 24 h, round, free-floating aggregates with sharp borders were observed (Figure 1A). The following days, spheres increased in size and began to adhere to the uncoated plastic surface around day 9-14 (Figure 1A). Shortly after attachment, NCLCs migrated outward from the sphere periphery. Attached spheres were removed, and the emerging NCLCs were cryopreserved. Remaining floating spheres were replated to obtain a second NCLC batch on day 16-18. NCLC identity was confirmed by SOX10 expression (Figure 1B). Notably, attached spheres itself displayed heterogeneous morphologies, including neural rosette-like structures characteristic of telencephalic progenitors (SI Figure 2B).

**Figure 1.**
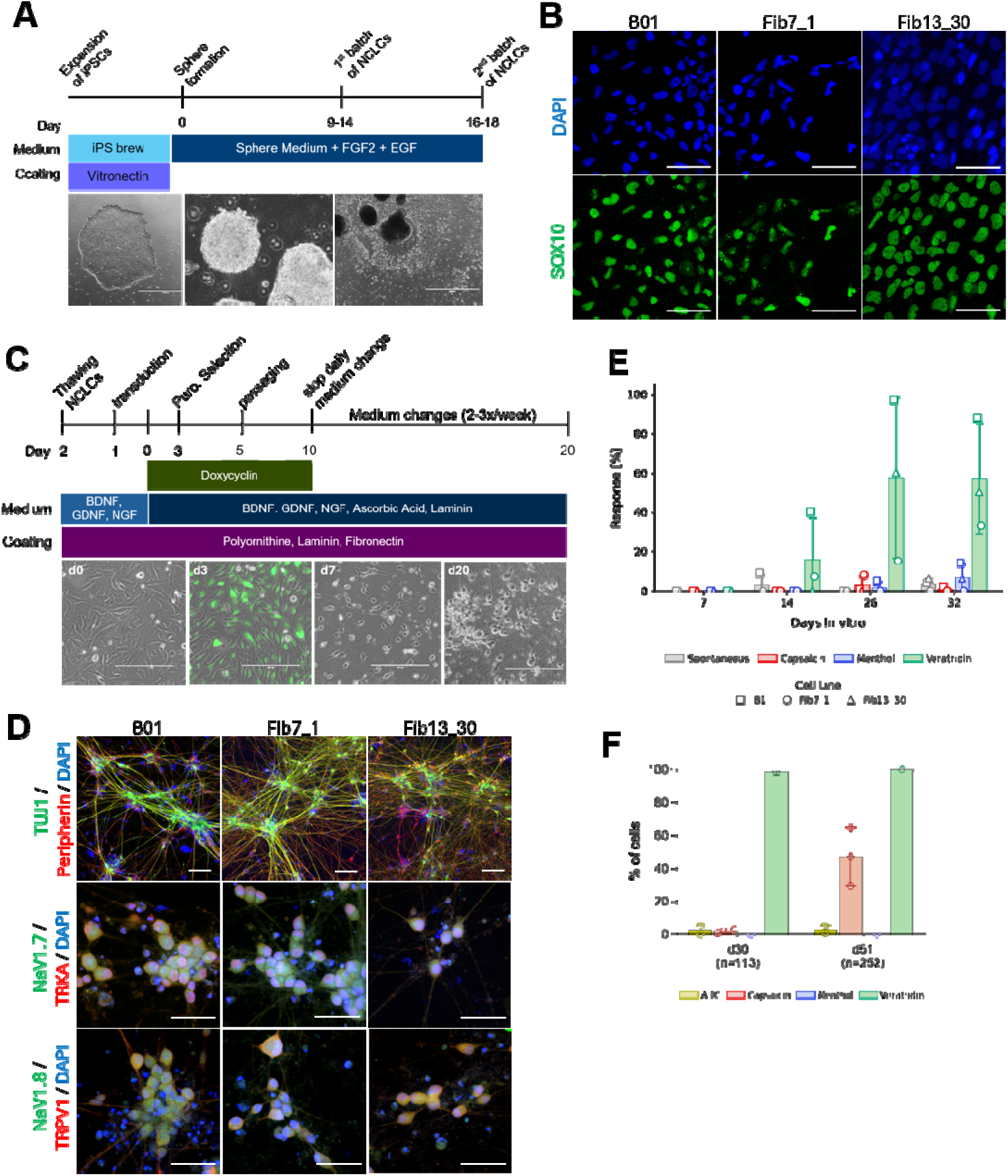
Differentiation of sensory neurons by ectopic expression of NGN1. (A) Schematic overview of the iPSC differentiation into neural crest-like cells (NCLCs). Scale bars from left to right: 200 µm, 1000 µm. (B) NCLCs derived from three independent iPSC lines express the neural crest marker SOX10. Nuclei are counterstained with DAPI. Scale bar: 50 µm. (C) NCLCs were further differentiated into sensory neurons as outlined schematically. Representative images show morphological changes, including neurite extension and the development of round, prominent somata. Scale bar: 200 µm. (D) After 20 days of differentiation, neurons express TUJ1, Peripherin, Nav1.7, Nav1.8, TRKA, and TRPV1, indicating acquisition of a nociceptor-like identity. The second antibody control (SI Figure 2A) showed no staining with the chosen imaging settings. Scale bar: 50 µm. (E) Calcium imaging was performed weekly between day 7 and day 32 of sensory-neuron differentiation. The percentage of spontaneously active cells and cells responding to menthol (TRPM8), capsaicin (TRPV1), or veratridine (Nav) is shown for each cell line. (F) Differentiation was extended to day 51 and the experiment repeated with Fib7_1. The percentage of cells responding to AITC (TRPA1), capsaicin, menthol, or veratridine is depicted. was determined from 2-3 independent experiments and are represented ± SD, with a total number of cells analyzed given by n.

The subsequent differentiation workflow is summarized in Figure 1C. Thawed NCLCs were transduced with a lentiviral mix encoding rtTA, NGN1, GFP, and a puromycin resistance cassette. NGN1 expression was induced the following day, resulting in GFP expression and enabling puromycin selection. After selection, cells transitioned from NCLCs to sensory-neuron progenitors and ultimately to neurons with round soma and dense neurite networks (Figure 1C). By day 20, neurons expressed TUJ1, Peripherin, Nav1.7, Nav1.8, TRKA, and TRPV1 (Figure 1D). A small fraction of non-neuronal cells remained detectable.

In addition, we observed that spheres were transducible with low efficiency, and neural rosette containing spheres were transduced at efficiencies comparable to NCLCs (SI Figure 2B). Those neural rosettes containing spheres express PAX6, while cells in the periphery of those spheres express SOX10 (SI Figure 2D). When neural rosettes were subjected to the sensory-neuron differentiation protocol, these cells generated neurons in less than 7 days (SI Figure 2C) and after 13 days, neurons show TUJ1 expression and absence of Peripherin pointing towards a telencephalic identity (SI Figure 2E). This highlights the need for stringent removal of spheres before sensory neuron differentiation.

To assess functional maturation of sensory neurons, we performed calcium imaging from day 7 to day 32 using veratridine, a neurotoxin that prevents inactivation of voltage-gated sodium channels (Figure 1E). Neurons do not exhibit spontaneous calcium transients. Neurons began responding to veratridine at day 14, and by day 26, all B1-derived neurons exhibited robust responses. The other two iPSC-lines matured more slowly, reaching only 30–50 % veratridine-responsive cells (Figure 1E). At these early time points, cells do not show reactivity to menthol (TRPM8-agonist) and capsaicin (TRPV1 agonist).

To further characterize sensory neuron functionality at later stages of differentiation, we tested Fib7_1-derived neurons for responses to AITC (TRPA1 agonist), capsaicin (TRPV1 agonist), and menthol (TRPM8 agonist) at d30 and d50 (Figure 1F). We found that, after 30 days of differentiation, 100 % of cells reacted to veratridine compared to only 30 % responders in the first differentiation batch, pointing towards a certain variability of maturation speed even in one cell line and between runs. In contrast to what is described in Schrenk Siemens et al. only a minor percentage of cells showed increased Ca^++^-levels in response to AITC and capsaicin and no cells responded to menthol treatment (Figure 1F). We repeated the experiment 3 weeks later. A mean of 45 % +/-20 % of cells reacted to capsaicin addition, while nearly no cells were reactive to AITC and menthol (Figure 1F).

In summary, we were able to implement the findings described and obtained NCLCs and sensory neurons based on marker expression (1). Responsiveness to veratridine was developed early albeit with an overall later response (d51 vs. d21) and lower percentage of cells responding to capsaicin (max. 60 % vs. 90 %). Thus, this protocol provides a good starting point for further optimization and standardization of key steps to increase reliability and output needed for translational applications.

### 3.2 Differentiation of NCLCs with Higher Yield

A prerequisite for the translational applicability of the protocol is to increase the efficiency for sensory neurons derived from multiple iPSC-lines. As a first step, we assessed how robustly NCLCs can be generated across a broad range of iPSC lines. In total, 31 iPSC lines from nine donors were subjected to NCLC differentiation (Figure 2, SI Figure 3). NCLCs were counted and cryopreserved as early as day 5 and not later than day 25, yielding up to 3.5×10^7^ cells from a single 10-cm dish of starting material, with an average output of 9,43×10^6^ cells at a mean collection day of 11.1.

**Figure 2.**
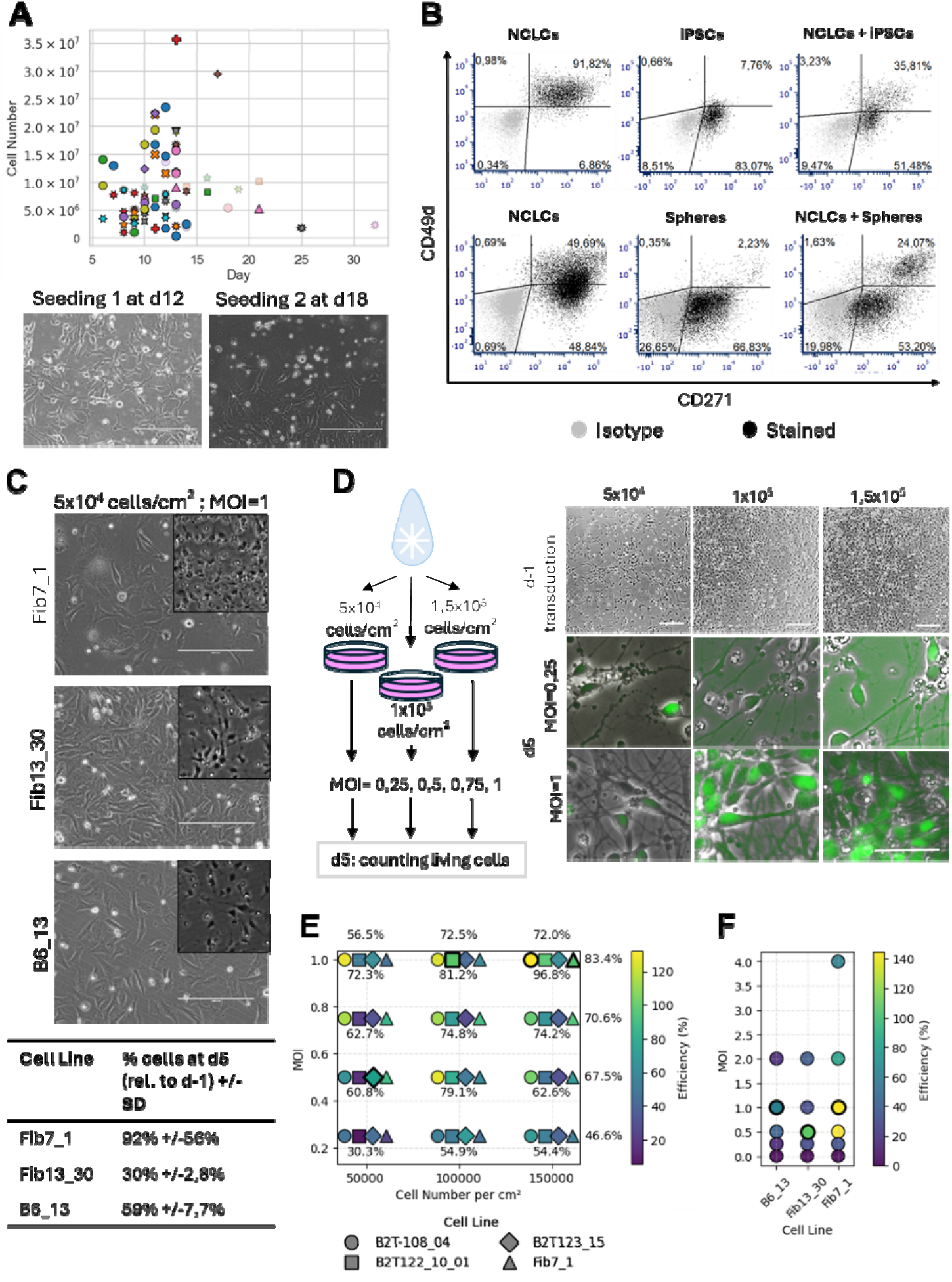
Differentiation of sensory neuron progenitors. (A) Differentiation of NCLCs from multiple iPSC lines. Time points (X-axis) and the number of cryopreserved NCLCs (Y-axis) are shown. Cell lines are color- and symbol-coded (SI Figure 3). Most NCLCs are generated during the first round of sphere attachment (dark colors), whereas the second round generally yields lower numbers. NCLC-typical morphology is maintained throughout both rounds; scale bar = 200 µm. (B) NCLCs can be identified by flow cytometry based on CD49d and CD271 expression. iPSCs and spheres do not express CD49d. Gates were set using 0.5 % positivity in the isotype control (grey). (C) Recovery of NCLCs after cryopreservation one day after thawing varies between cell lines. When transduced at MOI = 1, calculated from the initially seeded cell number, the proportion of the number of sensory neuron progenitors obtained at day 5 relative to the initial cell number seeded for transduction ranges from 30 % to 96 %, depending on viral load or cell density. Scale bar = 200 µm. (D) The MOI was adjusted to the initial seeding density, and live sensory neuron progenitors were quantified at day 5. Morphologically, progenitor yield depends on both seeding density and viral load. Exemplary images were obtained from cell line Fib7_1. Experimental scheme is shown; top scale bar = 200 µm, bottom scale bar = 50 µm. (E) Four independent cell lines seeded at three different densities were transduced with four MOIs. Cell lines are indicated by shape; color coding represents sensory neuron progenitor generation efficiency relative to the number of cells seeded. The condition with the highest efficiency is highlighted with a black outline. Efficiencies per cell number (top X-axis), MOI (right Y-axis), and condition (below symbols) are shown. (F) Three independent cell lines were transduced with higher MOIs. Again, the condition with the highest efficiency per cell line is marked with a bold outline. Efficiency is calculated relative to cells initially seeded.

To further increase the overall yield, spheres were replated after the first NCLC harvest to induce a second round of NCLC formation (Figure 2A, SI Figure 3 light colors). During this second cycle, up to 1.5×10^7^ cells were obtained (average 7.14×10^6^ cells at day 16.2), with the latest collections occurring at day 33. Morphologically, NCLCs retained their characteristic appearance throughout both rounds of differentiation (Figure 2A).

Since the purity of NCLC cultures represents a key requirement of successful sensory neuron differentiation (SI Figure 2B-E), we next aimed to establish a high-throughput assay to assess NCLC-culture purity. Based on literature, we identified two surface markers, CD49d and CD271, that could potentially allow discrimination between NCLCs, spheres, and iPSCs by flow cytometry (24–26). All three cell types were stained with this two-marker panel (Figure 2B). NCLCs expressed both CD49d and CD271, whereas spheres and iPSCs showed CD271 expression only, with substantially lower mean fluorescence intensity. To evaluate whether this panel is sufficient to quantify NCLC purity, NCLCs were mixed with either iPSCs or spheres at a 30 %/70 % ratio. The combination of CD49d and CD271 reliably resolved both populations, demonstrating that this marker set is suitable for high-throughput NCLC quality assessment in the future.

The efficiency of sensory-neuron differentiation depends not only on the yield and purity of NCLCs but also on the subsequent differentiation of cryopreserved NCLCs into sensory neuron progenitors. To test the variability between cell lines without the need to heavily expand the experiments, the line Fib7_1 was used as a reference throughout all experiments, complemented by additional iPSC lines from different donors. After thawing, NCLCs were seeded at 5×10^4^ cells/cm², which resulted in variable cell densities 24 h later due to differences in post-thaw recovery (Figure 2C). Since the multiplicity of infection (MOI) must be calculated based on the number of cells seeded, these variations directly influence the effective viral dose. When cells were transduced at MOI = 1, the proportion of sensory neuron progenitors at day 5 varied substantially between cell lines, ranging from 30 % to 92 % of initially seeded cells (Figure 2C).

For translational applications, however, a robust and reproducible generation of sufficient sensory neuron progenitors is essential to avoid time- and cost-intensive repetitions of downstream assays. To determine whether low differentiation efficiency results from insufficient cell density or from excessive viral load, NCLCs were seeded at different densities, and the MOI was adjusted accordingly, with different ratios tested for each cell number (MOI 0.25–1, Figure 2D). Sensory neuron progenitors were quantified at day 5. Morphologically, higher initial seeding densities resulted in stronger GFP expression and increased numbers of sensory neuron progenitors at MOI = 1. In contrast, low MOIs consistently produced weak GFP expression and poor progenitor output, independent of seeding density. This experiment was performed across four independent cell lines. For three of these lines, maximal progenitor yield was achieved with MOI = 1 combined with an initial seeding density of 1×10^5^ to 1.5×10^5^ cells/cm², corresponding to a differentiation efficiency ranging from 50 %-125 % to 90 % – 130 % relative to the number of cells seeded after thawing. Increasing the MOI to 2 or 4 markedly reduced efficiency (Figure 2E). These findings demonstrate that achieving maximal sensory neuron progenitor output requires a precise balance between seeding density and MOI in a cell line specific manner.

In summary, assuming that approximately 1,7×10^7^ NCLCs can be generated from a single 10-cm dish of iPSCs in average and that differentiation efficiencies of around 100 % are attainable, more than 17 million sensory neuron progenitors could be obtained by day 5-sufficient to support high-throughput experimental workflows. Notably, one cell line reached only 60 % efficiency under the most favourable conditions (B6_13, lowest density, MOI = 0.5; Figure 2E), highlighting persistent line-to-line variability.

### 3.3 Selection of Cell Culture Media for Sensory Neuron Development

Maturation of sensory neurons is usually carried out in high glucose media (up to 25mM glucose). Diabetic peripheral neuropathy is one of the most common and debilitating complications of diabetes mellitus and develops in around 50-60 % among diabetic patients (27). To develop the differentiation protocol towards broad applications, we aimed to choose a maturation medium that contains physiological glucose concentrations. For that, we compared the media composition used in 35 previously published studies (SI Table 1). In most published reports, DMEM/F12 or Neurobasal based media are used containing 17 mM - 25 mM glucose with different supplements. We selected key components from all published media and created two different media compositions, termed “physiological medium” and “core medium”. For the physiological medium, we adapted the glucose - concentration, NaPyruvate concentration and NaCl to physiological levels, whereas the core medium contains the components one finds in every medium used and reported in the literature. Sensory neuron progenitors of three independent cell lines were matured in 48-well MEA-Plates in all three media for 61 days; a time frame well described to allow Nav1.7 expression, a voltage gated sodium channel well described for its role in neuropathic pain (28). Cell lines were chosen based on previously published development of a neuropathic pain phenotype *in vitro*: a small fibre neuropathy cell line known to develop an hyperexcitable phenotype with increased firing on MEA (Fib13_30, (6)) and a cell line from a patient suffering from inherited erythromelalgia (IEM) who carries a disease causing mutation in Nav1.7 (B6_13; (29, 30). Together with a healthy control cell line, this setting was expected to identify a medium suitable for neuropathic pain modelling under physiological glucose conditions.

The spontaneous activity was recorded at 37°C (Figure 3A, SI Figure 4). Spontaneous activity occurred in all media, but the overall activity seems lower in physiological medium compared to core medium. Cells produce single electrode bursts in all media (SI Figure 4). Network bursting was not observed, and the synchrony index of firing was low indicating absence of functional neuronal network formation under all conditions (SI Figure 4). Core medium with high glucose concentrations lead to the development of a significant disease phenotype in the IEM cell line characterized by increased spontaneous firing in high glucose medium. A trend towards a disease phenotype characterized by increased firing of the SFN cell line was observed independent of the glucose concentration. Remarkably, the cell line B6_13 showed lower firing rates at low glucose levels compared to high glucose concentrations in the same medium pointing towards a glucose dependent phenotype that stresses the necessity of appropriate media composition for neuropathic pain modelling.

**Figure 3.**
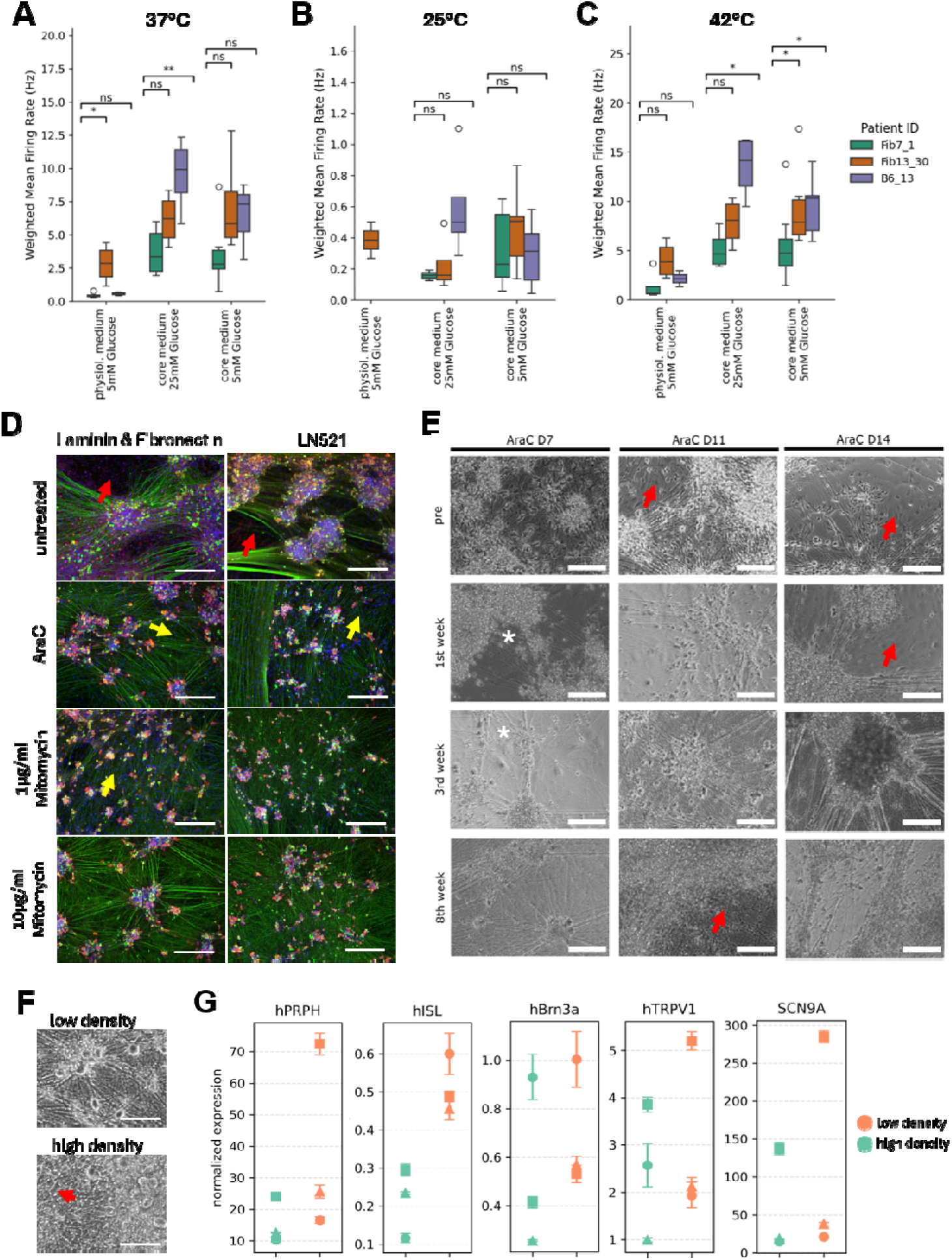
Identification of maturation medium and assessment of purity of sensory neuron cultures. (A–C) Multi-Electrode array recordings at day 61 from three independent cell lines (Fib7_1, healthy control; Fib13_30, SFN; B6_13, IEM) matured in three different media, two of which contained physiological glucose concentrations. Box plots show the weighted mean firing rate per well. Exact medium compositions are listed in SI Table 2. Spontaneous activity was recorded at 37 °C (A), 25 °C (B), and 42 °C (C). Statistical testing was performed using the Mann–Whitney U test (*p < 0.05; **p < 0.01). (D) Sensory neuron progenitors from Fib7_1 were seeded on two different coatings and treated with cytostatic selection agents to remove contaminating non-neuronal cells. Treatments were applied at day 13 using either 2 µM Ara-C for 20 h, 1 µg/ml mitomycin for 2 h, or 10 µg/ml mitomycin for 45 min. Cultures were stained for Peripherin (peripheral neuron marker, gree) and TRPV1 (nociceptor marker, red). Nuclei were counterstained with Hoechst (blue). (E) Treatment with Ara-C at day 7, day 11, or day 14 and subsequent development of sensory neuron cultures over 8 weeks. Representative images are shown. In (D, E), red arrows indicate strong contamination with non-neuronal cells leading to sensory neuron cluster fomation, yellow arrows indicate single contaminating cells without sensory neuron clusters, and white asterisks mark compromised neurites. Scale bar = 300 µm. (F, G) Seeding density at day 5 influences the purity of sensory neuron cultures at day 70, with lower densities generally yielding purer cultures. Purity was quantified by qRT-PCR (G) for Peripherin, ISL1, and BRN3A (sensory neuron markers), as well as TRPV1 and SCN9A (nociceptor-associated markers). Data represent mean ± SD of technical triplicates from three independent cell lines (symbols:) at different seeding densities (color). Normalized expression was calculated using the 2–^ΔΔCt^ method relative to the housekeeping genes GAPDH and h1C0RF43, with day 5 sensory neuron progenitors as reference.

Typically, sensory neurons respond to temperature changes by altering their spontaneous activity. All media compositions were tested at 25°C and 42°C (Figure 3B, C). Lowering the recording temperature to 25°C results in near loss of firing in all cell lines (Figure 3B, C, SI Figure 5A), and firing recovered and increased at 42°C (Figure 3C, SI Figure 5B) in all media. Still, physiological medium results in the lowest firing rate even at 42°C. The SFN and IEM cell lines show significantly higher weighted mean firing rates compared to the healthy control cell line when matured in core medium with low glucose concentration and with a recording temperature of 42 °C. This finding points towards detection of a disease phenotype at low glucose conditions and increased recording temperature albeit with high variability in the results (Figure 3C).

We chose core medium with reduced glucose for all our experiments since it allows sensory neuron maturation under low glucose concentrations and the detection of a neuropathic pain phenotype despite variability in data. To further test the functionality of sensory neurons derived in the newly developed medium, 1 µM or 10 µM capsaicin, 500 µM menthol, 250 µM AITC, and 10µM α,β-ATP were applied to sensory neurons from Fib7_1, B2T-131 and B2T-126 (cell lines from healthy donors) during MEA recordings (SI Figure 6). The neurons exhibited a transient increase in firing activity followed by a long-term reduction in spiking behaviour in response to these compounds which is expected due to desenstitization phenomenon after applying TRPV1, TRPM8 and TRPA1 agonists (SI Figure 6A). Since multiple electrodes were analyzed and the short-term increase in firing depends on the precise timing of compound arrival at the cells, we focused our analysis on the sustained effect. Therefore, we quantified the percentage of electrodes that changed their firing rate relative to their baseline activity before compound application. Pipetting alone or application of the solvents DMSO and EtOH altered activity in approximately 10–15 % of electrodes, whereas 15–30 % of electrodes showed changes in firing rate in response to the tested agonists (SI Figure 6B). Most neurons exhibited a reduction in firing frequency after stimulation (SI Figure 6C). The normalized change in firing rate relative to baseline reached statistical significance upon application of 10 µM capsaicin, 500µM menthol, and 250 µM AITC, while 10µM α,β-ATP did not reach significance, possibly reflecting distinct cellular reactivity as observed in the representative traces (SI Figure 6D).

### 3.4 Purification of sensory neuron cultures

Purity or at least defined compositions of matured sensory neuron cultures are crucial for most experiments, especially those analysing the whole cell population like expression analysis or MEA experiments. Most neuronal differentiation protocols include selection agents, like mitomycin-C or Ara-C, to reduce unwanted cell populations by inducing cell cycle arrest in actively proliferating cells leading to cell death (31, 32). Three commonly used treatment regimens were selected for further evaluation: 2 µM Ara-C for 16-24h, 1 µg/ml mitomycin for 2 hours and 10 µg/ml mitomycin for 45 minutes (1, 33, 34). Since we observed that presence of non-neuronal cells leads to cluster formation and poor attachments of neurons and coatings affect those parameters, cells were seeded to 4 different coatings: Geltrex, fibronectin and laminin, LN521 and LN521 with fibronectin. Untreated cultures show presence of non-neuronal cells, characterized by absence of Peripherin and TRPV1 staining, forming islands that push neurons towards strong cluster formation on all coatings tested (Figure 3D, SI Figure 7C, D). Ara-C-treatment and treatment with 1 µg/ml mitomycin causes clear reduction of non-neuronal cells. Still, non-neuronal cells are detectable in all cultures after 1µg/ml Mitomycin treatment albeit to lesser extent compared to untreated cultures. Ara-C treatment, on the other hand, lead to near loss of non-neuronal cells with complete absence in most cultures (Figure 3D, SI Figure 7C,D). High dose Mitomycin treatment resulted in the cleanest sensory neuron culture with most cells expressing Peripherin and TRPV1. Success of treatment and cluster formation was not dependent on coating. Attachments of cells was maintained with all coatings used throughout differentiation until d60. Since LN521, is a human recombinant protein, it allows a xeno-free sensory neuron culture and is, therefore, advantageous over all other coatings. LN521 is used in all subsequent experiments.

During iPSC differentiation to neurons, there is a critical transition period in which cells shift from dividing progenitors to postmitotic neurons. To determine the best time window for treatment, cells were treated at d7, d11 and d14 with Ara-C. Already in that short time window non-neuronal cells expanded drastically (red arrows) (Figure 3E, SI Figure 8A). Ara-C treatment as early as d7 strongly affects sensory neurons health. Signs of cell death were obvious, and neurite integrity was heavily compromised (white asterisk). However, cells recover during the next weeks and form clean, dense neurite networks and prominent soma typical for sensory neuron cultures. When treating the cells at d11, cluster formation of neurons is already observable due to increasing numbers of non-neuronal cells. Sensory neurons are less affected by treatment and still develop to typical sensory neuron cultures with absence of non-neuronal cells representing the best time point for Ara-C treatment (Figure 3E, SI Figure 8A). If treatment is performed 3 days later, a significant amount of non-neuronal cells was present at the time point of treatment and has already pushed neurons into prominent clusters. Cluster formation persisted even several weeks after treatment with non-neuronal cells still being present in culture. Still, in one cell line (SI Figure 8A right) a substantial number of contaminating cells was already present at d7 and expanded in culture even when treatment was performed at the earliest time point tested. Overall, this suggests that treatment times need to be balanced wisely to be as late as possible for neuron health but as early as possible for effectiveness. Efficiency depends on the amount of contaminating cells at the time point of treatment. To balance all results and allow high throughput differentiations, Ara-C treatment at d11 of differentiation was chosen for further experiments.

During all of our experiments we observed that not only the treatment paradigm with selection agents favours pure sensory neuron cultures but also the seeding density at d5 seems to impact on culture quality (Figure 3F, left). To test for the impact of seeding density at d5, sensory neuron progenitors were seeded to high (1,4×10^5^ – 2,1×10^5^ cells/cm²) and low (5,5×10^4^-8,3×10^4^ cells/cm²) density cultures and differentiated for 8 weeks (Figure 3F, SI Figure 8B). High density cultures display presence of non-neuronal cells while two out of three cell lines show nearly pure sensory neuron cultures when seeded at low density combined with Ara-C treatment at d11 (SI Figure 8B). In general, low density cultures express higher levels of the peripheral neuronal marker *PRPH* encoding for Peripherin, the sensory neuron marker *ISL1* and *POU3F1* and the nociceptor-associated markers Nav1.7 and TRPV1 (Figure 3G) pointing towards a beneficial effect of low density seeding for sensory neuron culture purity in addition to Ara-C treatment.

### 3.5 Maturation of Sensory Neurons

We gradually adapted the initial protocol into a standardized workflow with the overall aim of reducing workload, particularly on weekends, to allow for higher throughput of differentiations per experimenter. (SI Figure 9A). We observed that medium changes during NCLC differentiation can be performed three times per week, following a Monday–Wednesday–Friday schedule. In addition, during sensory neuron induction, the optimal timing for thawing NCLCs was found to be Monday. Freshly thawed NCLCs could be expanded for one additional day, which enabled synchronization of multiple cell lines despite variable post-thaw recovery. In cases of poor recovery (<60 % confluence) on Tuesday, additional NCLCs could be seeded into the same dish to ensure confluence for transduction the following day. Since high viral load consistently resulted in poor sensory neuron progenitor yield (Figure 2 C-F), we strongly recommend starting with low MOIs for transduction. If only few GFP-positive cells are detected at day 1, the transduction can be repeated using sphere medium supplemented with BDNF, GDNF and NGF (BGN). In this case, PBS washing and medium changes on day 2 are mandatory. With this approach, days 2, 9, and 10 (weekend) were omitted, reducing weekend workload, followed by twice-weekly medium changes after removal of cytostatic agents. We did not observe any obvious negative morphological effects associated with the reduced feeding schedule.

After 10 weeks in culture, iPSC-derived sensory neurons show multipolar morphology with dense neurite networks and prominent round soma (Figure 4B). When human DRG (hDRG) neurons are dissociated and placed into a 2D culture system, they lose their pseudounipolar structure, display round somata with diameters of 28–60 µm, and are frequently associated with glial support cells (35). In contrast, although iPSC-derived neurons in our standard maturation conditions also develop round, prominent soma, they reach markedly smaller sizes (median: 24 µm) after 70 days in culture (Figure 4 B-D, untreated condition).

**Figure 4.**
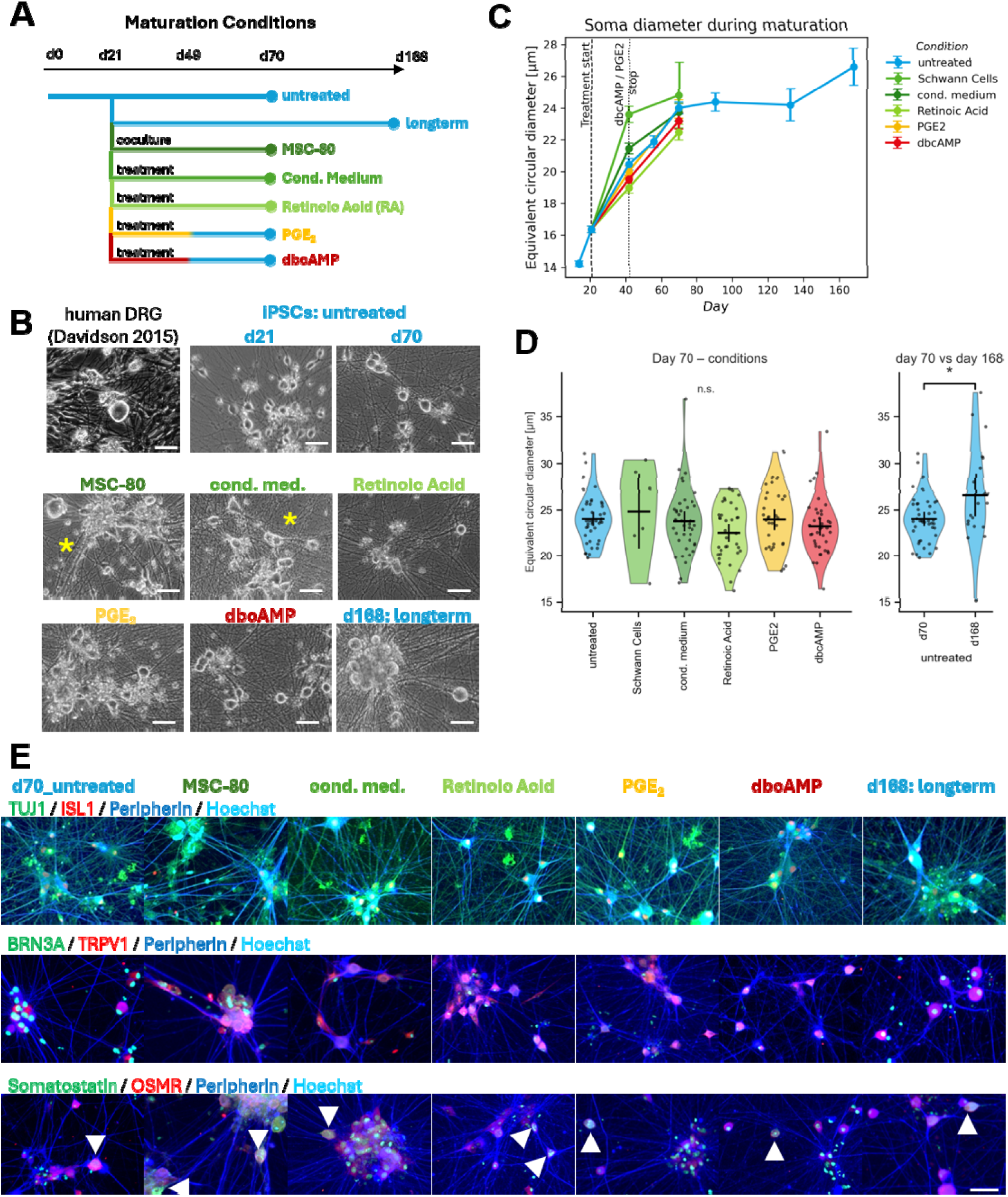
Maturation conditions for iPS-derived sensory neurons aiming to resemble hDRGs morphologically. (A) Maturation scheme and treatment timeline for sensory neuron differentiation. (B) hDRG morphology *in vitro* (35) (permission requested) compared with iPSC-derived sensory neurons after 70 days of maturation with the following conditions: in co-culture with MSC-80 cells, or after treatment with MSC-80 conditioned medium, prostaglandin E2 (PGE2), retinoic acid (RA), dbcAMP, or after long-term culture until day 168. Scale bar = 50 µm. Co-culture with MSC-80 or maturation in 50 % MSC-80 conditioned medium compromises neurite morphology (*) (C) Analysis of the surface-equivalent soma diameter over the course of iPSC-sensory neuron differentiation did not reveal differences between maturation conditions. (D) Distribution of soma diameters at day 70 across all conditions showed no significant differences (Kruskal–Wallis p = 2.8637 × 10⁻¹). After 168 days of maturation, soma diameter increased significantly (Mann–Whitney U test p = 3.2267 × 10⁻²). D70 meassurments are shown in the left and right figure for better comparison.(E) Across all conditions, neurons expressed TUJ1, Peripherin, ISL1, and TRPV1. BRN3A was not clearly detectable. Individual cells co-expressed OSMR and somatostatin (arrows). Scale bar = 50 µm.

To generate iPSC-derived sensory neurons that more closely resemble hDRG neurons *in vitro*, we tested six different maturation conditions aimed at promoting a more hDRG-like phenotype (Figure 4A). First, neurons were matured in co-culture with the mouse Schwann cell line MSC-80 to provide glial support and mimic *in vivo* cell–cell interactions. To avoid the presence of mouse cells while still supplying Schwann-cell-derived factors, we next supplemented core medium with 50 % MSC-80 conditioned medium. In a third approach, we added high-dose retinoic acid (RA) from day 21 onward, as RA has been reported to promote neurite outgrowth, support Schwann cell development, and induce PRDM12 expression—a transcription factor required for sensory neuron development (36–38). In addition, from day 21 to day 49, we supplemented the maturation medium with either prostaglandin E2 (PGE2) or dbcAMP. PGE2 is a known component of Schwann-cell-conditioned medium and increases expression of Nav1.7 and Nav1.8 in embryonic rat DRGs (39). dbcAMP has been shown to promote survival and neurite outgrowth in rat sensory neurons (40), and embryonic DRG-derived cell lines acquire more mature neuronal phenotypes in its presence (41). Notably, embryonic DRG neurons naturally exhibit high cAMP levels, which decline during development, suggesting that temporary limited addition could favour maturation (42). Lastly, the maturation time frame was extended to 6 months (168 days).

Morphologically, we did not detect obvious differences between the dbcAMP-, PGE_2_-, and RA-containing maturation conditions up to day 70 of differentiation (Figure 4B). In contrast, cultures matured in the presence of MSC-80 Schwann cells or Schwann-cell-conditioned medium showed signs of cell death, neurite disassembly, and detachment (yellow asterisk). Soma diameter equivalent to the measured soma area, increases during maturation reaching the maximum size with 22-24 µm at d70 (Figure 4C, SI Figure 9 B-D). Endpoint analysis of the soma size did not differ significantly between conditions (Figure 4C, D, SI Figure 9B-D). Prolonged maturation for six months resulted in soma sizes reaching up to 37 µm in diameter, with a median diameter of 27 µm (Mann–Whitney U-Test d70 vs d168: p = 3.2267×10^-2^), approaching the size of hDRG neuron diameters reported by Davidson et al. (Figure 4D and SI Figure 9 B, D). Neurons matured under all tested conditions expressed the pan-neuronal marker TUJ1, the peripheral neuron marker Peripherin, the sensory neuron marker ISL1, and the nociceptor marker TRPV1 (Figure 4E, SI Figure 10A-D). Thus, neither of the chosen conditions can be preferred over the other. Individual neurons also displayed co-expression of OSMR and somatostatin, markers associated with silent nociceptors (43), pointing to the presence of different sensory neuron types in our protocol.

### 3.6 RNAseq Revealed PGE2 as a Promising Supplement for Sensory Neuron Maturation

All maturation conditions directed cells toward sensory neuron fates, as confirmed by immunostaining for key markers (Figure 4E, SI Figure 10). To assess transcriptional differences between treatments, we performed bulk RNAseq on cultures at day 70 and on long-term matured neurons at day 168 (Figure 5). Across all conditions, cultures expressed high levels of pan-neuronal markers (MAP2, TUBB3, RBFOX3, SYN1, SYP, STMN2) and lower levels of developmental regulators such as PRDM12 and ASCL1 (Figure 5A). Genes associated with nociceptor, mechanoreceptor, and proprioceptor identities were detectable under all maturation conditions, with no obvious qualitative differences between treatments or cell lines (Figure 5A, SI Figure 11A-B). Ion channel expression was broad, including all α- and β-subunits of voltage-gated sodium channels (with *SCN4A* at lowest levels), as well as potassium, calcium, and HCN channels (SI Figure 11C-F).

**Figure 5.**
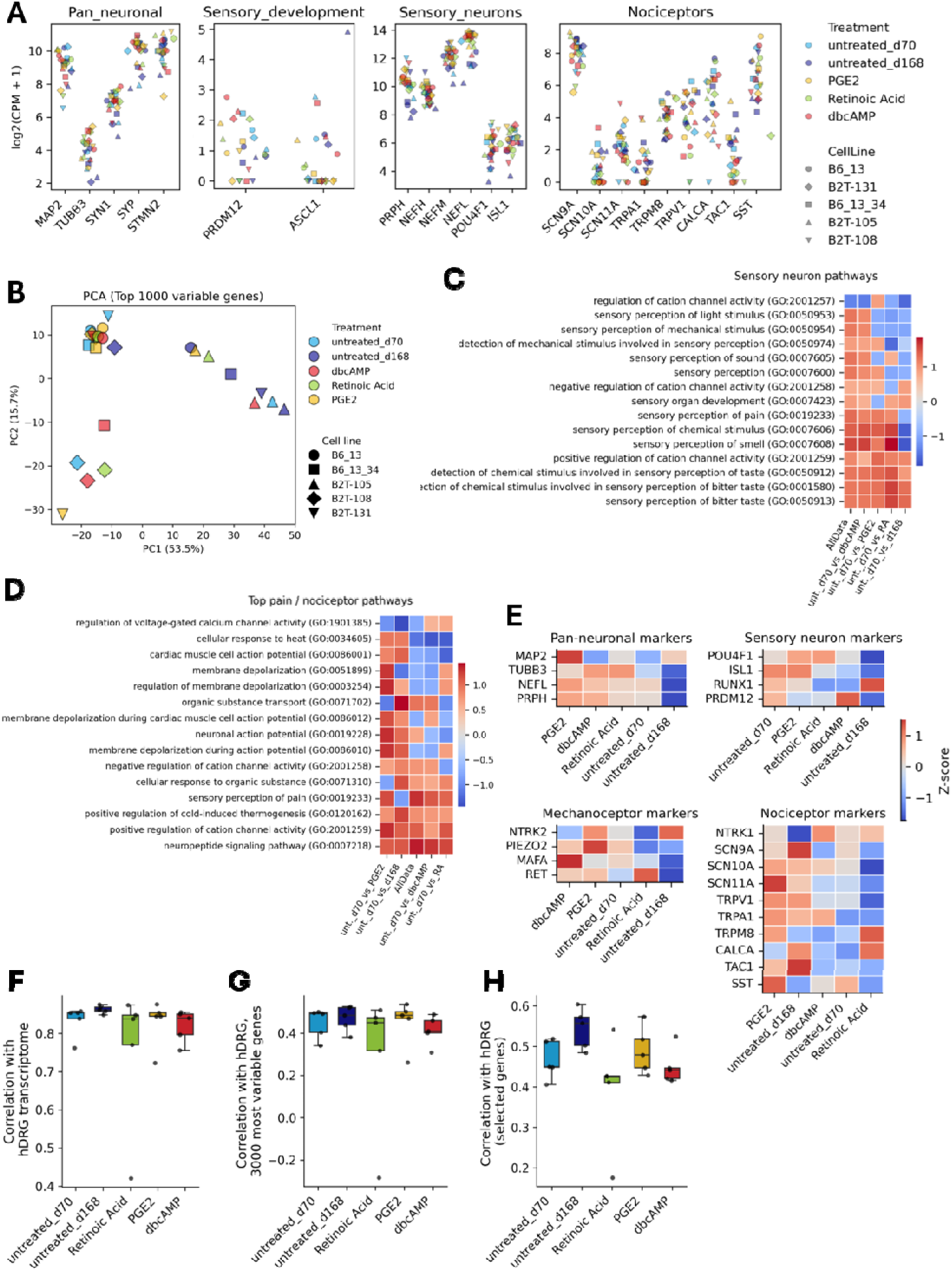
Transcriptomic characterization of iPSC-derived sensory neurons under different maturation conditions. (A) Expression levels of pan-neuronal, sensory-developmental, sensory-neuron, and nociceptor-associated gene groups across all maturation conditions. Each point represents one sample, color-coded by treatment and shaped by cell line. (B) Principal component analysis (PCA) of the top 1000 most variable genes. PC1 (53.5 %) separates long-term matured neurons (day 168) from day 70 samples, while PC2 (15.7 %) distinguishes a subset of samples primarily from cell line B2T-108. (C-D) Gene set enrichment analysis of sensory and pain-related pathways based on the mean from all cell lines compared to the untreated sample at d70 (unt.70). Pathways are ordered by expression level with overall highest expression on the lower left and lowest expression on the upper right side. Pathways are enriched across all conditions. dbcAMP-treatment produces the highest cumulative presence of sensory related pathways. PGE2-treated cultures display the highest overall enrichment across pain-related gene sets. (E) Expression of marker genes for pan-neuronal identity (*MAP2, TUBB3, NEFL, PRPH*), sensory neuron development (*POU4F1, ISL1, RUNX1, PRDM12*), mechanoreceptors (*NTRK2, PIEZO2, MAFA, RET*), and nociceptors (*SCN9A, SCN10A, SCN11A, TRPV1, TRPA1, TRPM8, CALCA, TAC1*). Conditons and genes are orderd based on highest to lowest expression level. PGE_2_ treatment results in the highest expression of pan-neuronal, mechanoceptor and nociceptor markers, while developmental sensory neuron markers remain low. (F-H) Correlation of iPSC-derived neurons with a pseudo-bulk human DRG reference (23) based on all detected genes (F), the 3000 most variable genes (G) and selected sensory-neuron marker sets (SI Table 5) (H). Although correlations do not reach statistical significance (ANOVA), long-term maturation and PGE_2_ treatment show the highest similarity trends to human DRG. Abbreviations: unt._70 = untreated at d70 of differentiation, vs = versus, RA = Retinoic Acid.

Principal component analysis (PCA) of the top 1000 most variable genes separated the samples into three major groups (Figure 5B): One group consisted predominantly of long-term matured neurons (day 168), which were separated along PC1 (53.5 % variance), indicating that extended maturation induces a distinct transcriptional state. Day 70 samples formed two groups: dbcAMP-, RA-, and PGE_2_-treated neurons group together, whereas a second group was composed mainly of samples from cell line B2T-108 (PC2, 15.7 % variance). Overall, samples separate more strongly by cell line than by treatment, suggesting that donor-specific effects and maturation time dominate over treatment-specific transcriptional changes at day 70.

To evaluate whether specific treatments promote sensory or nociceptor identities, we performed DESeq2 comparisons between untreated day 70 samples and matched treated samples, followed by gene set enrichment analysis using Gene Ontology Biological Process (GO:BP, 2021 release) gene sets obtained via Enrichr (Xie et al., 2021) (Figure 5C, SI Figure 12). Neuronal gene sets related to synapse, membrane, and axon/dendrite biology were enriched across all conditions, whereas developmental gene sets were only weakly represented. PGE_2_-treated cultures showed the highest overall enrichment across neuronal and sensory-related pathways, suggesting enhanced neuronal differentiation (Figure 5C). For sensory neuron-specific gene sets, dbcAMP-treated neurons showed the highest cumulative enrichment, followed by PGE_2_, although, as expected, several sensory perception pathways (light, mechanical, sound, sensory organ development) were only weakly expressed in PGE_2_-treated cultures. This pattern suggests that PGE_2_ may bias differentiation toward a nociceptor-like identity rather than broadly enhancing sensory neuron diversity. Consistent with this, analysis of pain- and nociception-related pathways (Figure 5D) showed that PGE_2_-treated neurons exhibited the highest overall expression followed by long-term matured neurons and dbcAMP and RA treatment.

Marker-level analysis supported these findings (Figure 5E). PGE_2_ treatment resulted in the highest expression of mature sensory neuron markers, whereas developmental markers (*PRDM12, RUNX1, NTRK1, NTRK2*) remained low. Nociceptor-associated sodium channels (*Nav1.7, Nav1.8, Nav1.9*) were expressed at low to moderate levels across conditions, with increased expression after long-term maturation and highest levels in PGE_2_-treated neurons. A similar pattern was observed for TRP channels (*TRPV1, TRPM8, TRPA1*). Peptidergic nociceptor markers *TAC1* and *CALCA* increased primarily after long-term maturation, with lower expression in RA- and PGE2-treated cultures. Somatostatin (*SST*), a marker of silent nociceptors implicated in neuropathic pain (43), was strongly upregulated in PGE_2_-treated neurons and only weakly expressed in other conditions. Together, these findings indicate that PGE_2_ treatment between day 21 and day 49 during NGN1-driven differentiation promotes the emergence or maturation of nociceptor-like neurons. Mechanoreceptor-associated genes such as *PIEZO2, NTRK2, MAFA* and *RET* were inconsistently expressed across all maturation conditions, indicating that a subset of cells retains or acquires mechanosensory features depending on maturation, with PGE_2_, showing highest expression across all markers.

Finally, to assess similarity to hDRG neurons, we compared our samples to a pseudo-bulk reference generated from single-nucleus RNA-seq data (23). Correlation values (Pearson’s r) range from −1 to 1, with higher values indicating greater transcriptional similarity to the hDRG reference. Correlations based on all genes and the 3000 most variable genes (Figure 5F,G), as well as selected sensory neuron marker sets (Figure 5H), did not reach statistical significance. Nevertheless, long-term matured neurons and PGE_2_-treated cultures showed the highest overall similarity trends to the hDRG reference. This trend is more pronounced when a custom gene list (SI Table 5) is used for comparison. When restricting the analysis to selected sensory neuron marker genes (Figure 5H), clearer trends emerged, with long-term matured neurons and PGE_2_-treated cultures showing higher similarity to the hDRG reference, while RA-treated samples displayed lower similarity.

### 3.7 Functional Maturation of Sensory Neurons on MEA

To assess the functional maturation of sensory neurons in the presence of Retinoic Acid, dbcAMP or PGE_2_, and during long-term differentiation at defined cell densities, extracellular neuronal activity was recorded using MEA across multiple time points and five cell lines (Figure 6, SI Figure 13). Spontaneous activity emerged around day 14, and the mean firing rate peaked between day 42–50 in all conditions, apart from of the 60,000-cells-per-chip condition, which showed a delayed peak after day 100 (Figure 6A, SI Figure 13). After reaching a maximum mean firing rate of approximately 1.5–2 Hz, activity declined over time to 0.2–0.4 Hz and decreased further during extended maturation. Burst frequency (i.e. the number of bursts occurring during recording time) followed a similar trajectory, peaking between day 42–50 at 0.2–0.3 Hz and subsequently dropping to <0.1–0.2 Hz even after long-term differentiation. Only the 60,000-cell condition maintained elevated burst frequencies throughout maturation. All MEA parameters showed substantial variability across time points and conditions (SI Figure 13).

**Figure 6.**
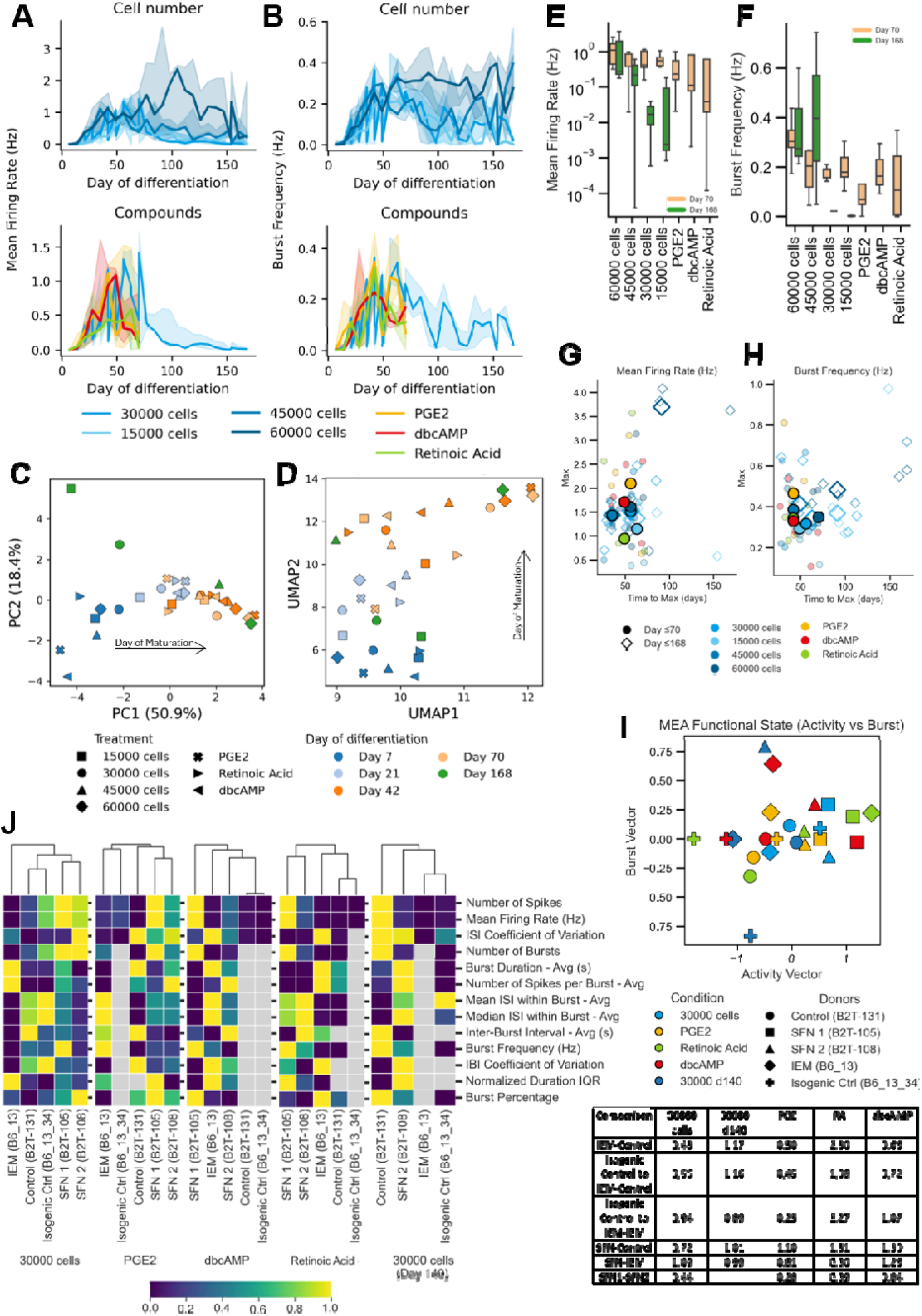
MEA-based functional characterization of sensory neurons matured under different conditions. (A–B) Development of mean firing rate (Hz) and burst frequency (Hz) during neuronal maturation. Curves represent median ± Interquartile Range across five independent cell lines matured in the presence of PGE_2_, Retinoic Acid, dbcAMP or with varying seeding cell numbers. (C–D) Principal component analysis (PCA) and UMAP of the median MEA parameters across donors at defined maturation time points (color coded). Samples cluster primarily by maturation time rather than treatment (shape). (E–F) Comparison of mean firing rate and burst frequency at day 70 and day 168 across maturation conditions. No significant differences were detected. (G–H) Maximum mean firing rate and burst frequency and the corresponding time-to-maximum for each condition. Except for the 60,000-cell condition, all treatments reached their maximal activity before day 70. Fante colors represent the mean per cell line and bright colors show median per treatment (I) To assess donor-specific functional differences relevant for pain-disease modelling, each patient line was analyzed separately. An activity vector (mean firing rate, ISI coefficient of variation, number of spikes) and a burst vector (all burst-related metrics (see J)) were computed. Samples from all maturation conditions scatter broadly, with PGE_2_ showing the smallest inter-patient distance. Table (lower panel) shows euklidian distances between indicated samples. (J) Unsupervised clustering of the median MEA parameters across all donors and maturation conditions. Cultures matured with 30,000 cells, with or without PGE_2_ or RA, cluster both SFN patient lines (B2T-105 and B2T-108) closely together. The IEM line (B6_13) clusters near its CRISPR-corrected counterpart (B6_13_34) only under PGE_2_ maturation and after prolonged maturation with 30000 cells until d140.

To account for electrophysiological development over time when assessing treatment-specific effects, the median of all MEA parameters extracted from the recording software from five independent cell lines across all conditions was subjected to principal component analysis (PCA) and UMAP at defined time points (day 7, 21, 42, 70 and 168; Figure 6C, D, SI Figure 14). In both analyses, samples clustered primarily by differentiation stage rather than treatment. Recordings from day 42 and day 70 grouped closely, whereas day 168 samples were broadly dispersed. PC1 was dominated by spike- and burst-related activity metrics, indicating that global activity was the primary driver of sample separation. PC2 reflected temporal burst structure, suggesting that changes in burst organization contributed to later-stage variability (SI Figure 14A). UMAP feature contributions mirrored the PCA loadings, with UMAP1 dominated by activity-related parameters (MFR, spikes, burst frequency) and UMAP2 reflecting burst-structure metrics (IBI, ISI variability, burst duration), indicating that both global activity and temporal burst organization shape the electrophysiological landscape (SI Figure 14B).

Since overall developmental differences are mainly driven by time than by treatments, d70 and d168 was analysed in depth (Figure 6E, F, SI Figure 15). Mean firing rate and burst frequency differed by cell density, with higher densities trending toward higher activity at day 70. Both parameters were reduced after long-term differentiation. In cultures treated with small molecules (dbcAMP, Retinoic Acid, PGE_2_), mean firing rate and burst frequency were generally lower than in the 30,000-cell reference condition, and no significant differences were detected between the treatments. Even when comparing the time to reach the maximum mean firing rate or burst frequency relative to the peak value, no treatment-specific differences were observed (Figure 6G, H, SI Figure 16).

In summary, sensory neurons exhibited characteristic developmental stages, with increasing activity early during maturation and a pronounced activity and bursting peak between day 42–50, followed by a marked decline during further development. This early activity peak is consistent with known developmental stages of embryonic neuronal development (44). The persistent high activity in the 60,000-cell condition likely reflects density-driven simultaneous multi cell recordings from each electrode rather than altered maturation. Although maturation treatments induced mRNA-transcriptional changes, these did not translate into detectable functional differences in global MEA recordings, and it becomes clear that the duration of differentiation has a larger impact.

### 3.8 Modelling of Neuropathic Pain

Patient-specific modelling of nociceptive drive underlying neuropathic pain is one of the most promising applications of this differentiation protocol. Therefore, the optimal maturation condition should reliably support the detection of disease-relevant electrophysiological phenotypes. Given that three of the five donor lines originated from individuals with neuropathic pain, we next assessed whether donor genetic background influenced electrophysiological maturation (Figure 6 I,J). In all maturation experiments, we included two independent cell lines from patients with SFN associated with gain-of-function variants in *SCN10A* (p.Ile1225Thr in SCN10A of unknown significance; SFN1 (B2T-105)) or *SCN11A* (Tyr66Ser in SCN11A,likely pathogenic (45, 46); SFN2 (B2T-108)), and one line from a patient with IEM carrying a gain-of-function mutation in SCN9A (p.Q875E (29, 47); B6_13). The Nav1.7 mutation in the IEM line had been repaired by CRISPR-Cas9, generating an isogenic control. A fifth line (B2T-131, Control) served as an unrelated healthy control. To determine which maturation condition is most suitable for MEA-based disease modelling, we hypothesized that (i) the two SFN lines should group closely together due to their shared disease diagnosis, (ii) the healthy control should separate from the SFN lines, and (iii) the IEM line should display altered excitability relative to its CRISPR-repaired counterpart, which should in turn resemble the healthy control while retaining donor-specific background features.

To compare overall activity across the five lines differentiated with PGE_2_, Retinoic Acid, dbcAMP, or under long-term conditions plated at 30,000 cells per MEA chip, an activity vector and a burst vector were computed from all available MEA parameters for each sample (cell line × treatment × day 70 or day 168) and plotted in a two-dimensional activity–burst space (Figure 6I). Samples distributed primarily along the activity axis, with only minor separation along the burst axis. The smallest distances between cell lines were observed in untreated 30,000-cell cultures and in cultures treated with PGE_2_ or dbcAMP, whereas long-term differentiation and Retinoic Acid treatment produced the greatest spread. Under PGE_2_ maturation, all samples group relatively close to each other, indicating similar differentiation into sensory neurons. Still, the two SFN lines localized close to each other across both vectors and were separated from the healthy control, consistent with the expected disease-related phenotype similarity. The CRISPR-repaired IEM line positioned between the IEM and control lines, with separation driven mainly by burst-related parameters suggesting partial rescue of the phenotype while retaining donor-specific features.

Unsupervised clustering of spontaneous activity under these conditions mirrored these relationships (Figure 6J). Cultures matured with 30,000 cells (untreated), PGE_2_, or Retinoic Acid showed consistent clustering of the two SFN lines, while the IEM, CRISPR-repaired, and control lines displayed variable clustering patterns (Fig. 6J). The healthy control line (B2T-131) formed a distinct branch only after long-term differentiation, however, under other conditions, it clustered closely with the CRISPR-repaired line. The IEM and CRISPR-repaired lines showed the greatest similarity under PGE2 and long-term differentiation, whereas in other conditions the CRISPR-repaired line tended to cluster closer to the healthy control.

Phenotypically, across all maturation conditions, both SFN cell lines exhibited the highest mean firing rates and the highest number of bursts (Figure 6J). This pattern suggests that the SFN phenotype is reflected by a general increase in spontaneous activity and altered burst structure, independent of the maturation condition. In contrast, the IEM line, its CRISPR-repaired counterpart, and the healthy control showed overall lower activity, with very low burst numbers or complete absence of bursts in the control and CRISPR-repaired lines (Figure 6J). When comparing burst duration, the IEM line tended to generate longer bursts than either the CRISPR-repaired or control line, indicating a mutation-associated alteration in burst dynamics. Interestingly, long-term differentiation resulted in a higher mean firing rate in the control line compared to the SFN lines (Figure 6J), highlighting that maturation stage can modulate donor-specific activity patterns. These patterns highlight that donor-specific electrophysiological signatures are present but are strongly modulated by maturation condition and are heavily affected by the data variability inherent to MEA-based data sets.

Overall, the patterns observed across donors and treatments are consistent with our initial hypothesis. Across conditions, SFN lines most consistently grouped together under PGE_2_, Retinoic Acid and 30,000 cells (untreated) conditions, whereas separation between the IEM and CRISPR-repaired line was more apparent with a seeding density of 30000 cells (untreated), dbcAMP, Retinoic Acid and long-term differentiation conditions where the CRISPR-repaired line clustered closer to the healthy control than to the IEM line. However, under PGE_2_ treatment, the CRISPR repaired cell line resembles more closely its IEM cell line.

Morphological analysis, RNA-seq, and MEA recordings did not clearly identify a single universally optimal condition because of high variability and donor-specific effects. However, the data indicate that PGE_2_ or 30,000 untreated cells differentiated for 70 or 168 days most reliably reveal disease-associated similarities (e.g., between SFN lines) in spontaneous activity. This coincides with the highest expression of sensory-neuron, nociceptor, and pain-associated genes.

## 4. Discussion

iPSCs possess the ability to differentiate into virtually every cell type of the human body while retaining the donor-specific genotype. This makes them particularly suitable for developing patient-specific *in vitro* disease models from somatic cell types that are otherwise difficult to obtain from living individuals. To fully exploit this potential, robust, reproducible, and well-characterized differentiation protocols are required to direct pluripotent stem cells towards a defined somatic fate. Such protocols form the basis for personalized disease modelling and drug screening, especially when only minor changes in functional aspects of specific somatic cells are sufficient to describe the individual cellular disease phenotype. Here, to establish such a protocol suitable to model nociceptive activity underlying neuropathic pain, we deconstructed the NGN1-driven differentiation method for sensory neurons from iPSCs published by Schrenk-Siemens et al. into its essential components: NCLC differentiation, lentiviral transduction, sensory neuron differentiation and maturation. We systematically characterized each step with the aim of generating a standardized robust workflow that reliably yields high-quality sensory neurons from multiple donors.

We selected the NGN1-driven protocol because its two-step structure offers unique advantages: (i) differentiation of iPSCs into NCLCs, enabling the generation of well-characterized cryostocks, followed by (ii) differentiation of NCLCs into sensory neurons through ectopic expression of NGN1, which synchronizes the maturation timeline.

We implemented the published protocol (1) in our laboratory without excessive effort and successfully obtained NCLCs and sensory neurons expressing key lineage markers and showing capsaicin responsiveness at late stages. Overall, however, the differentiation timeline appeared slower in our hands. Spontaneous attachment of Spheres to uncoated culture dishes, followed by migration, were previously reported to occur between day 6 and day 9 (1, 48). The literature does not specify the day of cryopreservation, or the number of NCLCs obtained. Based on our experience, cryopreservation occurs 1–3 days after attachment. In our hands, some iPSC lines were ready for cryopreservation as early as day 6, most between day 10–15, and others only after 21 days - slightly later than reported (1, 48). Schrenk-Siemens et al. reported capsaicin responsiveness in 92.2 ± 5.9 % of neurons after at least three weeks of differentiation. Even after 50 days in culture, we reached a maximum of 65 % in one cell line. At earlier time points, we did not detect clear responses to capsaicin, menthol, or AITC in calcium imaging experiments. It is important to note that iPSCs inherently exhibit heterogeneity, and as our study did not include the original cell lines used by Schrenk-Siemens et al., discrepancies were expected. iPSC-Culture conditions may further contribute to variability. Schwartzentruber et al. demonstrated significant differences between sensory neurons derived from iPSCs cultured on mouse embryonic fibroblasts versus Matrigel in E8 medium. E8-derived neurons showed higher neuronal content in RNA-seq data, likely reflecting lineage priming (16). The NGN1 protocol was originally published using Matrigel and home-made E8 medium, whereas we used commercially available iPS Stem MACS Brew medium (Miltenyi Biotec) combined with Vitronectin XF (Gibco) coating. Thus, our starting material likely differed from that used in the original publication. Nevertheless, all three cell lines initially used generated NCLCs and sensory neurons with appropriate marker expression, highlighting both the robustness of the protocol and the need for further standardization across multiple cell lines.

The inherent variability of iPSCs throughout the differentiation process must be considered when designing a protocol suitable for high-throughput patient-specific disease modelling and drug screening. To address this, we used the same cell line (Fib7_1) across all initial steps of the workflow and included different independent cell lines for validation. For each step, the resulting conditions were directly applied to the next step, where they were validated with different cell lines. This sequential, cross-validated approach ensured that each component of the protocol performs robustly across genetically distinct backgrounds. As a result, the final protocol integrates optimization steps derived from multiple donors and is therefore broadly applicable to most, albeit not all, cell lines. The resulting protocol provides flexibility to accommodate cell-line-specific differences. iPSCs subjected to NCLC differentiation required different timings for cryopreservation and yielded variable numbers of NCLCs, likely due to lineage priming. Cryostocking allowed us to compensate for these discrepancies. After thawing, we incorporated one day of expansion before transduction to adjust cell density even after poor recovery.

A dense layer of NCLCs was transduced with lentiviruses encoding NGN1 and rtTA. Viral load was a key determinant of sensory neuron progenitor yield at day 5. Low multiplicities of infection (MOIs) resulted in insufficient progenitor numbers, whereas high MOIs consistently reduced viability, indicating that NCLCs are particularly sensitive to lentiviral burden. This is consistent with previous reports showing that high VSV-G exposure can promote membrane fusion and fusion-associated cellular stress and cytotoxicity (49–52). High MOIs also increase transgene copy number and expression; GFP overexpression alone can induce oxidative and proteotoxic stress (53–56). To balance transgene expression and viability, we identified an MOI of 0.75 combined with 100 % cell density as optimal across cell lines. Lower densities, even with adjusted MOIs, resulted in reduced cell numbers at day 5. When density was too low, reducing viral load and repeating transduction on day 1 without altering the overall timeline allowed partial compensation for poor recovery or suboptimal virus quality.

In this stepwise optimization process, we also identified several parameters - aside from transduction conditions - that were consistently critical across all cell lines and therefore represent fixed criteria in the final protocol: One of the most influential factors was the combination of seeding density of sensory neuron progenitors combined with cytostatic treatment to obtain pure sensory neuron cultures. We compared Ara-C and Mitomycin-C treatment at day 11 and observed that both agents improved culture purity compared to untreated controls. Short, high-dose Mitomycin-C treatment resulted in the cleanest cultures. Nevertheless, we selected Ara-C for all subsequent experiments because it had already been established in our laboratory for small-molecule–driven sensory neuron differentiation from iPSCs (57, 58). Both Ara-C and Mitomycin-C target actively dividing cells through DNA-replication–dependent mechanisms. Ara-C requires incorporation into replicating DNA during S-phase to exert its cytotoxic effects (59), whereas Mitomycin-C acts as a DNA cross-linker that interferes with DNA replication (34). During iPSC differentiation to neurons, there is a critical transition period where cells shift from dividing progenitors to postmitotic neurons. Accordingly, we observed neuronal damage and neurite disintegration when cytostatic treatment was applied too early during maturation. Unfortunately, Ara-C has been also reported to kill postmitotic embryonic chicken DRG neurons in a concentration dependent manner (60). Most published sensory neuron differentiation protocols, therefore, rely on Mitomycin-C (1, 61). Mitomycin-C has also been successfully integrated into several CNS-neuron differentiation procedures. For example, Hiller et al. demonstrated that Mitomycin-C treatment during differentiation of human iPSC-derived midbrain dopaminergic neurons effectively reduced proliferating non-neuronal cells without impairing neuronal viability, function, or *in vivo* engraftment (34). These findings support the notion that Mitomycin-C is generally more neuron-sparing than Ara-C. In the future, a detailed comparative analysis of both cytostatic agents in our sensory neuron system will be required to identify the most effective agent with the lowest negative impact on neuronal health and maturation.

A second universally critical parameter identified across all cell lines was the seeding density of sensory neuron progenitors. In contrast to the NCLC stage, where high densities supported efficient transduction, we found that sensory neuron progenitors are best seeded at low densities to obtain homogenous, high-purity neuronal cultures. High-density cultures consistently resulted in the expansion of residual proliferative cells and the emergence of non-neuronal contaminants, leading to mixed cultures that could not be rescued by subsequent Ara-C treatment, as reflected by reduced expression of sensory neuron markers. This indicates that once sensory neuron progenitors are seeded too densely, proliferative cells gain a competitive advantage that is no longer reversible by anti-mitotic agents. Thus, low-density seeding emerged as a fixed criterion for achieving high-purity sensory neuron cultures. However, the exact optimal density appears to be cell-line dependent and requires further evaluation. These observations suggest that cell–cell contact and local microenvironmental cues strongly influence lineage commitment at this stage. Notch signalling, which is activated through direct cell-cell contact, plays a central role in coordinating differentiation decisions during development (62). In zebrafish sensory neuron development, Notch signalling inhibits neurogenesis, and inhibition of Notch through DAPT is a key component of widely used small-molecule–driven sensory neuron differentiation protocols based on Chambers et al. (61, 63). Consequently, reduced cell–cell contact at low seeding densities likely decreases endogenous Notch activation, thereby promoting sensory neuron maturation and limiting the initial expansion of non-neuronal cells. This early reduction in non-neuronal cell numbers can then be further stabilized by cytostatic treatment, provided that treatment occurs at a stage when contaminating cells are still sparse.

The stepwise nature of the NGN1-driven protocol offers the unique opportunity to integrate three natural checkpoints into the differentiation timeline, which may in the future allow early prediction of differentiation success. As a first checkpoint, we established FACS as a quality-control step for NCLCs and selected CD49d and CD271 as markers. Migratory neural crest cells derived from iPSCs are known to express both markers and retain the competence to differentiate into glial cells and neurons. In the absence of SOX10, the generation of CD271 (p75) and CD49d-positive migratory neural crest cells is impaired, resulting in significantly reduced numbers of Peripherin-, TUJ1-, GFAP- and S100B-positive cells upon differentiation into neurons or glia, respectively (24). This supports the assumption that the NCLCs generated in our system are multipotent and not only capable of differentiating into sensory neurons upon NGN1 induction, but also into Schwann cells and potentially additional neural crest-derived lineages when exposed to appropriate patterning cues (48). We also observed that while CD271 is broadly expressed, CD49d is sufficient to discriminate between iPSCs, spheres, and NCLCs. Spheres frequently contain neural rosettes, a characteristic morphological feature of anterior CNS progenitors, which can give rise to central nervous system neurons and glia when exposed to defined patterning signals (64). Consistent with this, NGN1-transduced spheres expressed PAX6 and were capable of generating peripherin-negative neurons in our proof-of-principle experiment, highlighting that pure NCLC-populations are a prerequisite for successful sensory neuron differentiation. This indicates that the initial differentiation step from iPSCs to spheres and subsequently to NCLCs yields a system with the intrinsic potential to generate multiple cell types relevant for pain modelling. Importantly, these cell types could be derived from a single patient-specific sample, potentially providing a versatile platform for future personalized disease modelling.

The 2^nd^ checkpoint at d5, the time point of passaging during maturation, was used in our hands to transfer living progenitors to the appropriate culture format in the right density, and would allow viability measures, cell number or marker expression to be checked as well.

The 3^rd^, and last quality checkpoint at the end of differentiation is used to test for the presence of the right cell type. We checked for expression of sensory neuron and nociceptor associated marker by immunostaining and RNAseq and confirmed sensory neuron identity. Unfortunately, high throughput-based marker expression readouts of neurons are limited due to their tightly packed 3-dimensional clusters with dense neurite networks that prevent obtaining single cells e.g. for FACS, and interfere with high throughput 2-dimensional microscopy techniques available. This is even more complicated by a lack of identified markers for sensory neurons or reliable antibodies. However, developing high throughput automatised quality controls for neurons would be needed for translational applications in the future to efficiently validate differentiation success across cell lines right before experiments for disease modelling and drug screenings are conducted. For modelling neuropathic pain, sensory neurons, or more specifically nociceptors are needed that show expression of pain related genes, especially voltage gated sodium channels, and physiological electrophysiological function.

To mature sensory neuron progenitors, we selected a medium that contains physiological levels of glucose since diabetes often leads to neuropathies in patients. Commonly used differentiation media contain 25mM glucose which is highly diabetic. Adjusting for non-diabetic, more physiological conditions required formulating a new medium with reduced glucose content, for which we incorporated key components from established differentiation protocols to create a complete and supportive maturation environment. Under these conditions, we were able to reproduce a previously published SFN–associated phenotype characterized by increased excitability (6). The resulting neurons responded to capsaicin, AITC, menthol, and α/β-ATP, consistent with a nociceptor-like sensory neuron identity.

Despite these functional hallmarks, it is well established that iPSC-derived neurons remain developmentally immature and do not fully recapitulate their human *in vivo* counterparts. Sensory neurons generated *in vitro* typically remain small, multipolar, and exhibit transcriptional and functional profiles distinct from adult human DRG neurons (65–67). To address this, we tested several maturation strategies that have been proposed to enhance neuronal development, including supplementation with PGE_2_, retinoic acid, and dbcAMP, and extended the differentiation period up to six months while varying initial cell densities. None of these conditions markedly altered the morphology of the derived sensory neurons. Soma size remained small compared to human DRG neurons *in vitro*, reaching approximately 25 µm at day 70 and a median of 27 µm at day 168, whereas human DRG neurons typically exceed 28 µm in diameter (35). It is noteworthy that nociceptors are categorized as small diameter neurons in several species, including human. As expected, we did not observe pseudounipolar neurons by phase-contrast imaging since pseudounipolarity depend on glial support during development (67).

RNA-seq expression analysis revealed that, across all maturation conditions, the derived neurons expressed markers characteristic of neuronal, peripheral neuronal, sensory neuron, mechanoreceptor, proprioceptor, and nociceptor lineages. Among the tested maturation cues, PGE₂ frequently produced the strongest transcriptional effects. This is consistent with its biological role: PGE₂ is a major component of Schwann-cell–conditioned medium and has been shown to increase sodium channel expression in embryonic mouse DRG neurons, *in vitro* (39). Similarly, we observe increased Nav1.8 and Nav1.9 expression, both reported to show low expression in iPS-derived sensory neurons (1, 65). In adult systems, PGE_2_ functions as a potent inflammatory mediator that modulates Nav channel activity and sensitizes nociceptors (68, 69). To minimize acute inflammatory effects, PGE₂ supplementation in our protocol was restricted to days 21–49. Thus, the transcriptional changes detected by RNAseq likely reflect long-term maturation effects rather than transient inflammatory activation. Developmental markers associated with sensory neuron lineage specification, including *PRDM12* (pan-sensory), *cMET* (peptidergic lineage), and *RUNX1* (non-peptidergic lineage), showed lower expression levels in PGE₂-treated cultures compared to untreated controls. At the same time, PGE₂-treated samples exhibited increased expression of markers associated with mature sensory neurons and nociceptors. This pattern may indicate a shift from early developmental programs toward a more mature transcriptional state in response to PGE₂. We detected nociceptor subtype markers under all maturation conditions. Human nociceptors predominantly display a peptidergic phenotype, characterized by expression of *TAC1* (substance P) and *CALCA* (CGRP) (70). Both genes showed their highest expression levels in long-term differentiated cultures, suggesting that acquisition of peptidergic identity requires extended maturation time. In contrast, *SST*—encoding somatostatin and associated with silent nociceptors (43)—showed the highest expression in PGE₂-treated samples. These nociceptor subclass was also described to express the voltage gated sodium channels Nav1.8 and Nav1.9 and *TAC1* (43). This finding implies that PGE₂ may preferentially promote maturation of specific nociceptor subtypes. Taken together, these results suggest that prolonged differentiation and PGE_2_ supplementation influence distinct aspects of sensory neuron maturation. A combination of extended culture time and targeted modulators such as PGE₂ may therefore represent a promising strategy to enhance human-like nociceptor identity. Further functional and proteomic analyses will be required to determine the optimal maturation conditions.

To evaluate the functional impact of the different maturation conditions, we employed MEA recordings, which offer a medium-throughput platform for assessing neuronal excitability. Across all conditions, we did not detect differences in spontaneous activity at day 70, and despite the transcriptional effects of PGE₂, no PGE₂-specific changes in MEA parameters were observed. Instead, MEA features changed primarily as a function of maturation time, characterized by an initial increase in spontaneous activity followed by a decline during prolonged culture. In the absence of human DRG recordings as a direct reference, MEA measurements from patients provide an important functional readout to assess the physiological relevance our model. In this context, neurons derived from patients with SFN displayed increased spontaneous activity, consistent with previously reported hyperexcitability phenotypes. This indicates that several of our maturation conditions (untreated, PGE₂, retinoic acid) are capable to capture disease-relevant functional alterations. The interpretation of the IEM cell line and its CRIPSR repaired counterpart over all maturation conditions is more challenging. Depending on the maturation condition, the CRISPR line showed partial convergence either toward the patient phenotype or toward the healthy control. It remains unclear whether CRISPR-based correction restores a “healthy-like” electrophysiological baseline or whether genetic background effects and differentiation variability influence the expected rescue. This highlights open questions regarding the influence of genetic background, maturation state, and culture variability in MEA-based phenotyping. Overall, while MEA recordings provide valuable first-level functional insights, they cannot yet – at least based on our data - determine which maturation condition yields the most physiologically relevant nociceptor phenotype.

One additional limitation arises from the inherent variability of MEA recordings itself. Neurons settle stochastically on the chip, resulting in electrodes that may capture only neurites, a single neuron, multiple neurons, or no cells at all. This spatial randomness contributes substantially to variability in MEA Data and cannot be fully controlled experimentally. Increasing the number of controls, patient lines, and biological replicates will help mitigate this variability, but it is unlikely to eliminate it entirely. Consequently, direct comparisons between different disease entities remain a challenge, as absolute activity levels are strongly influenced by electrode occupancy. Still, MEA recordings remain highly suitable for drug screening, since normalization within each well allows robust detection of relative drug-induced changes independent of baseline variability.

Taken together, among the tested conditions, seeding 30,000 cells per MEA combined with PGE₂ maturation until day 49 and analysis at day 70 currently provides the best compromise between nociceptor marker expression, functional maturation, and detectability of patient-specific electrophysiological phenotypes. Nevertheless, this conclusion must be considered preliminary, as PGE₂ is an inflammatory mediator and long-term effects cannot be fully excluded. Further systematic evaluation across larger cohorts and complementary functional assays will be required to define the optimal maturation condition to derive hDRG-like neurons for patient-specific disease modelling and drug screening in the future.

## 5. Conclusion

In summary, we present a robust, scalable and standardized protocol for the differentiation of multiple iPSC lines with diverse genetic backgrounds into sensory neurons. The workflow relies on simple, modular steps and therefore provides a promising basis for future translational adaptation, including cost reduction, xeno-free implementation, and potential automation. Nevertheless, the dependence on lentiviral NGN1 induction may limit applicability in laboratories without S2 facilities and poses challenges for clinical-grade manufacturing, highlighting the need for alternative strategies to achieve controlled NGN1 expression. Despite these remaining limitations, our study establishes a solid entry point for future refinement for downstream applications ranging from patient-specific disease modelling and drug screening to functional genomics and drug discovery.

We deconstructed the NGN1-based sensory neuron differentiation protocol into its essential components and systematically validated each step across multiple independent iPSC lines. This refinement resulted in a robust and scalable workflow that enables the differentiation of diverse patient iPSCs in medium-throughput settings while providing some flexibility to compensate for the inherent heterogeneity of iPSCs. Owing to its modular structure and integrated checkpoints, the protocol can be gradually adapted to meet future translational requirements, including cost reduction, xeno-free implementation, increased throughput and automated processing. Although we cannot evaluate entirely if our iPSC-derived sensory neurons recapitulate the *in vivo* state of human DRG neurons, well-defined and reproducible protocols such as ours are essential building blocks for advancing translational applications. They provide a reliable starting point for further optimization and will facilitate the development of improved cellular models for human sensory neuron biology and disease.

## 7. Declarations

### 7.1 Ethics Approval and Consent to Participate

Participants donated blood or fibroblasts for stem cell reprogramming following informed consent. The procedure was approved by the local ethics committee at the Medical Faculty of the RWTH Aachen (EK 243/18), as published before for (Fib13_30, (6)), or as stated by courielle (Fib7_1).

### 7.2 Consent for Publication

Not applicable.

### 7.3 Availability of Data and Materials

The data generated during the current study are available from the corresponding author.

### 7.4 Competing Interests

ALa receives counselling fees from Grünenthal, Orion and Netri.

MFD reports grants from Pfizer Pharmaceuticals (ASPIRE 2018); consulting fees from Pfizer, Alnylam, Akcea, AstraZeneca, Amicus Therapeutics, Applied Therapeutics, and Sobi; payment or honoraria from Pfizer, Alnylam, Akcea, Amicus Therapeutics, AstraZeneca, and Sobi; support for attending meetings and/or travel from Pfizer, Alnylam, Akcea, AstraZeneca, Sobi, and Amicus Therapeutics; participation on advisory boards for Pfizer, Alnylam, Akcea, AstraZeneca, and Purpose Pharma; and a leadership role in the European CMT Research Association (ERCA).

AKK is currently employed by Grünenthal.

RR received fees as a speaker or for counselling services from Aristo Pharma, Avextra, cannamedical, Grünenthal, Tilray, Vayamed.

### 7.5 Funding

This work was funded by the Deutsche Forschungsgemeinschaft (German Research Foundation), 363055819/GRK2415 Mechanobiology of 3D epithelial tissues (ME3T) to ALa; 368482240/GRK2416, MultiSenses-MultiScales to ALa; and LA 2740/6-1 to ALa, by the BMBF consortium “Bio2Treat” (German Federal Ministry of Education and Research / Bundesministerium für Bildung und Forschung, BMBF, “Chronische Schmerzen-Innovative medizintechnische Lösungen zur Verbesserung von Prävention, Diagnostik und Therapie”, contract number 13GW0334B to ALa and RR. ALa was supported by a grant from the Interdisciplinary Center for Clinical Research within the Faculty of Medicine at the RWTH Aachen University (IZKF TN1-1/IA 532001).

### 7.6 Authors’ Contributions

AN planned and performed experiments, and analysed the data, interpreted the results and wrote the manuscript.

MM performed the seeding-density and Ara-C treatment time point experiments and contributed data to viral load experiments.

VD performed the soma-size quantification

CG conducted lentiviral production and performed initial experiments to reproduce the original protocol.

LM analysed MEA-Data for cell responses to ion channel agonists.

MFD, NWMB and RR selected and examined patients.

FK, LB, and IK carried out the RNAseq runs and contributed to parts of the analysis.

AE performed and analyzed live-cell calcium imaging experiments.

AKK performed the reprogramming and generation of iPSCs from the IEM patient and crispr-repaired clones.

SuS reprogrammed and established iPSC-lines with B2T-ID.

IK and MZ conceived the study

ALa conceived and supervised the study, interpreted the data, and allocated the funding.

All authors participated in reviewing the manuscript.

## 7.7 Acknowledgements

We would like to express our gratitude to the participants for taking part in our research.

We thank Prof. Wolfgang Wagner (Institute for Stem Cell Biology, RWTH Aachen University Medical School, Aachen, Germany; Helmholtz Institute for Biomedical Engineering, RWTH Aachen University Medical School, Aachen, Germany) for providing the iPSC-line “B1”. Plasmids for lentiviral production were kindly provided by Dr. Katrin Schrenk-Siemens (Institute of Pharmacology, Heidelberg University, Heidelberg, Germany) and through Addgene (71). The mouse Schwann cell line MSC-80 (72) was kindly provided by Dr. Burkhard Gess (Clinics for Neurology, Uniklinik RWTH Aachen, Germany).

This work was supported by the Flow Cytometry Facility of the Interdisciplinary Center for Clinical Research (IZKF) within the Faculty of Medicine at RWTH Aachen University.

AI assistance (ChatGPT) was used to generate initial code templates for data processing and analysis. Final code reflects substantial manual revision. We used Microsoft Copilot (version accessed in March 2026) to support text revision. All outputs were reviewed, verified, and adapted by the authors.

## 6. List of Abbreviations

Ara-C: Cytarabine
BDNF: Brain-Derived Neurotrophic Factor
cAMP: Cyclic Adenosine Monophosphate
CGRP: Calcitonin Gene-Related Peptide
CPM: Counts Per Million
CT: Cycle Threshold
d: day of differentiation
DAPI: 4’,6-Diamidino-2-phenylindole
DRG: Dorsal Root Ganglion
EGF: Epidermal Growth Factor
eGFP: Enhanced Green Fluorescent Protein
EtOH: Ethanol
FACS: Fluorescence-Activated Cell Sorting
FGF2: Fibroblast Growth Factor 2
Fura-2 AM: Acetoxymethyl Ester of Fura-2
GAPDH: Glyceraldehyde-3-phosphate Dehydrogenase
GDNF: Glial Cell Line-Derived Neurotrophic Factor
GSEA: Gene Set Enrichment Analysis
IBI: Inter-Burst-Interval
iPSC: Induced Pluripotent Stem Cell
IQR: Interquartile Range
ISI: Inter-Spike Interval
LN521: Laminin-521
MEA: Multi-Electrode Array
MFR: Mean Firing Rate
MOI: Multiplicity of Infection
NGF: Nerve Growth Factor
NGN1: Neurogenin-1
NCLC: Neural Crest-Like Cell
PCA: Principal Component Analysis
PFA: Paraformaldehyde
PGE₂: Prostaglandin E₂
RA: Retinoic Acid
ROI: Region of Interest
rtTA: Reverse Tetracycline-Controlled Transactivator
TRP: Transient Receptor Potential
unt.: untreated

## Notes

### Competing Interest Statement

ALa receives counselling fees from Gruenenthal, Orion and Netri. MFD reports grants from Pfizer Pharmaceuticals (ASPIRE 2018); consulting fees from Pfizer, Alnylam, Akcea, AstraZeneca, Amicus Therapeutics, Applied Therapeutics, and Sobi; payment or honoraria from Pfizer, Alnylam, Akcea, Amicus Therapeutics, AstraZeneca, and Sobi; support for attending meetings and/or travel from Pfizer, Alnylam, Akcea, AstraZeneca, Sobi, and Amicus Therapeutics; participation on advisory boards for Pfizer, Alnylam, Akcea, AstraZeneca, and Purpose Pharma; and a leadership role in the European CMT Research Association (ERCA). AKK is currently employed by Gruenenthal. RR received fees as a speaker or for counselling services from Aristo Pharma, Avextra, cannamedical, Grünenthal, Tilray, Vayamed.

## References

1. Schrenk-Siemens K, Pohle J, Rostock C, Abd El Hay M, Lam RM, Szczot M, et al. Human Stem Cell-Derived TRPV1-Positive Sensory Neurons: A New Tool to Study Mechanisms of Sensitization. Cells. 2022;11(18).

2. Soliman N, Kersebaum D, Lawn T, Sachau J, Sendel M, Vollert J. Improving neuropathic pain treatment - by rigorous stratification from bench to bedside. Journal of neurochemistry. 2024;168(11):3699–714.

3. Jensen TS, Baron R, Haanpää M, Kalso E, Loeser JD, Rice ASC, et al. A new definition of neuropathic pain. Pain. 2011;152(10):2204–5.

4. Scholz J, Finnerup NB, Attal N, Aziz Q, Baron R, Bennett MI, et al. The IASP classification of chronic pain for ICD-11: chronic neuropathic pain. Pain. 2019;160(1):53–9.

5. Baskozos G, Hébert HL, Pascal MM, Themistocleous AC, Macfarlane GJ, Wynick D, et al. Epidemiology of neuropathic pain: an analysis of prevalence and associated factors in UK Biobank. Pain reports. 2023;8(2):e1066.

6. Namer B, Schmidt D, Eberhardt E, Maroni M, Dorfmeister E, Kleggetveit IP, et al. Pain relief in a neuropathy patient by lacosamide: Proof of principle of clinical translation from patient-specific iPS cell-derived nociceptors. EBioMedicine. 2019;39:401–8.

7. McDermott LA, Weir GA, Themistocleous AC, Segerdahl AR, Blesneac J, Baskozos G, et al. Defining the Functional Role of Na V 1.7 in Human Nociception. 2018.

8. Cao L, Aoibhinn McDonnell, Anja Nitzsche, Aristos Alexandrou, Pierre-Philippe Saintot, Alexandre J.C. Loucif, et al. Pharmacological reversal of a pain phenotype in iPSC-derived sensory neurons and patients with inherited erythromelalgia. 2016.

9. Kalia AK, Rösseler C, Granja-Vazquez R, Ahmad A, Pancrazio JJ, Neureiter A, et al. How to differentiate induced pluripotent stem cells into sensory neurons for disease modelling: a functional assessment. Stem Cell Res Ther. 2024;15(1):99.

10. Volpato V, Webber C. Addressing variability in iPSC-derived models of human disease: guidelines to promote reproducibility. Disease models & mechanisms. 2020;13(1).

11. Saito-Diaz K, Street JR, Ulrichs H, Zeltner N. Derivation of Peripheral Nociceptive, Mechanoreceptive, and Proprioceptive Sensory Neurons from the same Culture of Human Pluripotent Stem Cells. Stem Cell Reports. 2021;16(3):446–57.

12. Cantor EL, Shen F, Jiang G, Tan Z, Cunningham GM, Wu X, et al. Passage number affects differentiation of sensory neurons from human induced pluripotent stem cells. Sci Rep. 2022;12(1):15869.

13. Haubenreich C, Lenz M, Schuppert A, Peitz M, Koch P, Zenke M, et al. Epigenetic and Transcriptional Shifts in Human Neural Stem Cells after Reprogramming into Induced Pluripotent Stem Cells and Subsequent Redifferentiation. International journal of molecular sciences. 2024;25(6).

14. Kim K, Doi A, Wen B, Ng K, Zhao R, Cahan P, et al. Epigenetic memory in induced pluripotent stem cells. Nature. 2010;467(7313):285–90.

15. Nasu A, Ikeya M, Yamamoto T, Watanabe A, Jin Y, Matsumoto Y, et al. Genetically matched human iPS cells reveal that propensity for cartilage and bone differentiation differs with clones, not cell type of origin. PLoS One. 2013;8(1):e53771.

16. Schwartzentruber J, Foskolou S, Kilpinen H, Rodrigues J, Alasoo K, Knights AJ, et al. Molecular and functional variation in iPSC-derived sensory neurons. Nat Genet. 2018;50(1):54–61.

17. Galiakberova A, Ivanov S, Golov A, Artyuhov A, Zolkin A, Kondratyev N, et al. Transcriptomic profiling of neural cultures from the KYOU iPSC line via alternative differentiation protocols. Frontiers in molecular neuroscience. 2025;18:1661986.

18. Zhang Y, Pak C, Han Y, Ahlenius H, Zhang Z, Chanda S, et al. Rapid single-step induction of functional neurons from human pluripotent stem cells. Neuron. 2013;78(5):785–98.

19. Servetti M, Caramia M, Parodi G, Loiacono F, Nano E, Biddau G, et al. Optimization of Transcription Factor-Driven Neuronal Differentiation from Human Induced Pluripotent Stem Cells for Disease Modelling and Drug Screening. Stem cell reviews and reports. 2025;21(3):816–33.

20. El Wazan L, Urrutia-Cabrera D, Wong RC. Using transcription factors for direct reprogramming of neurons in vitro. World journal of stem cells. 2019;11(7):431–44.

21. Ma Q, Fode C, Guillemot F, Anderson DJ. Neurogenin1 and neurogenin2 control two distinct waves of neurogenesis in developing dorsal root ganglia. Genes & development. 1999;13(13):1717–28.

22. Schindelin J, Arganda-Carreras I, Frise E, Kaynig V, Longair M, Pietzsch T, et al. Fiji: an open-source platform for biological-image analysis. Nature Methods. 2012;9(7):676–82.

23. Nguyen MQ, von Buchholtz LJ, Reker AN, Ryba NJ, Davidson S. Single-nucleus transcriptomic analysis of human dorsal root ganglion neurons. Elife. 2021;10.

24. Lai X, Liu J, Zou Z, Wang Y, Wang Y, Liu X, et al. SOX10 ablation severely impairs the generation of postmigratory neural crest from human pluripotent stem cells. Cell death & disease. 2021;12(9):814.

25. Thomas R, Menon V, Mani R, Pruszak J. Glycan Epitope and Integrin Expression Dynamics Characterize Neural Crest Epithelial-to-Mesenchymal Transition (EMT) in Human Pluripotent Stem Cell Differentiation. Stem cell reviews and reports. 2022;18(8):2952–65.

26. Lee G, Kim H, Elkabetz Y, Al Shamy G, Panagiotakos G, Barberi T, et al. Isolation and directed differentiation of neural crest stem cells derived from human embryonic stem cells. Nat Biotechnol. 2007;25(12):1468–75.

27. Sandireddy R, Yerra VG, Areti A, Komirishetty P, Kumar A. Neuroinflammation and oxidative stress in diabetic neuropathy: futuristic strategies based on these targets. International journal of endocrinology. 2014;2014:674987.

28. Liu Y, Balaji R, de Toledo MAS, Ernst S, Hautvast P, Kesdoğan AB, et al. The pain target Na(V)1.7 is expressed late during human iPS cell differentiation into sensory neurons as determined in high-resolution imaging. Pflugers Archiv: European journal of physiology. 2024;476(6):975–92.

29. Stadler T, O’Reilly AO, Lampert A. Erythromelalgia mutation Q875E Stabilizes the activated state of sodium channel Nav1.7. The Journal of biological chemistry. 2015;290(10):6316–25.

30. Kesdoğan AB, Neureiter A, Gaebler AJ, Kalia AK, Körner J, Lampert A. Analgesic effect of Botulinum toxin in neuropathic pain is sodium channel independent. Neuropharmacology. 2024;253:109967.

31. Grant S. Ara-C: cellular and molecular pharmacology. Advances in cancer research. 1998;72:197–233.

32. Bass PD, Gubler DA, Judd TC, Williams RM. Mitomycinoid alkaloids: mechanism of action, biosynthesis, total syntheses, and synthetic approaches. Chem Rev. 2013;113(8):6816–63.

33. Young GT, Gutteridge A, Fox H, Wilbrey AL, Cao L, Cho LT, et al. Characterizing human stem cell-derived sensory neurons at the single-cell level reveals their ion channel expression and utility in pain research. Mol Ther. 2014;22(8):1530–43.

34. Hiller BM, Marmion DJ, Gross RM, Thompson CA, Chavez CA, Brundin P, et al. Mitomycin-C treatment during differentiation of induced pluripotent stem cell-derived dopamine neurons reduces proliferation without compromising survival or function in vivo. Stem Cells Translational Medicine. 2020;10(2):278–90.

35. Davidson S, Copits BA, Zhang J, Page G, Ghetti A, Gereau RWt. Human sensory neurons: Membrane properties and sensitization by inflammatory mediators. Pain. 2014;155(9):1861–70.

36. Corcoran J, Shroot B, Pizzey J, Maden M. The role of retinoic acid receptors in neurite outgrowth from different populations of embryonic mouse dorsal root ganglia. Journal of Cell Science. 2000;113(14):2567–74.

37. Desiderio S, Vermeiren S, Van Campenhout C, Kricha S, Malki E, Richts S, et al. Prdm12 Directs Nociceptive Sensory Neuron Development by Regulating the Expression of the NGF Receptor TrkA. Cell Reports. 2019;26(13):3522–36.e5.

38. Maden M. Retinoic acid in the development, regeneration and maintenance of the nervous system. Nature Reviews Neuroscience. 2007;8(10):755–65.

39. Kantarci H, Elvira PD, Thottumkara AP, O’Connell EM, Iyer M, Donovan LJ, et al. Schwann cell-secreted PGE(2) promotes sensory neuron excitability during development. Cell. 2024;187(17):4690–712.e30.

40. Rydel RE, Greene LA. cAMP analogs promote survival and neurite outgrowth in cultures of rat sympathetic and sensory neurons independently of nerve growth factor. Proc Natl Acad Sci U S A. 1988;85(4):1257–61.

41. Haberberger RV, Barry C, Matusica D. Immortalized Dorsal Root Ganglion Neuron Cell Lines. Frontiers in cellular neuroscience. 2020; Volume 14 - 2020.

42. Cai D, Qiu J, Cao Z, McAtee M, Bregman BS, Filbin MT. Neuronal cyclic AMP controls the developmental loss in ability of axons to regenerate. J Neurosci. 2001;21(13):4731–9.

43. Körner J, Howard D, Solinski HJ, Mancilla Moreno M, Haag N, Fiebig A, et al. Molecular architecture of human dermal sleeping nociceptors. Cell. 2026;189(6):1820–35.e22.

44. Luhmann HJ, Sinning A, Yang J-W, Reyes-Puerta V, Stüttgen MC, Kirischuk S, et al. Spontaneous Neuronal Activity in Developing Neocortical Networks: From Single Cells to Large-Scale Interactions. Frontiers in Neural Circuits. 2016; Volume 10 - 2016.

45. van den Braak NWM, Kuehs S, Peschke GZ, Namer B, Lischka A, Eggermann K, et al. Altered NaV1.9 channel activity in two Tyr66Ser variant carriers with small fiber dysfunction. The Journal of general physiology. 2025;157(6).

46. Dohrn MF, Dumke C, Hornemann T, Nikolin S, Lampert A, Espenkott V, et al. Deoxy-sphingolipids, oxidative stress, and vitamin C correlate with qualitative and quantitative patterns of small fiber dysfunction and degeneration. Pain. 2022;163(9):1800–11.

47. Skeik N, Rooke TW, Davis MD, Davis DM, Kalsi H, Kurth I, et al. Severe case and literature review of primary erythromelalgia: novel SCN9A gene mutation. Vascular medicine (London, England). 2012;17(1):44–9.

48. Bajpai R, Chen DA, Rada-Iglesias A, Zhang J, Xiong Y, Helms J, et al. CHD7 cooperates with PBAF to control multipotent neural crest formation. Nature. 2010;463(7283):958–62.

49. Burns JC, Friedmann T, Driever W, Burrascano M, Yee JK. Vesicular stomatitis virus G glycoprotein pseudotyped retroviral vectors: concentration to very high titer and efficient gene transfer into mammalian and nonmammalian cells. Proc Natl Acad Sci U S A. 1993;90(17):8033–7.

50. Hoffmann M, Wu YJ, Gerber M, Berger-Rentsch M, Heimrich B, Schwemmle M, et al. Fusion-active glycoprotein G mediates the cytotoxicity of vesicular stomatitis virus M mutants lacking host shut-off activity. The Journal of general virology. 2010;91(Pt 11):2782–93.

51. Feliciano D, Ott CM, Espinosa-Medina I, Weigel AV, Benedetti L, Milano KM, et al. YAP1 nuclear efflux and transcriptional reprograming follow membrane diminution upon VSV-G-induced cell fusion. Nat Commun. 2021;12(1):4502.

52. Pichlmair A, Diebold SS, Gschmeissner S, Takeuchi Y, Ikeda Y, Collins MK, et al. Tubulovesicular structures within vesicular stomatitis virus G protein-pseudotyped lentiviral vector preparations carry DNA and stimulate antiviral responses via Toll-like receptor 9. Journal of virology. 2007;81(2):539–47.

53. Ansari AM, Ahmed AK, Matsangos AE, Lay F, Born LJ, Marti G, et al. Cellular GFP Toxicity and Immunogenicity: Potential Confounders in in Vivo Cell Tracking Experiments. Stem cell reviews and reports. 2016;12(5):553–9.

54. Toran PT, Wohlfahrt M, Foye J, Kiem HP, Wojchowski DM. Assessment and streamlined preparation of low-cytotoxicity lentiviral vectors for mobilized human hematopoietic stem cell transduction. Experimental hematology. 2020;86:28–42.e3.

55. Zhang B, Metharom P, Jullie H, Ellem KA, Cleghorn G, West MJ, et al. The significance of controlled conditions in lentiviral vector titration and in the use of multiplicity of infection (MOI) for predicting gene transfer events. Genetic vaccines and therapy. 2004;2(1):6.

56. Mu T, Qin Y, Liu B, He X, Liao Y, Sun J, et al. In Vitro Neural Differentiation of Bone Marrow Mesenchymal Stem Cells Carrying the FTH1 Reporter Gene and Detection with MRI. BioMed research international. 2018;2018:1978602.

57. Neureiter A, Eberhardt E, Lampert A. Differentiation of iPS-Cells into Peripheral Sensory Neurons. Methods Mol Biol. 2022;2429:175–88.

58. Eberhardt E, Namer B, Neureiter A, Körner J, Jørum E, Kurth I, et al. Spontaneous activity in pain patient stem cell-derived sensory neurons arises from one functional subclass. Pain. 2025.

59. Iwasaki H, Huang P, Keating MJ, Plunkett W. Differential Incorporation of Ara-C, Gemcitabine, and Fludarabine Into Replicating and Repairing DNA in Proliferating Human Leukemia Cells. Blood. 1997;90(1):270–8.

60. Wallace TL, Johnson EM, Jr. Cytosine arabinoside kills postmitotic neurons: evidence that deoxycytidine may have a role in neuronal survival that is independent of DNA synthesis. J Neurosci. 1989;9(1):115–24.

61. Chambers SM, Mica Y, Lee G, Studer L, Tomishima MJ. Dual-SMAD Inhibition/WNT Activation-Based Methods to Induce Neural Crest and Derivatives from Human Pluripotent Stem Cells. Methods Mol Biol. 2016;1307:329–43.

62. Gozlan O, Sprinzak D. Notch signaling in development and homeostasis. Development (Cambridge, England). 2023;150(4).

63. Cornell RA, Eisen JS. Delta/Notch signaling promotes formation of zebrafish neural crest by repressing Neurogenin 1 function. Development (Cambridge, England). 2002;129(11):2639–48.

64. Elkabetz Y, Panagiotakos G, Al Shamy G, Socci ND, Tabar V, Studer L. Human ES cell-derived neural rosettes reveal a functionally distinct early neural stem cell stage. Genes & development. 2008;22(2):152–65.

65. Deng T, Jovanovic VM, Tristan CA, Weber C, Chu P-H, Inman J, et al. Scalable generation of sensory neurons from human pluripotent stem cells. Stem Cell Reports. 2023;18(4):1030–47.

66. Zurek NA, Ehsanian R, Goins AE, Adams IM, Petersen T, Goyal S, et al. Electrophysiological Analyses of Human Dorsal Root Ganglia and Human Induced Pluripotent Stem Cell-derived Sensory Neurons From Male and Female Donors. The Journal of Pain. 2024;25(6).

67. LeBlang CJ, Pazyra-Murphy MF, Silagi ES, Dasgupta S, Tsolias M, Miller T, et al. Satellite glial contact enhances differentiation and maturation of human iPSC-derived sensory neurons. Stem Cell Reports. 2025;20(10).

68. Maingret F, Coste B, Padilla F, Clerc N, Crest M, Korogod SM, et al. Inflammatory mediators increase Nav1.9 current and excitability in nociceptors through a coincident detection mechanism. The Journal of general physiology. 2008;131(3):211–25.

69. Villarreal CF, Sachs D, Funez MI, Parada CA, de Queiroz Cunha F, Ferreira SH. The peripheral pro-nociceptive state induced by repetitive inflammatory stimuli involves continuous activation of protein kinase A and protein kinase C epsilon and its Na(V)1.8 sodium channel functional regulation in the primary sensory neuron. Biochemical pharmacology. 2009;77(5):867–77.

70. Bhuiyan SA, Nagi SS, Sankaranarayanan I, Semizoglou E, Usoskin D, Yang L, et al. A Reference Atlas of the Human Dorsal Root Ganglion. bioRxiv. 2025.

71. Dull T, Zufferey R, Kelly M, Mandel RJ, Nguyen M, Trono D, et al. A third-generation lentivirus vector with a conditional packaging system. Journal of virology. 1998;72(11):8463–71.

72. Boutry JM, Hauw JJ, Gansmüller A, Di-Bert N, Pouchelet M, Baron-Van Evercooren A. Establishment and characterization of a mouse Schwann cell line which produces myelin in vivo. Journal of neuroscience research. 1992;32(1):15–26

