## Supplementary data for "A Framework for NGN1-Induced Sensory Neuron Differentiation for Disease Modelling and Drug Screening"

### 9. Supplementary Information

**SI Table 1 previously published medium composition and time frame for differentiation of sensory neurons.**

Media composition and the maximum time of differentiation was retrieved from 35 different publications reporting the differentiation of human pluripotent stem cells to sensory neurons between 2015 and 2025.

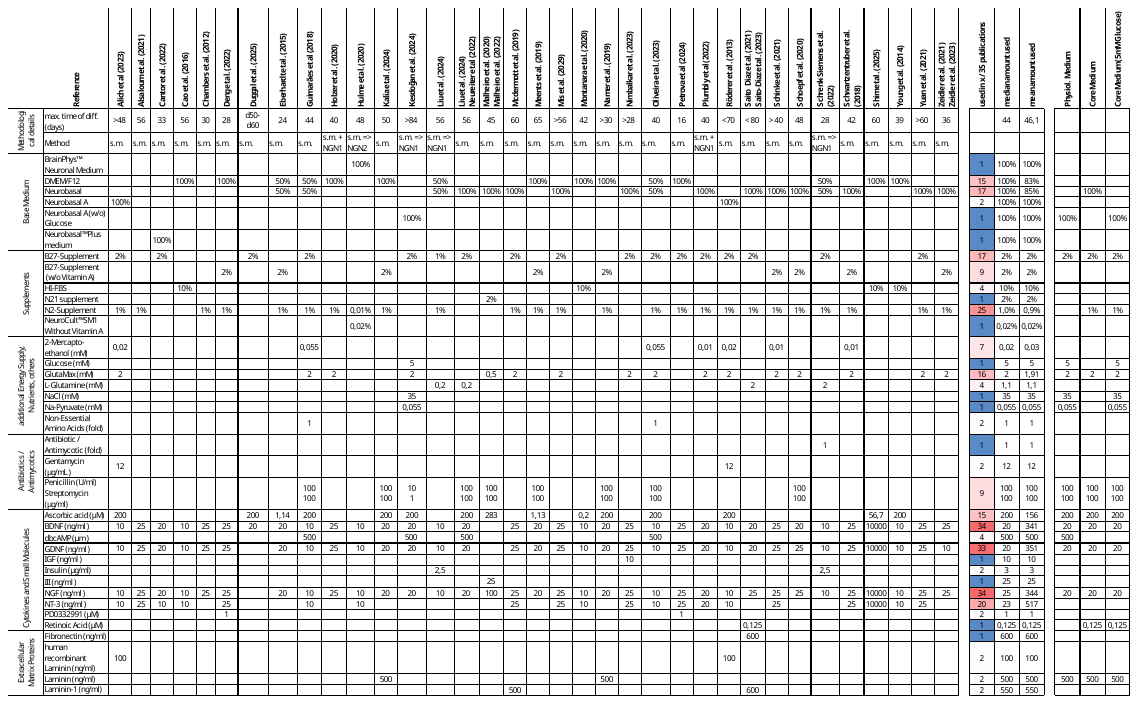

**SI Table 2 iPSCs used in the study.**

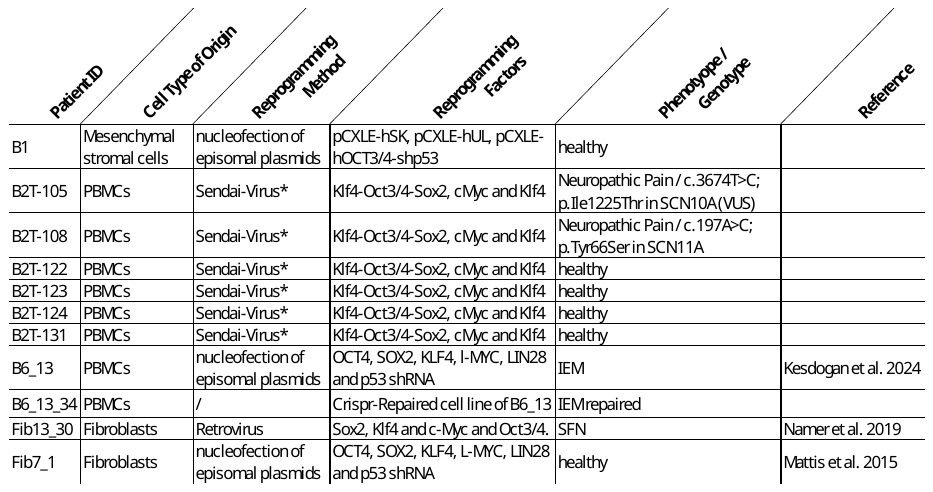

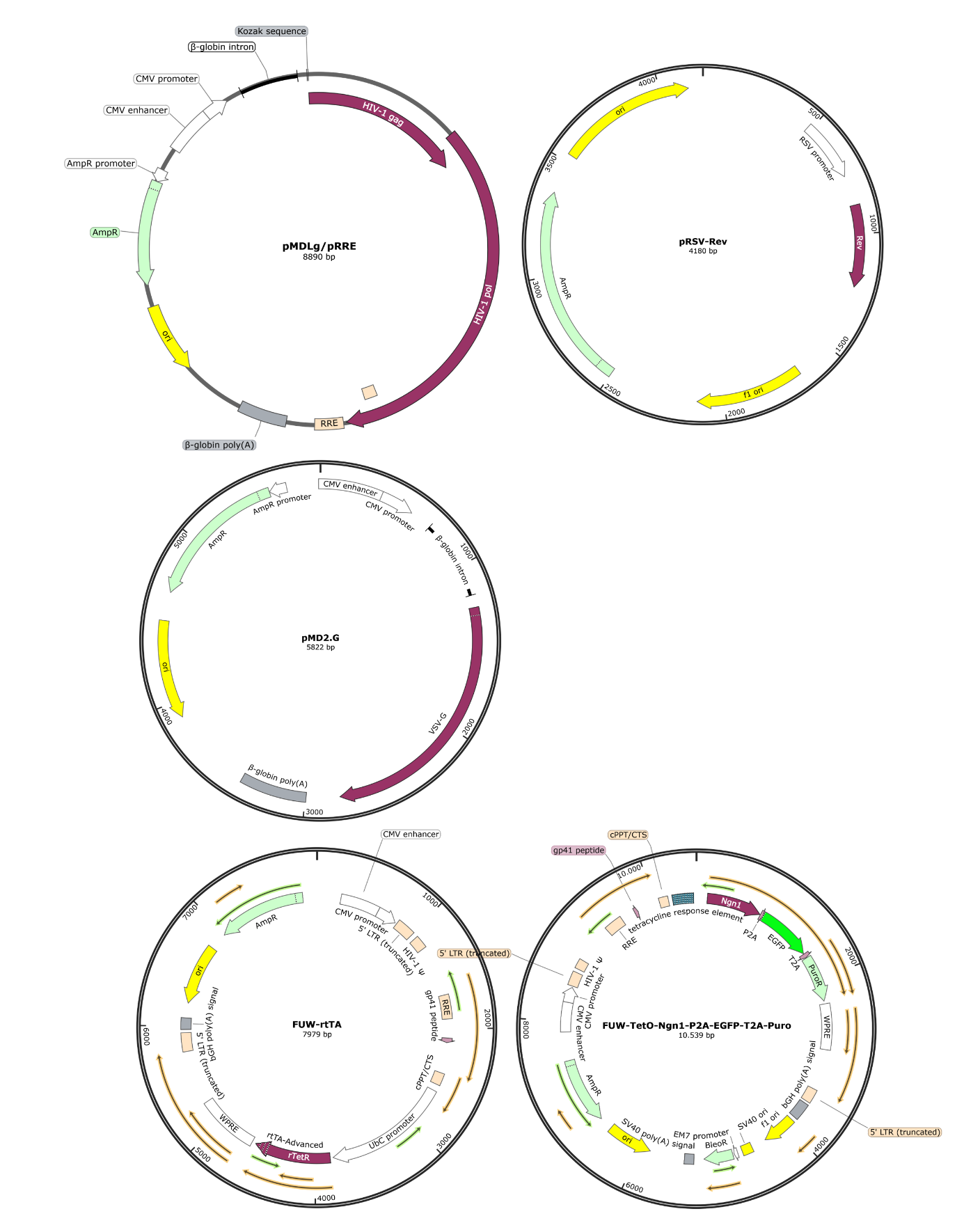

**SI Figure 1 Lentiviral Plasmids for production of lentiviral particles for the NGN1-driven differentiation.**

Initially, all plasmids were kindly provided by Katrin Schrenk Siemens (Institute for Pharmacology, University hospital Heidelberg). Subsequently, plasmid pMDLg/pRRE was replaced by the same plasmid distributed by Addgene (#12251, pMDLg/pRRE was a gift from Didier Trono (Addgene plasmid # 12251 ; http://n2t.net/addgene:12251 ; RRID:Addgene_12251).

**SI Table 3 Antibodies**

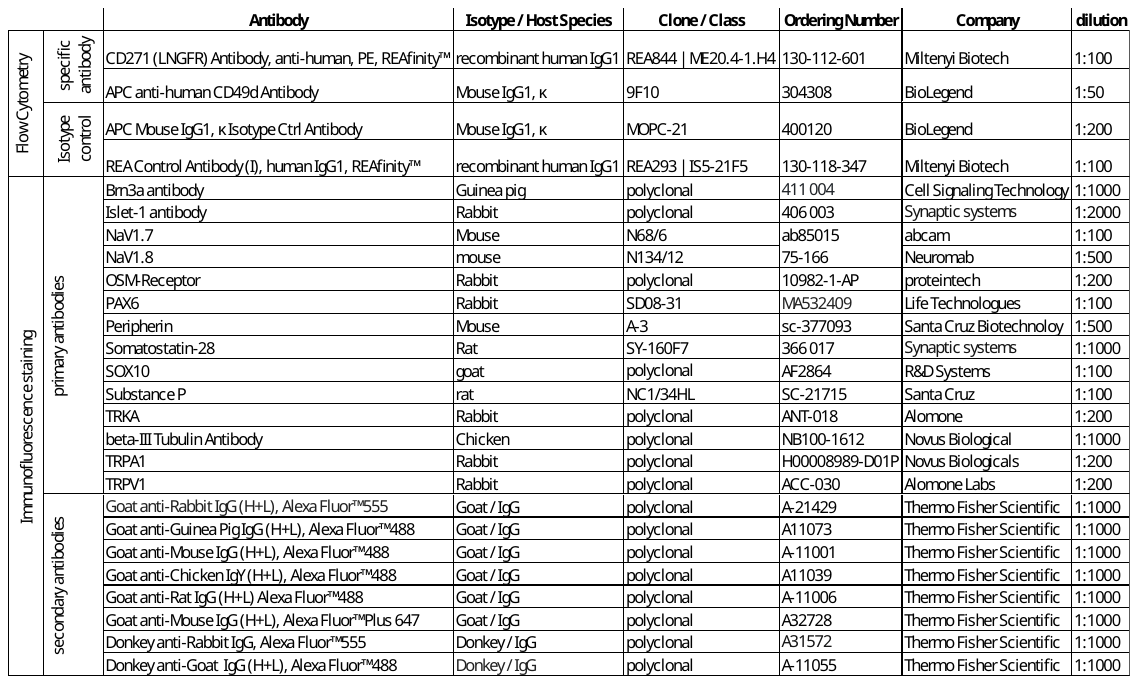

**SI Table 4 qPCR-Primer**

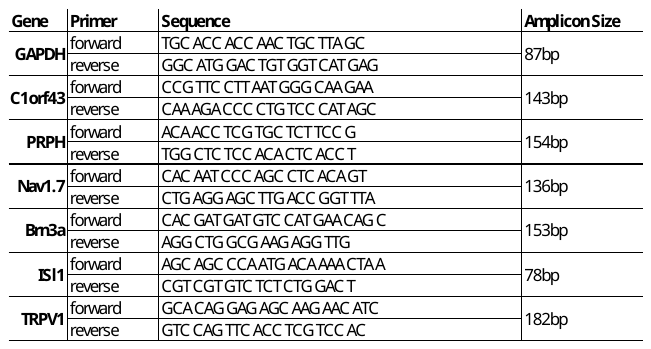

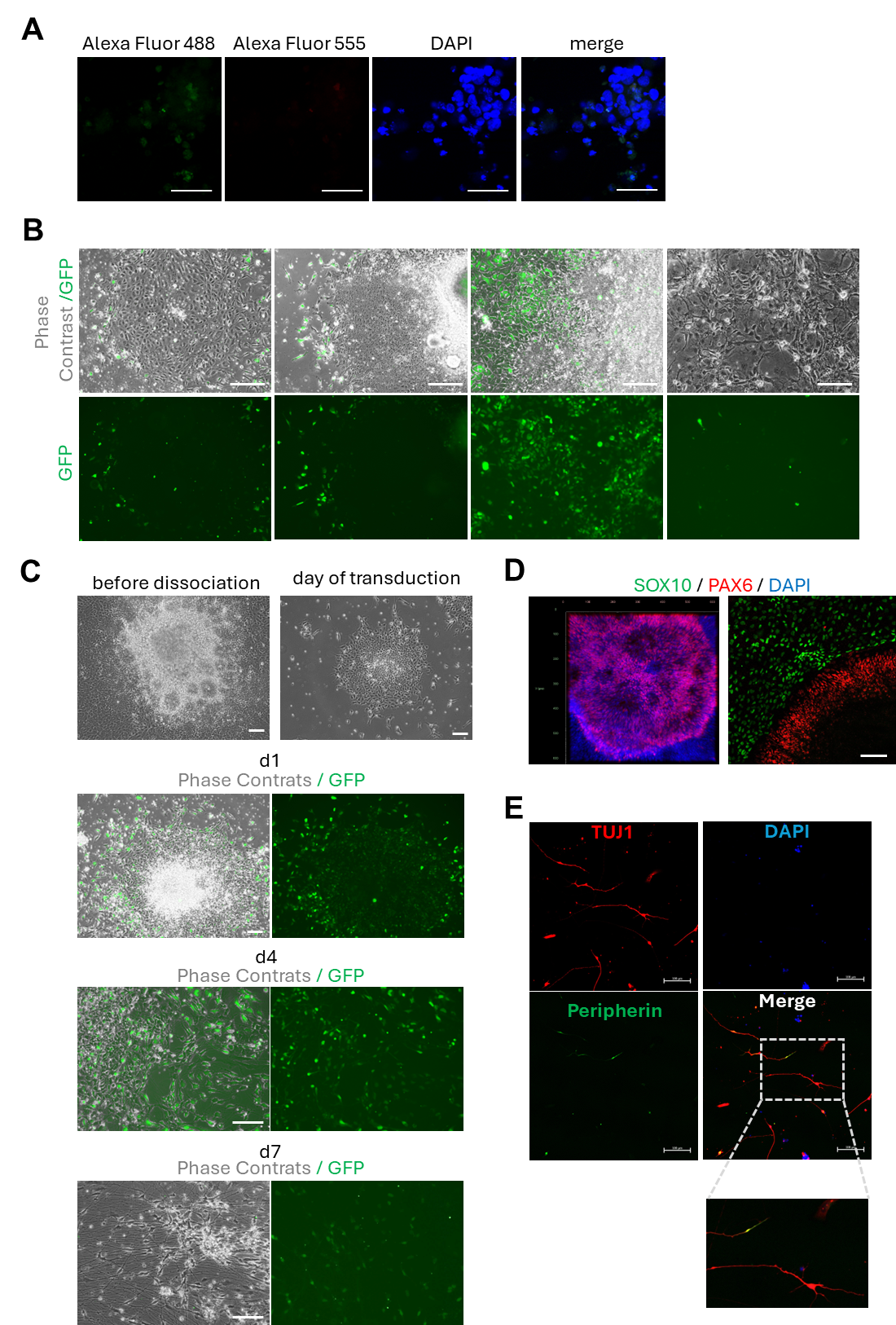

**SI Figure 2 Differentiation of multiple cell identities during NCLC-development.**

(A) Secondary antibody control for all immunostainings shown in Figure 1. Scale bar: 50 µm. (B) When attached to geltrex coated dishes, spheres contain heterogeneous cell populations that can be transduced with NGN1‑ and rtTA‑lentiviruses, albeit with low efficiency. Neural‑rosette‑like structures are frequently present and are readily transducible. Scale bar: 100 µm. (C) When neural rosettes are isolated, dissociated, and transduced, neurons develop over the following days. Scale bar: 100 µm. (D) Immunostainings confirm neural‑rosette identity by PAX6 expression, while peripheral cells express the neural‑crest marker SOX10. Scale bar: 100 µm. (E) At day 13 after transduction, neurons derived from neural rosettes express TUJ1 but not Peripherin. The combination of PAX6 and TUJ1 expression together with the absence of SOX10 and Peripherin indicates a CNS, most likely telencephalic, neuronal identity. Scale bar: 100 µm.

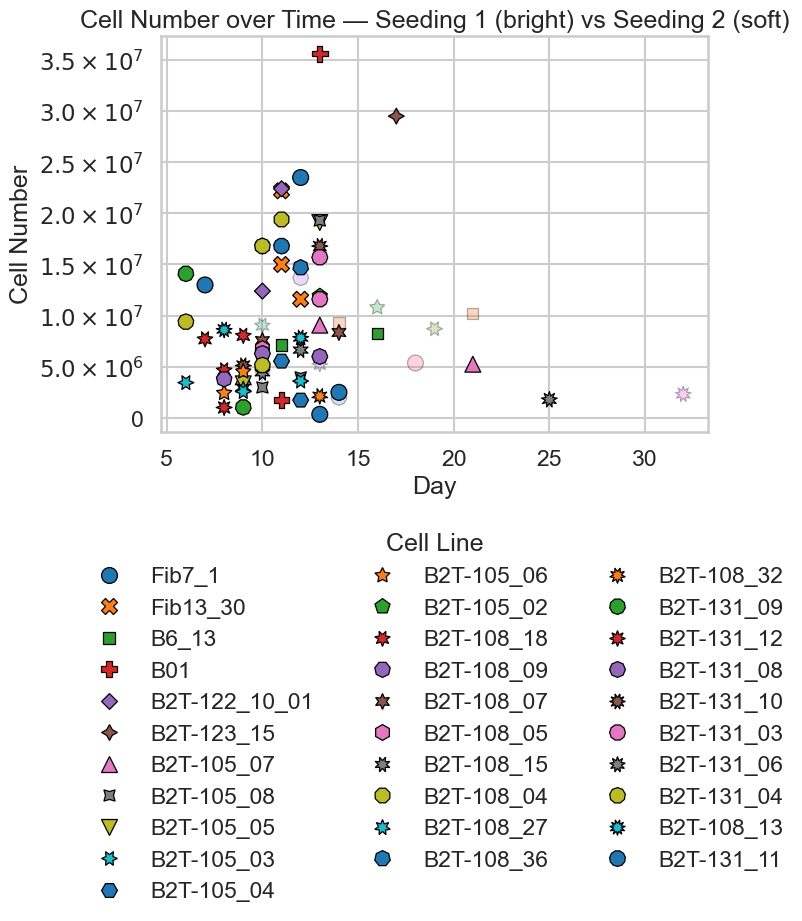

SI Figure 3 Differentiation of NCLCs from multiple cell lines.

Time points (X‑axis) and the number of cryopreserved NCLCs (Y‑axis) are shown. The figure belongs to Figure 2A.

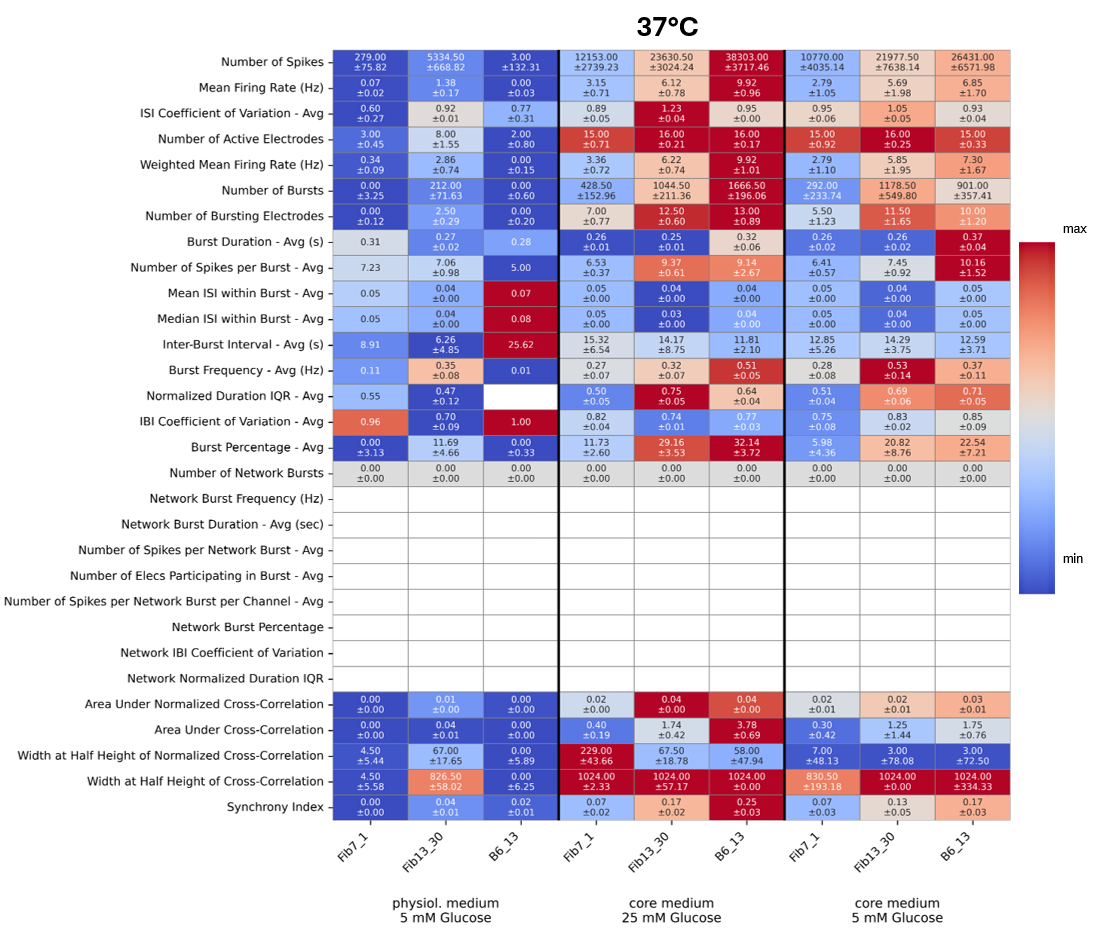

**SI Figure 4** Heatmap of all MEA-Parameters recorded at 37°C and in three different media.

MEA-Recordings of 3 independent cell lines (Fib13_30, B6_13 and Fib7_1) matured in physiological medium, core medium with high [glucose] or low [glucose]. Mean +/-SEM is given per cell and colour code is scaled to maximum value (red) per parameter. White colour indicates no value.

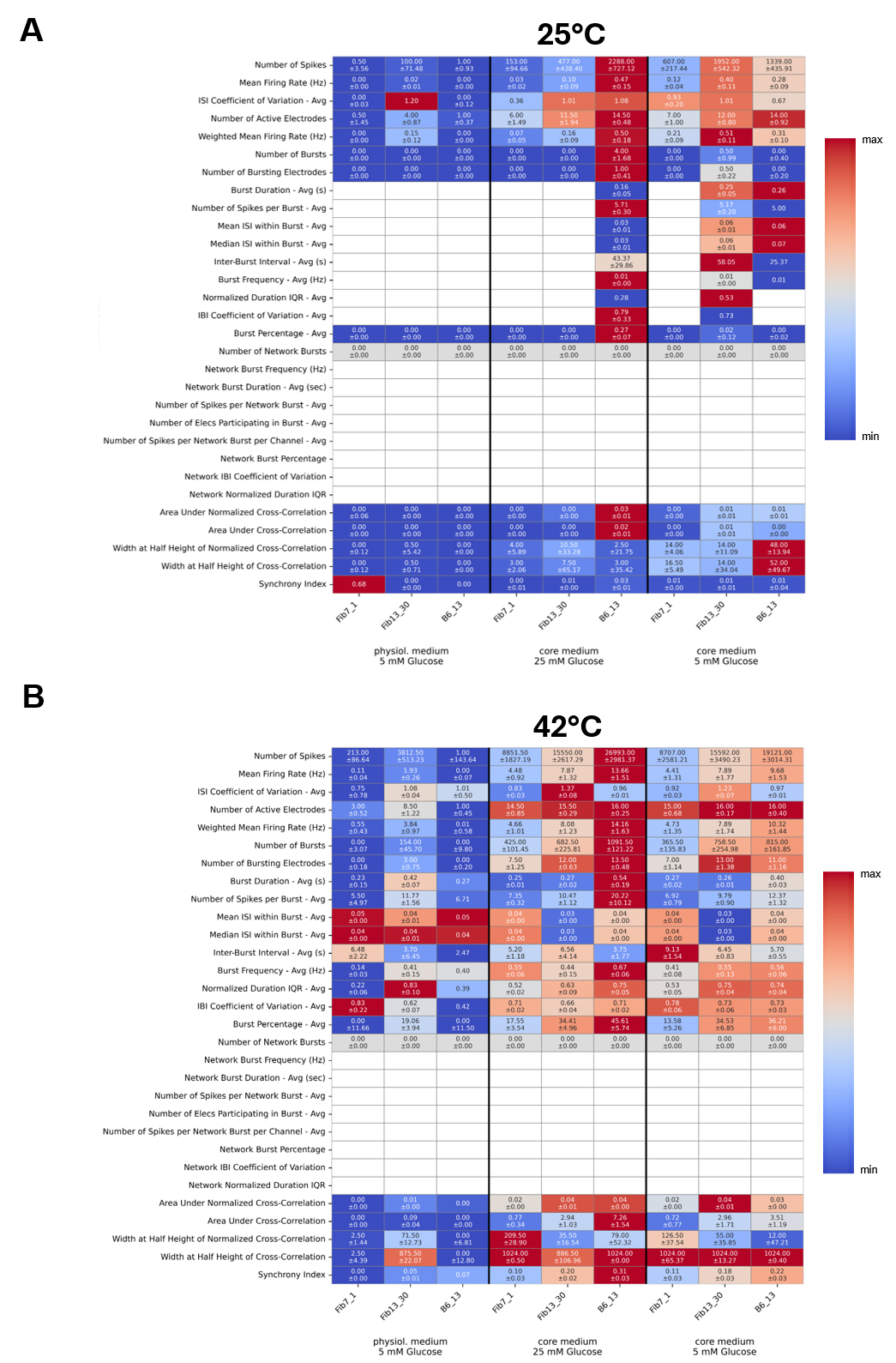

**SI Figure 5 Heatmap of all MEA-Parameters recorded at 25°C and 42°C with 3 different media.**

MEA-Recordings of 3 independent cell lines (Fib13_30, B6_13 and Fib7_1) matured in physiological medium, core medium with high [glucose] or low [glucose]. Mean +/-SEM is given per cell and colour code is scaled to maximum value (red) per parameter. White colour indicates no value.

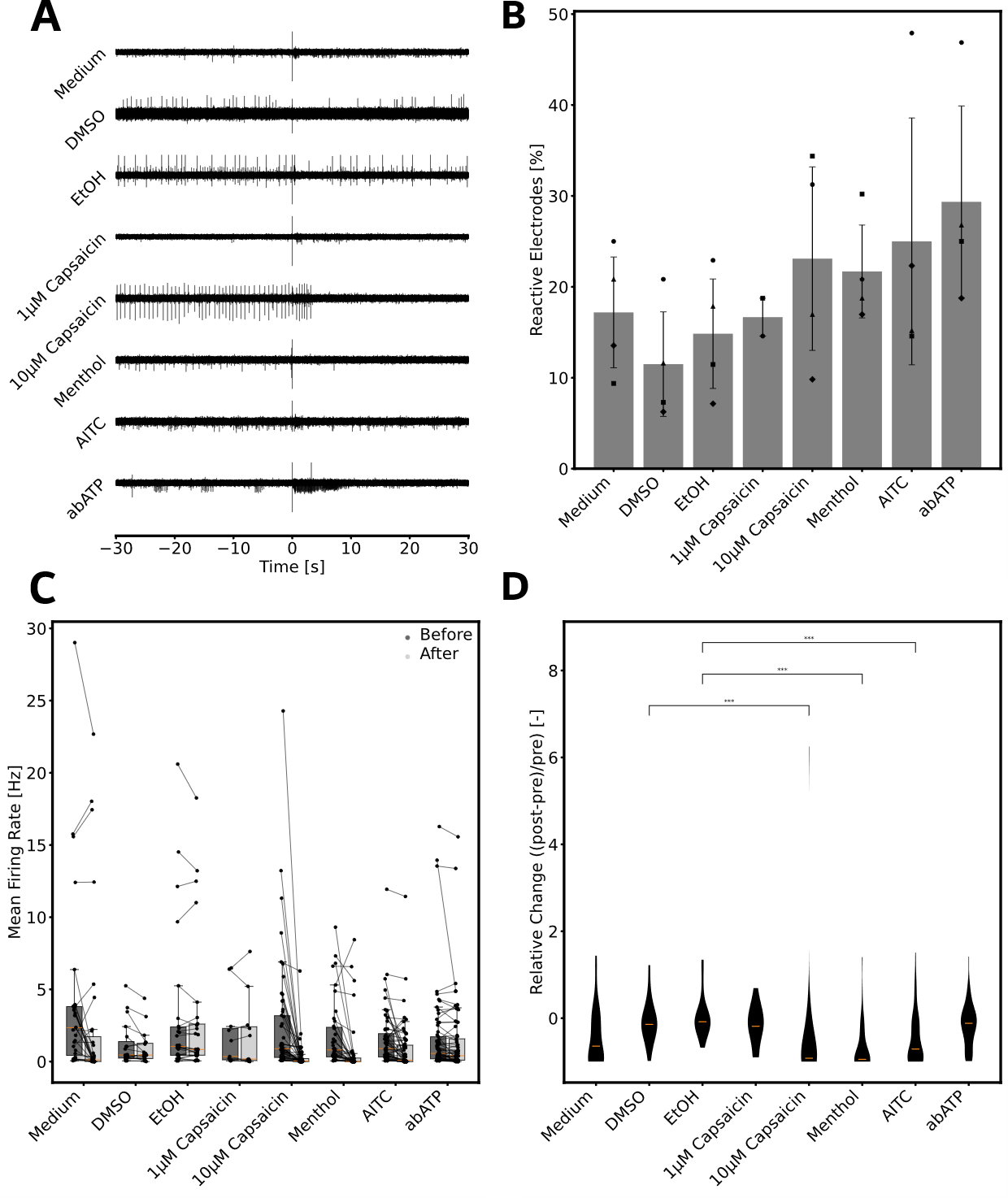

SI Figure 6 Reaction of sensory neurons matured in core medium to ion channel agonists.

(A) Representative traces of cell reaction to compound addition. Traces are centred around the injection time (t=0) with 30 seconds before and after the injection. Traces are individually clipped to [-50, 35] µV before plotting. (B) Electrodes with a firing rate above 0.042 Hz before and after the injection as well as a Jensen-Shannon-Distance greater than 0. Bars indicate mean of all recordings, and error bars indicate standard deviation. (C) Mean firing rate before (dark grey) and after the injection (light grey) as paired samples and boxplots. Each electrode was treated as an independent sample. Medians are indicated by orange lines; whiskers indicate 1^st^ and 3^rd^ quartile ∓ 1.5-time interquartile range respectively. (D) Normalized change in mean firing rate after the compound injection. The change in firing rate was normalized to the firing rate after the injection. Orange bar indicates median. Statistical significance was assessed using a two-sided Mann-Whitney-U test between the compound and its respective solvent control. False discovery rate correction was performed using the Bejamini-Hochberg method (n = 5). p-values are given by * < 0.05, ** < 0.01, *** < 0.001.

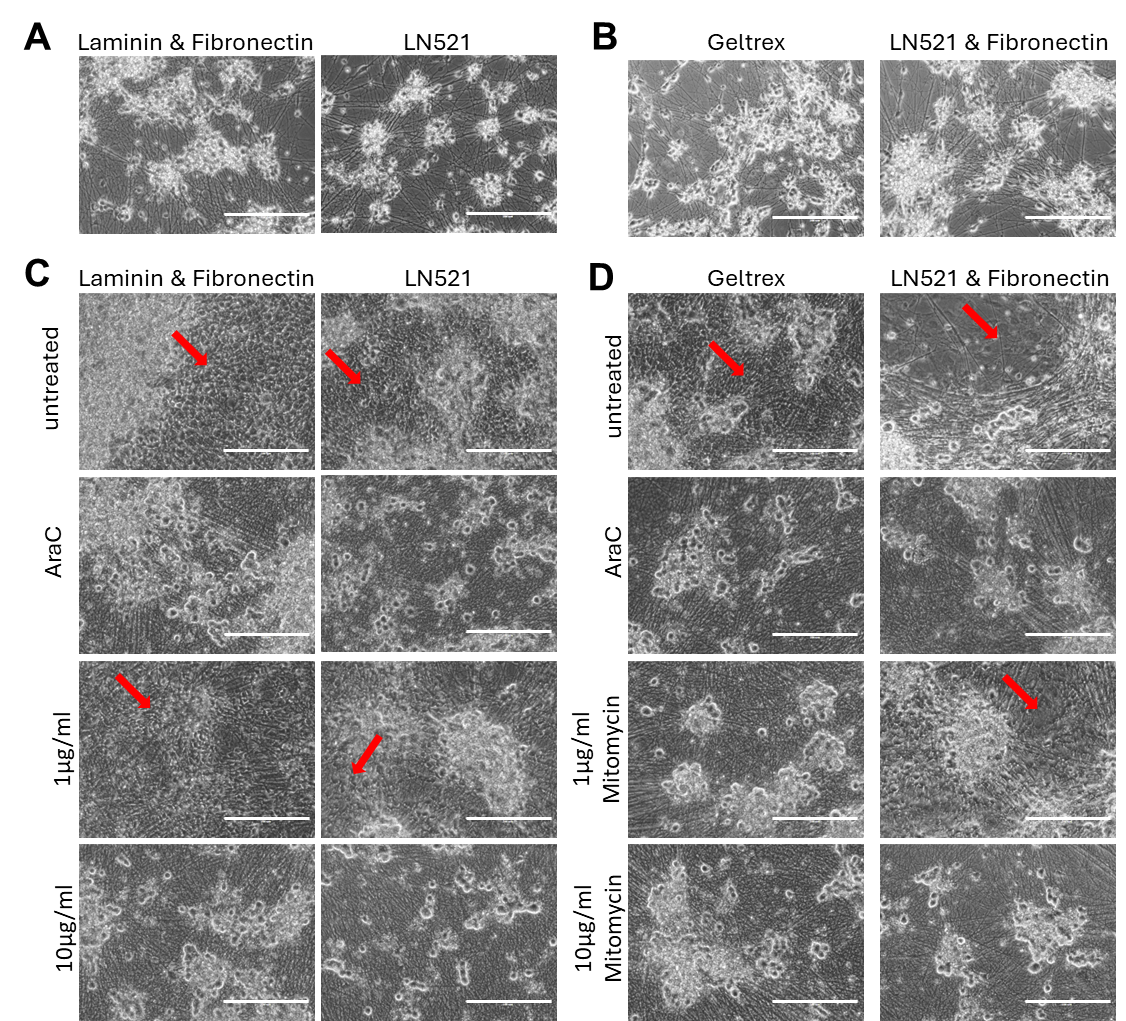

**SI Figure 7 Impact of coating and cytostatics treatment on sensory neuron purity.**

(A, B) Sensory neurons matured on Laminin + Fibronectin, LN521, Geltrex, or LN521 + Fibronectin at day 12 of maturation. At day 13, cultures were treated with cytostatic agents to negatively select contaminating non‑neuronal cells. (C, D) Sensory neuron cultures 29 days after treatment (day 42) with either 2 µM Ara‑C, 1 µg/ml mitomycin for 2 h, or 10 µg/ml mitomycin for 45 min. Contaminating cells are still present after treatment with 1 µg/ml mitomycin for 2 h or in untreated controls (red arrows). Coating material does not influence the presence of non‑neuronal cells or cell attachment throughout differentiation. Scale bar = 200 µm.

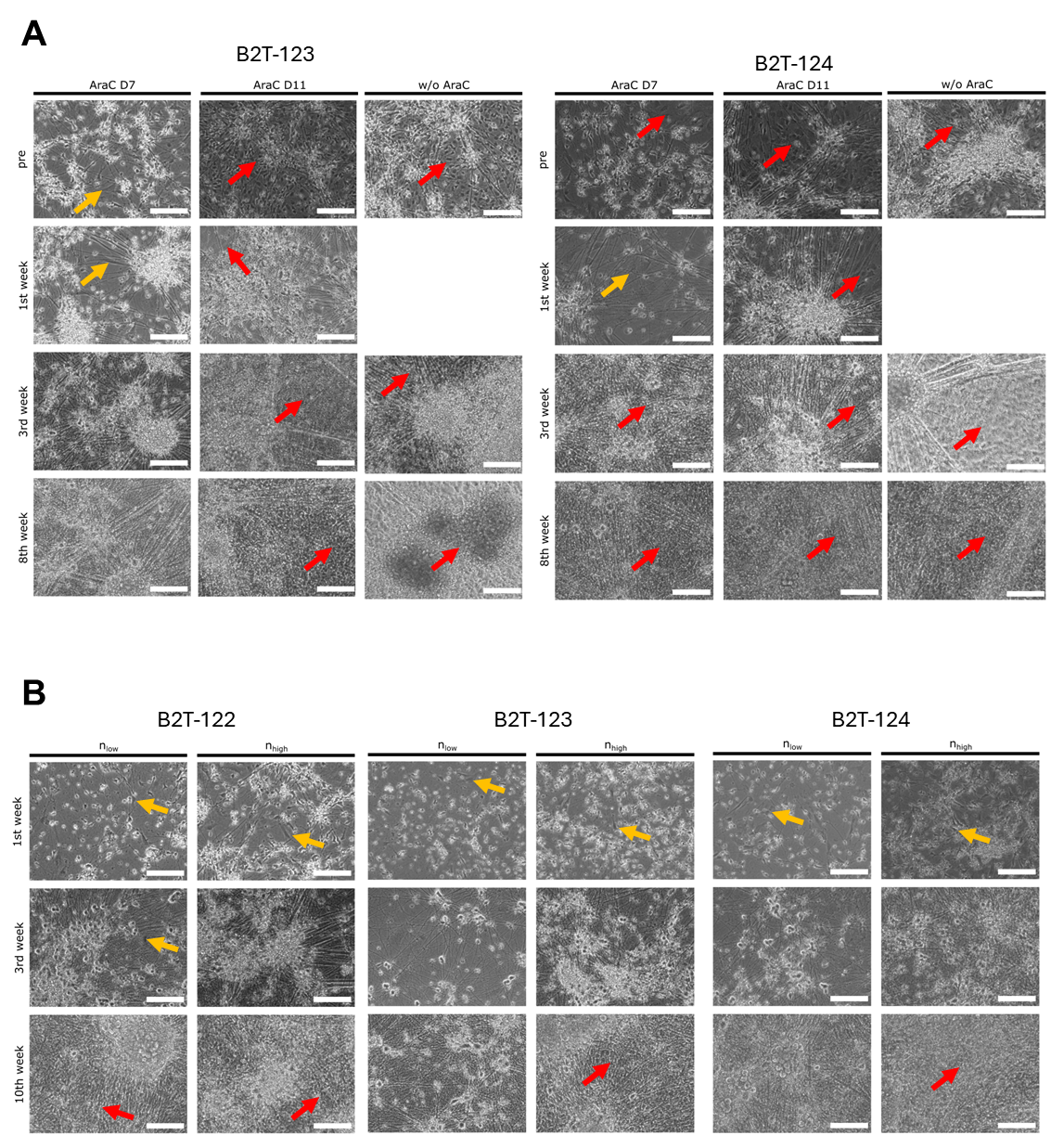

**SI Figure 8 Impact of Ara-C treatment, time point and seeding density on sensory neuron culture purity**

(A) Two independent cell lines treated with Ara‑C at day 7, day 11, or day 14 and subsequent culture development until week 8. (B) Three independent cell lines seeded at high (1,4x10^5^ – 2,1x10^5^ cells/cm²) and low (5,5x10^4^-8,3x10^4^ cells/cm² density at day 5. Ara‑C treatment was performed at day 11. Representative images of neurons at 1 week, 3 weeks, and 10 weeks of maturation are shown. Scale bar = 300 µm. Red arrows indicate strong contamination with non‑neuronal cells; orange arrows indicate single contaminating cells.

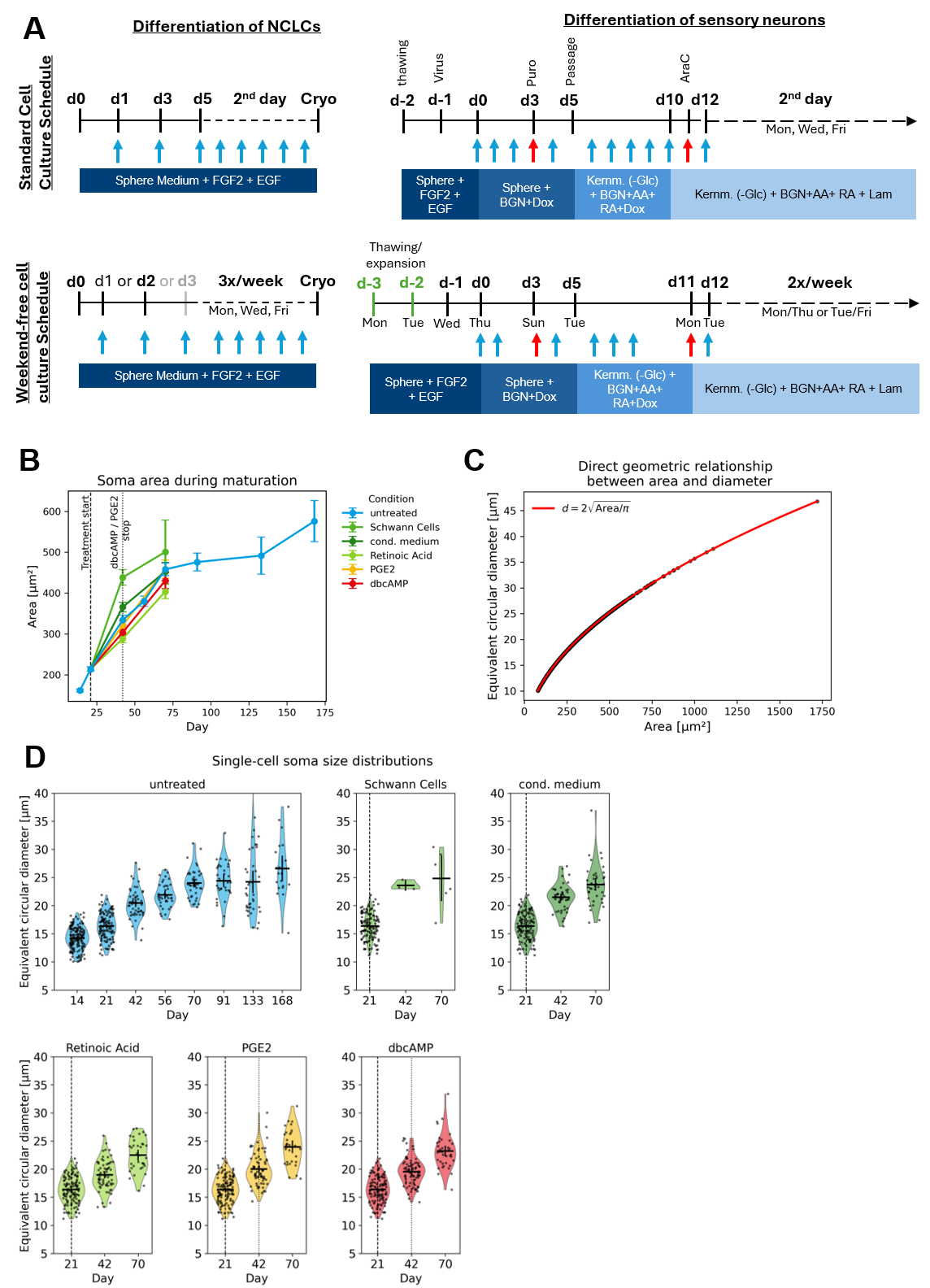

**SI Figure 9 Weekend-free differentiation protocol and development of neurons during maturation**

(A) Differentiation scheme depicting the standard and weekend-free optimised time line. Arrows indicate medium changes and red arrows indicate treatment with selection agents, Puromycin (d3) or Ara-C or Mitomycin, respectively. Abbreviations: d = day, Cryo = Cryopreservation. (B) Soma are measured during maturation of sensory neurons under 6 different conditions. Median +/-SEM is depicted. (C) Direct geometric relationship between area and equivalent circular diameter. For better comparison with previously published data, soma diameter was calculated based on measured soma area. (D) Distribution of the equivalent circular diameter per day of differentiation and maturation condition. Start and stop of treatment is represented as vertical dotted lines.

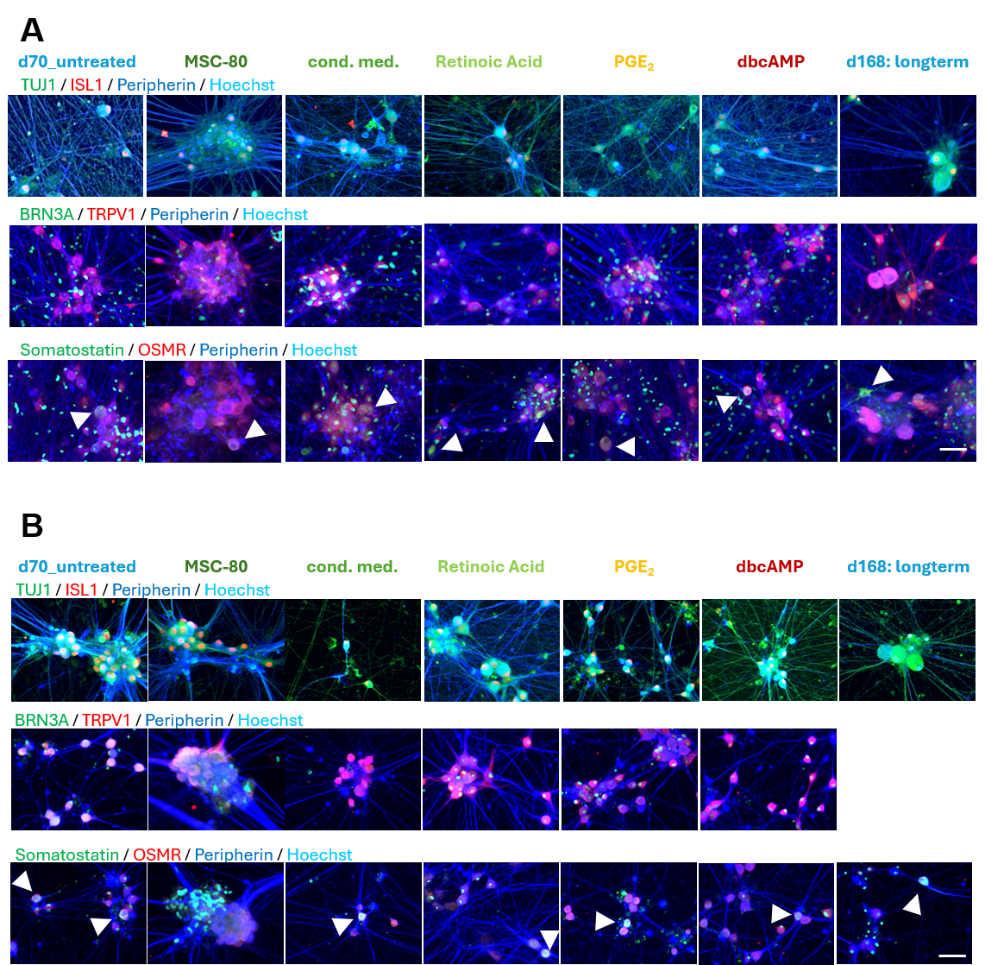

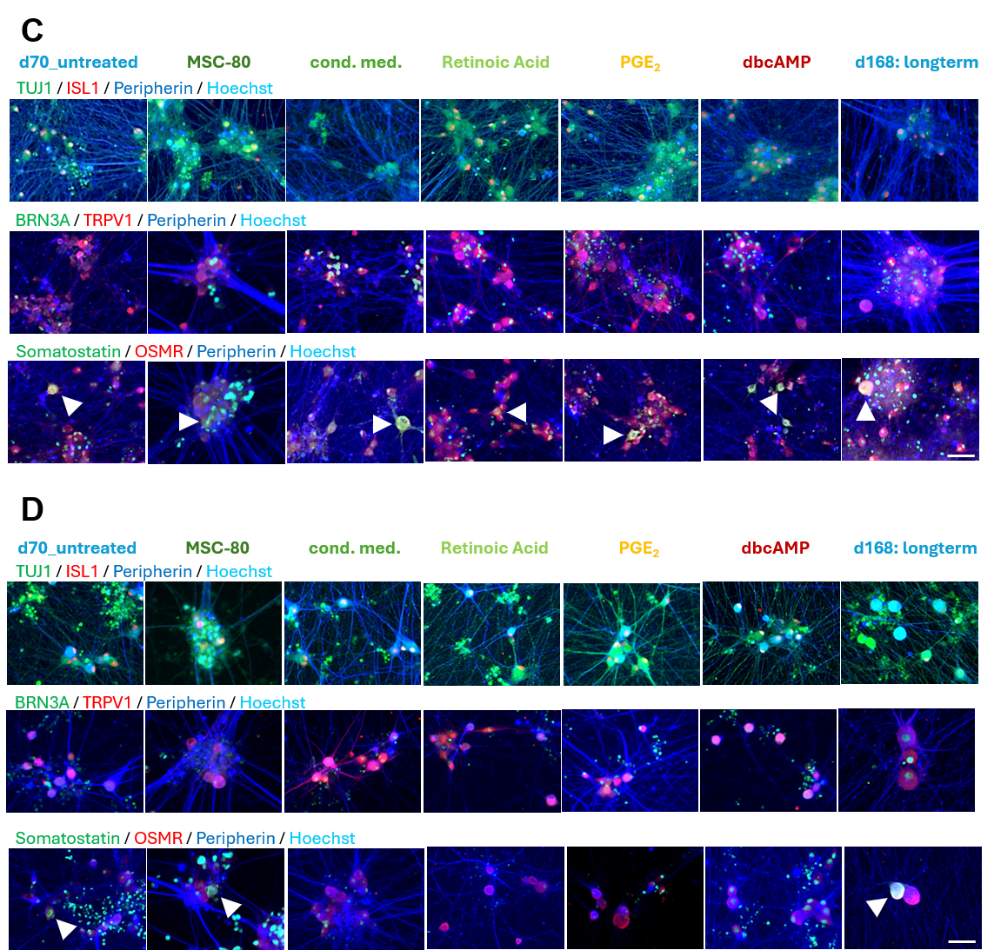

SI Figure 10 Immunostainings of sensory neurons matured under given conditions.

(A/B/C/D) at d70 or d168 neurons of 4 independent cell lines were immunostained for the detection of TUJ1, Peripherin, ISL1, BRN3A, TRPV1, Somatostatin and OSMR. Nuclei are counterstained with Hoechst (Cyan). Somatostatin expressing cells are indicated by arrows. Scale bar = 50µm.

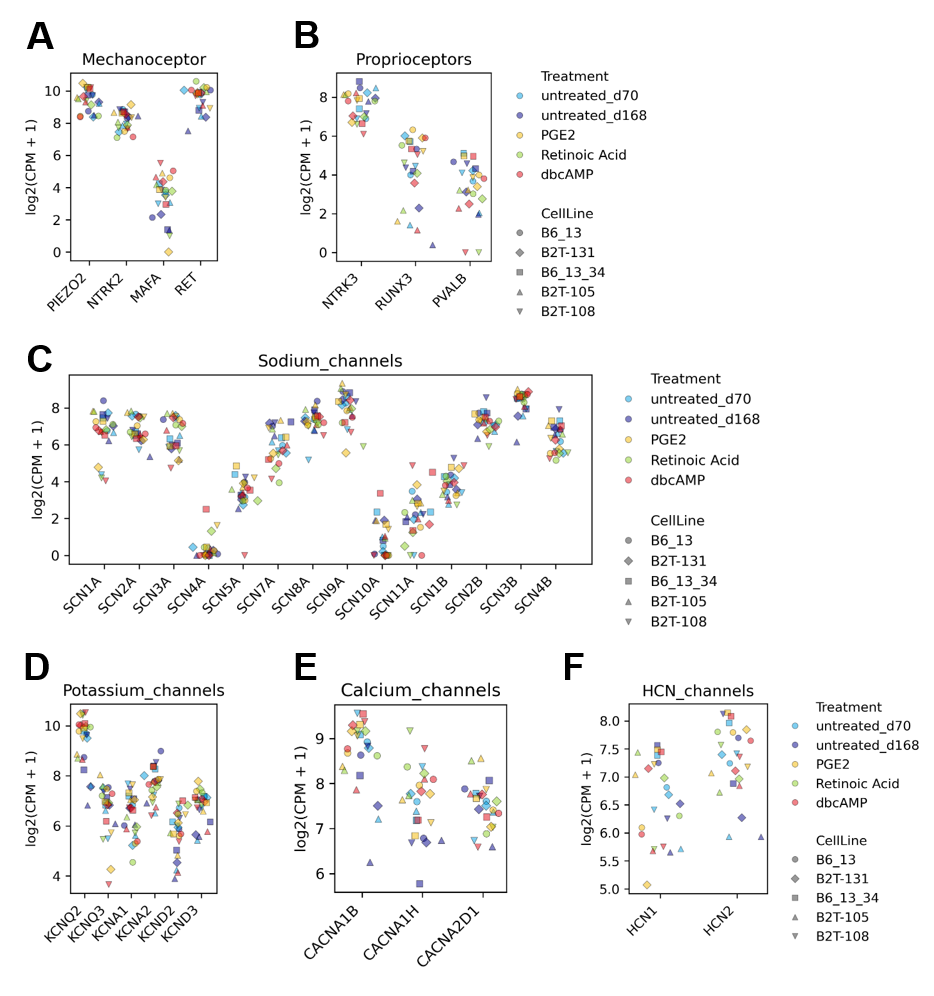

SI Figure 11 mRNA-Marker expression under different sensory neuron maturation condition in five independent cell lines.

(A-F) Expression levels of mechanoceptor, proprioceptor, voltage gated sodium channels alpha and beta subunits, potassium, calcium and HCN channel - associated gene groups across all maturation conditions. Each point represents a sample, colour‑coded by treatment and shaped by cell line.

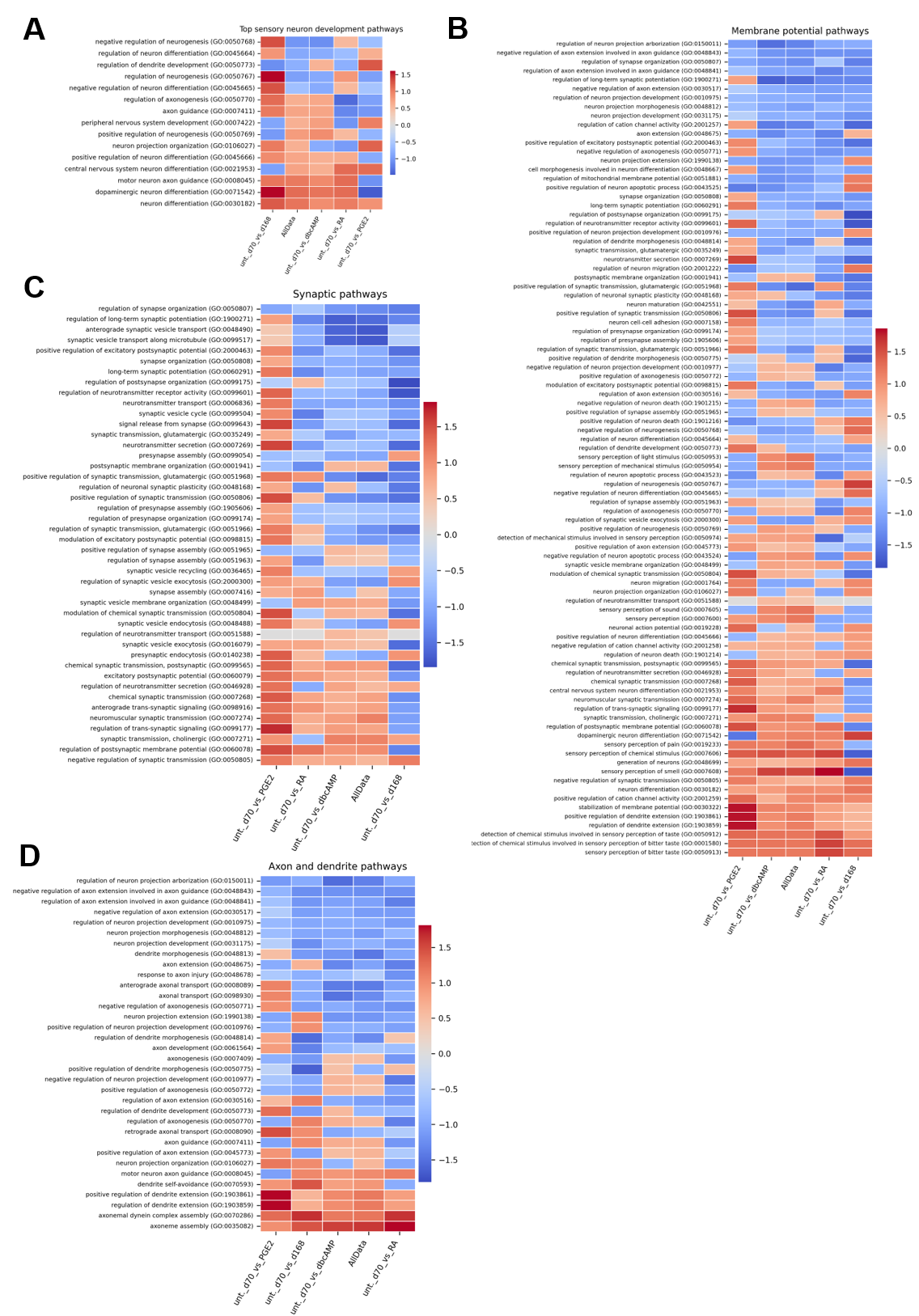

**SI Figure 12 GO‑term enrichment for sensory‑neuron development, membrane‑potential regulation, synaptic and axonal/dendritic pathways across maturation conditions relative to untreated d70 samples.**

(A) GO:BP enrichment of sensory neuron development pathways based on DESeq2 comparisons of each maturation condition versus its matched day 70 control. Pathways related to axon guidance, neuronal morphogenesis, neurogenesis, and neural crest derived lineages show condition‑dependent modulation, with several maturation conditions displaying enrichment relative to day 70. (B) GO:BP enrichment of membrane‑potential–related pathways, including regulation of resting membrane potential, depolarization, ion transport, and voltage‑gated channel activity. Enrichment patterns vary across maturation conditions, reflecting differences in the transcriptional regulation of electrophysiological gene programs. (C) Heatmap showing enrichment of Gene Ontology Biological Process (GO:BP) terms related to synaptic organization and signalling across maturation conditions, calculated relative to matched day 70 controls. Pathways include synapse formation, synaptic transmission, and regulation of synaptic structure. Enrichment patterns vary across conditions. (D) GO:BP enrichment of axonal and dendritic development pathways, including axon guidance, dendrite morphogenesis, and projection regulation. Together, these analyses indicate that transcriptional signatures associated with synaptic and structural neuronal pathways are not strongly modulated by prolonged culture but PGE_2_ increases overall expression.

**SI Table 5 selected genes for comparison with pseudo-bulk RNAseq expression profiles.**

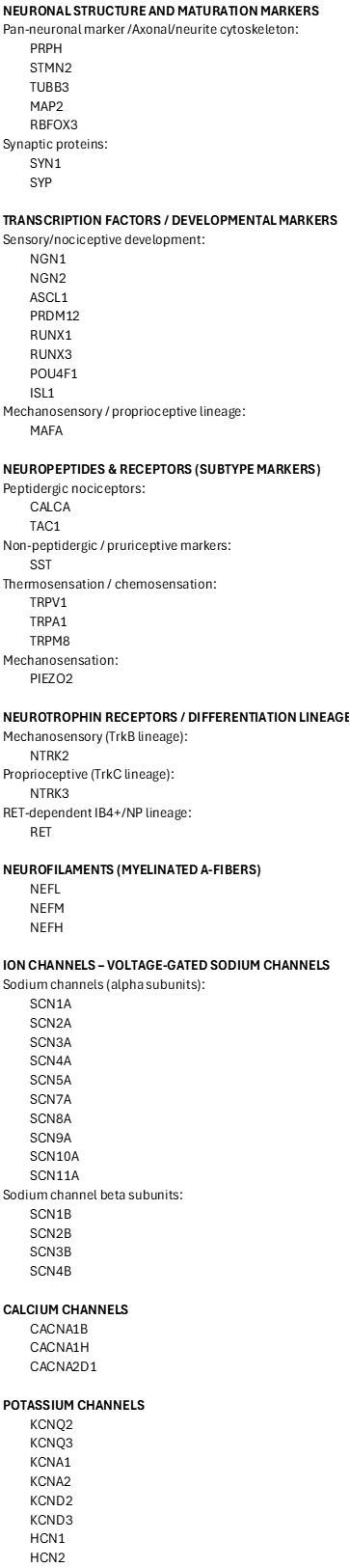

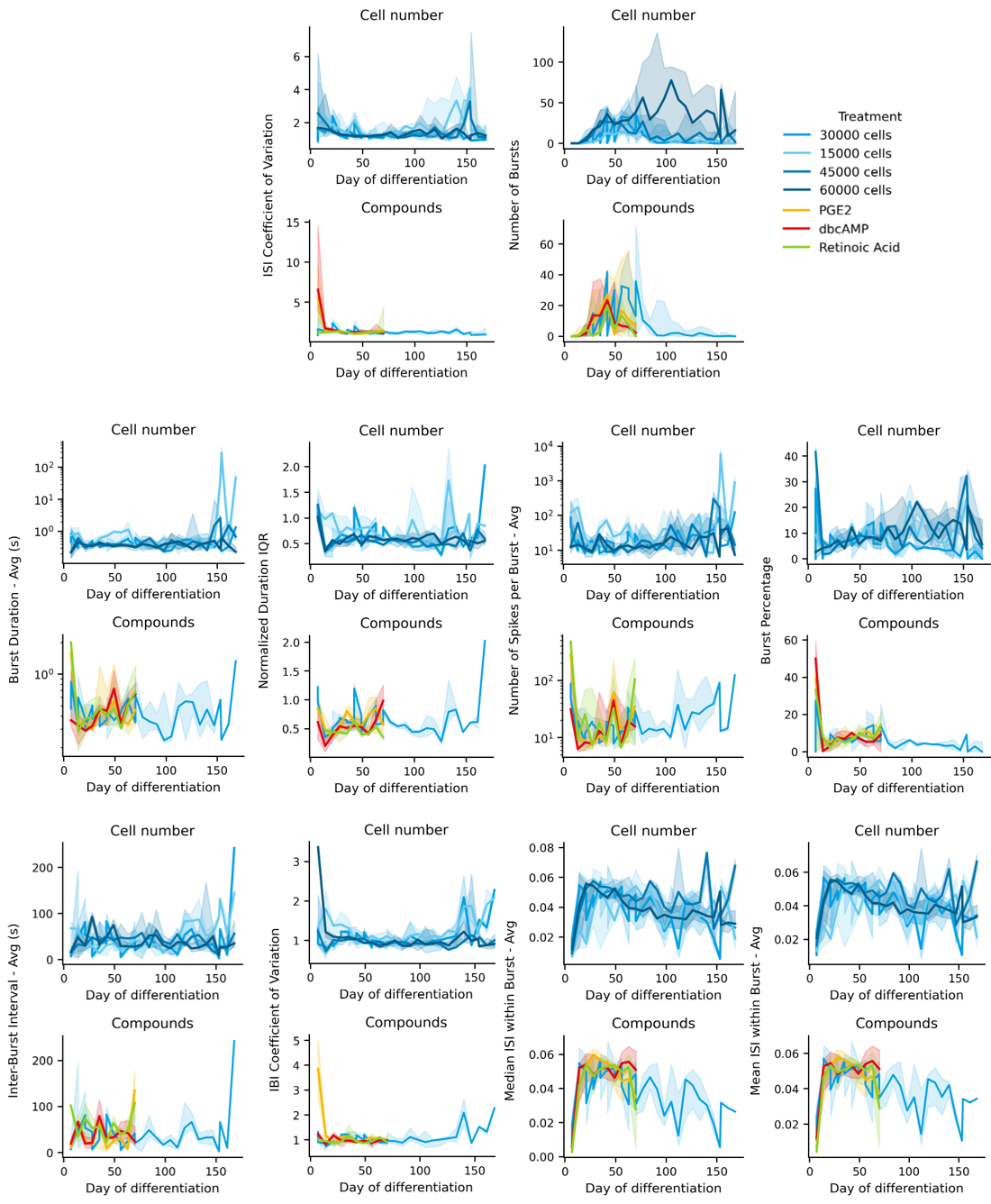

**SI Figure 13 Extended MEA metrics across maturation conditions and cell densities.**

Thirteen electrophysiological parameters were tracked over time across different cell densities (15k–60k) and maturation treatments (PGE_2_, dbcAMP, Retinoic Acid) in five independent cell lines. Time‑course plots of spike‑based and burst‑related metrics including number of spikes, ISI coefficient of variation, number of bursts, burst duration, burst percentage, and spikes per burst. Additional metrics including interburst interval (IBI), IBI variability, and intra‑burst ISI dynamics reveal consistent maturation patterns across conditions. No treatment‑specific functional signature emerged, and most conditions reached peak activity before day 70. Median curves across donors are shown with shaded IQR. Together, these data support the conclusion that functional maturation is primarily driven by time and cell density, with limited impact of chemical maturation cues on MEA‑derived activity profiles

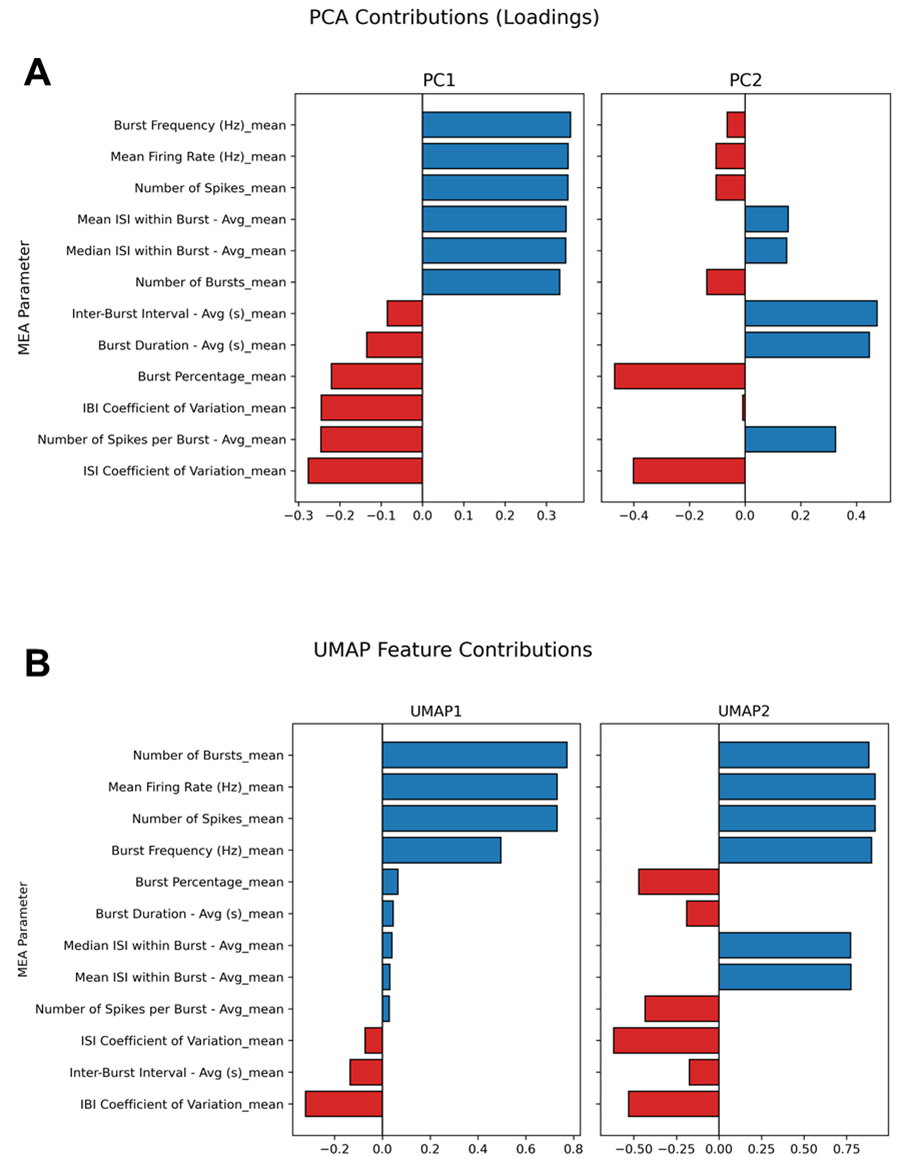

**SI Figure 14 Bar plots showing the contribution of individual MEA parameters to the dimensionality reduction analyses presented in Figure 6.**
(A) PCA loadings for PC1 and PC2 indicate that spike‑ and burst‑related metrics (mean firing rate, number of spikes, burst frequency, number of bursts) dominate the variance captured by PC1, whereas temporal burst structure (inter‑burst interval, burst duration, intra‑burst ISI metrics) contributes more strongly to PC2.
(B) UMAP feature contributions for UMAP1 and UMAP2 reveal a similar pattern, with activity‑related parameters shaping the primary dimension and burst‑structure parameters influencing the secondary dimension.

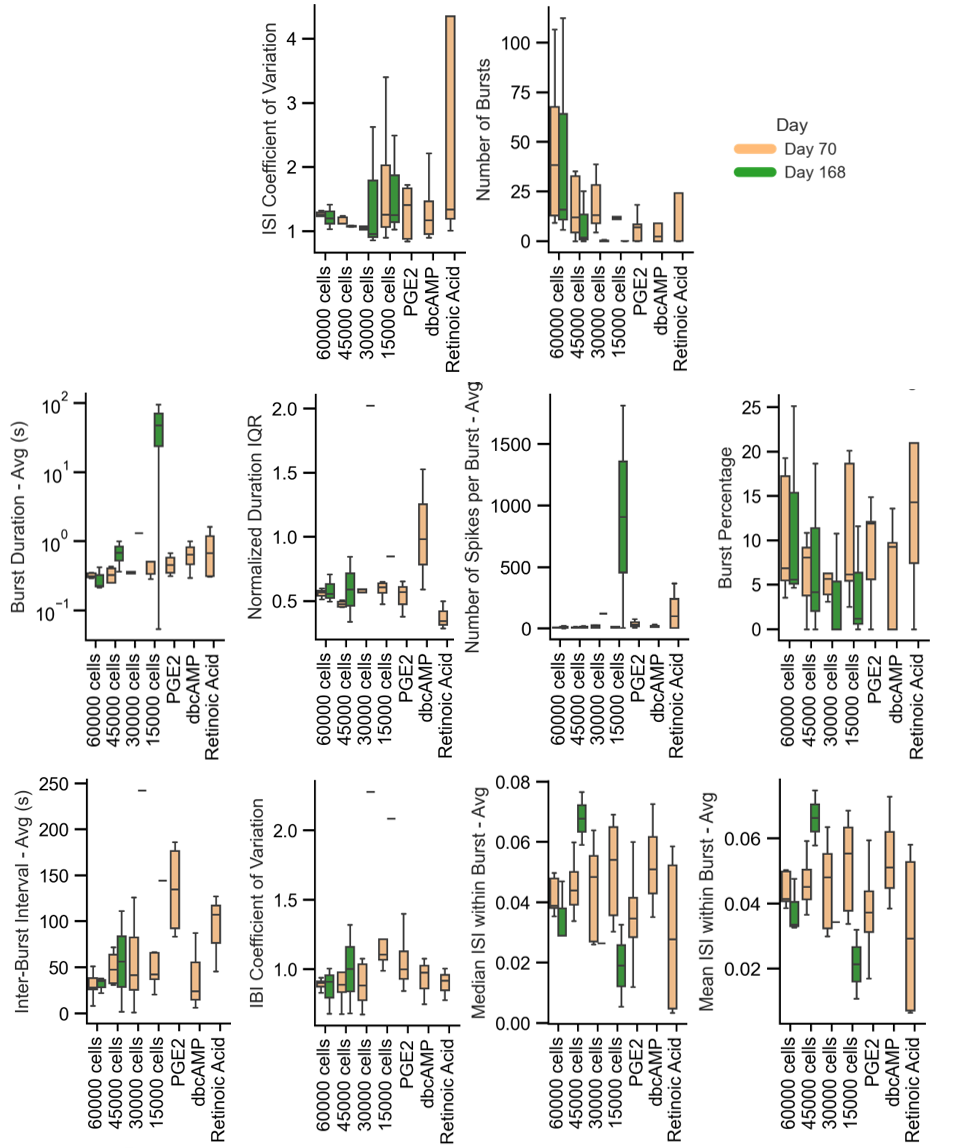

**SI Figure 15 Extended MEA metrics across maturation conditions and cell densities at d70 and d168.**

Box plots comparing eleven electrophysiological parameters across different cell densities (15k–60k) and maturation conditions (±PGE2 dbcAMP, RA) at day 70 (green) and day 168 (orange). Metrics include spike‑based activity (number of spikes, ISI coefficient of variation), burst dynamics (number of bursts, burst duration, burst percentage, spikes per burst), and temporal structure (inter‑burst interval, IBI variability, intra‑burst ISI metrics). Across most conditions, activity parameters increase between day 70 and day 168, with no consistent treatment‑specific enhancement. Supplementation with RA, PGE_2_ or dbcAMP did not yield distinct functional profiles compared to standard conditions, suggesting limited impact of these factors on overall functional maturation.

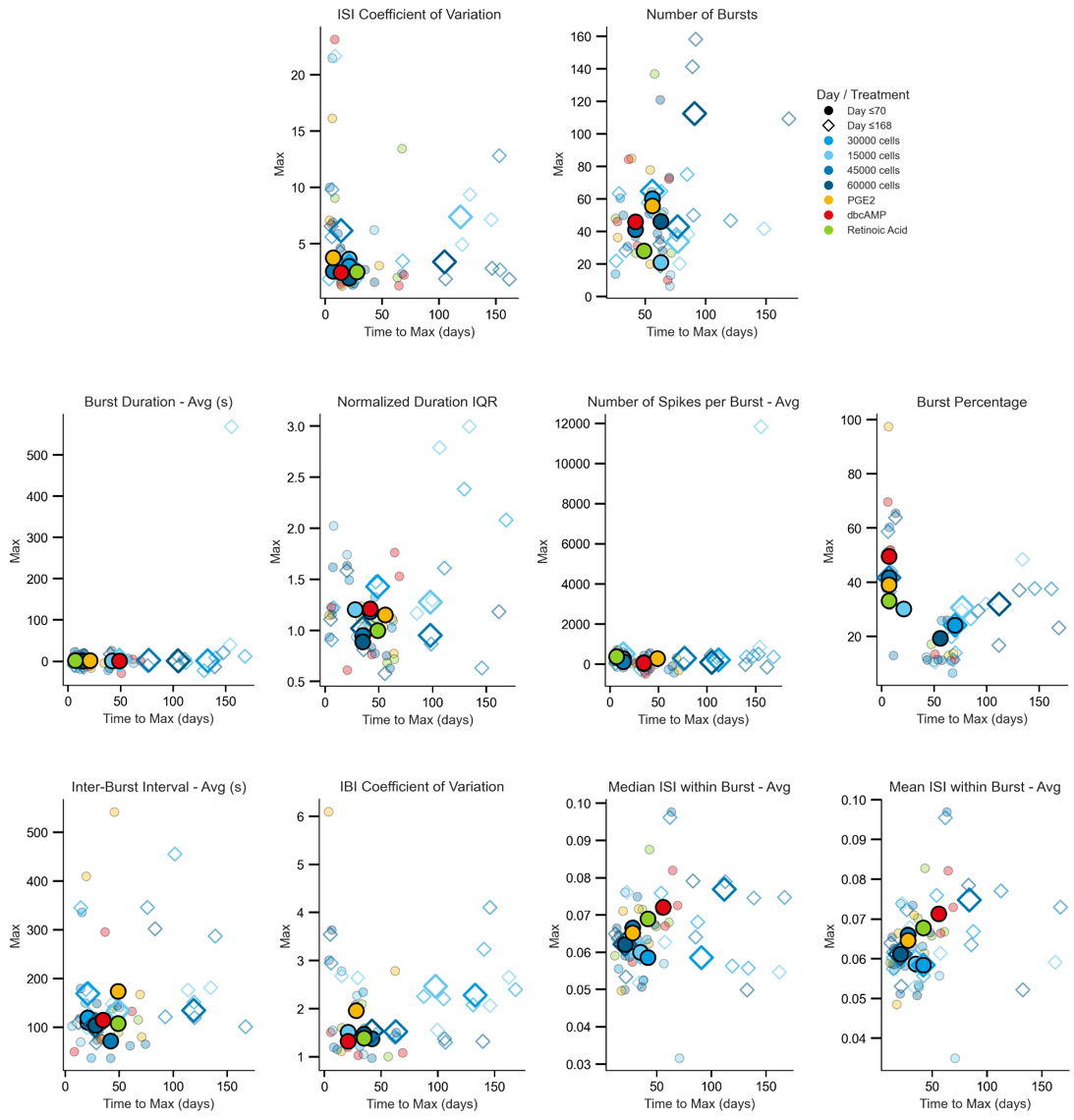

**SI Figure 16 Maximum activity values and time‑to‑maximum across MEA parameters and maturation conditions.**

Scatter plots showing the maximum value (“Max”) and corresponding time‑to‑maximum for all MEA parameters across different cell densities (15k–60k) and maturation treatments (PGE_2_, dbcAMP, RA). Each point represents the median across donors, shape‑coded by maturation stage (≤ day 70, ≤ day 140) and colour coded by treatment condition.
Parameters include spike‑based activity (number of spikes, ISI coefficient of variation), burst dynamics (number of bursts, burst duration, burst percentage, spikes per burst), and temporal structure (inter‑burst interval, IBI variability, intra‑burst ISI metrics). Across all metrics, most conditions reach their maximal activity before day 70, with no treatment‑specific enhancement detectable. These results support the conclusion that functional maturation plateaus early and is largely independent of chemical maturation cues. Bright colours represent the median per treatment and fate colours indicate median per cell line.
